# A neural circuit for wind-guided olfactory navigation

**DOI:** 10.1101/2021.04.21.440842

**Authors:** Andrew M.M. Matheson, Aaron J. Lanz, Ashley M. Medina, Al M. Licata, Timothy A. Currier, Mubarak H. Syed, Katherine I. Nagel

**Affiliations:** Neuroscience Institute, NYU Medical Center, 435 E 30th St. New York, NY 10016, USA; Center for Neural Science, NYU, New York, NY, 4 Washington Place, New York, NY 10003; Department of Biology, 219 Yale Blvd NE, University of New Mexico, Albuquerque, NM 87131

## Abstract

To navigate towards a food source, animals must frequently combine odor cues that tell them what sources are useful with wind direction cues that tell them where the source can be found. Where and how these two cues are integrated to support navigation is unclear. Here we identify a pathway to the *Drosophila* fan-shaped body (FB) that encodes attractive odor and promotes upwind navigation. We show that neurons throughout this pathway encode odor, but not wind direction. Using connectomics, we identify FB local neurons called hΔC that receive input from this odor pathway and a previously described wind pathway. We show that hΔC neurons exhibit odor-gated, wind direction-tuned activity, that sparse activation of hΔC neurons promotes navigation in a reproducible direction, and that hΔC activity is required for persistent upwind orientation during odor. Based on connectome data, we develop a computational model showing how hΔC activity can promote navigation towards a goal such as an upwind odor source. Our results suggest that odor and wind cues are processed by separate pathways and integrated within the FB to support goal-directed navigation.

## Introduction

Searching for a resource such as food requires integration of multiple sensory cues. In natural environments, food odors are often transported by wind, forming turbulent plumes (Murlis et al. 1992, Cardé and Willis, 2008). Within these plumes, instantaneous odor concentration is a poor cue to source direction (Crimaldi and Kossef 2001, Webster and Weissburg 2001, Celani et al. 2014). Thus, many organisms have evolved a strategy of using odor information to gate upwind (or upstream) movement to locate the source of an attractive odor (David et al. 1983, Wolf and Wehner, 2000, Page et al. 2011, van Breugel et al. 2014, Alvarez-Salvado et al. 2018). This strategy complements those observed in odor gradients, where odor increases drive straighter trajectories, while odor decreases drive re-orientation (e.g. Schulze et al. 2015). Navigation towards potential food sources thus requires integration of directional information about the prevailing wind (often derived from mechanosensation), with information about the identity and quality of odors carried on that wind (Alvarez-Salvado et al. 2018, van Breugel et al. 2014). Where and how these two types of information are integrated to support navigation towards an odor source is not clear.

In the insect brain, several conserved central neuropils have been implicated in olfactory food search and navigation (Fig. 1A). The mushroom body (MB) and lateral horn (LH) have been implicated in learned and innate olfactory processing respectively (deBelle and Heisenberg, 1994, Hige et al. 2015b, Schlegel et al. 2021). Subsets of MB and LH output neurons (MBONs and LHONs) promote approach and avoidance behavior (Aso et al. 2014, Owald et al. 2015, Dolan et al. 2019, Huoviala et al. 2020). A number of putative mechanosensory inputs to the MB have been identified (Bates et al. 2020) and wind intensity signals have been observed in certain MB compartments (Mamiya et al. 2008). The LH also receives input from mechanosensory centers in a discrete ventral region (Dolan et al. 2019, Bates et al. 2020). However, it is not known whether the MB and LH represent wind direction signals, as well as odor identity and value signals, to support navigation.

**Fig. 1:**
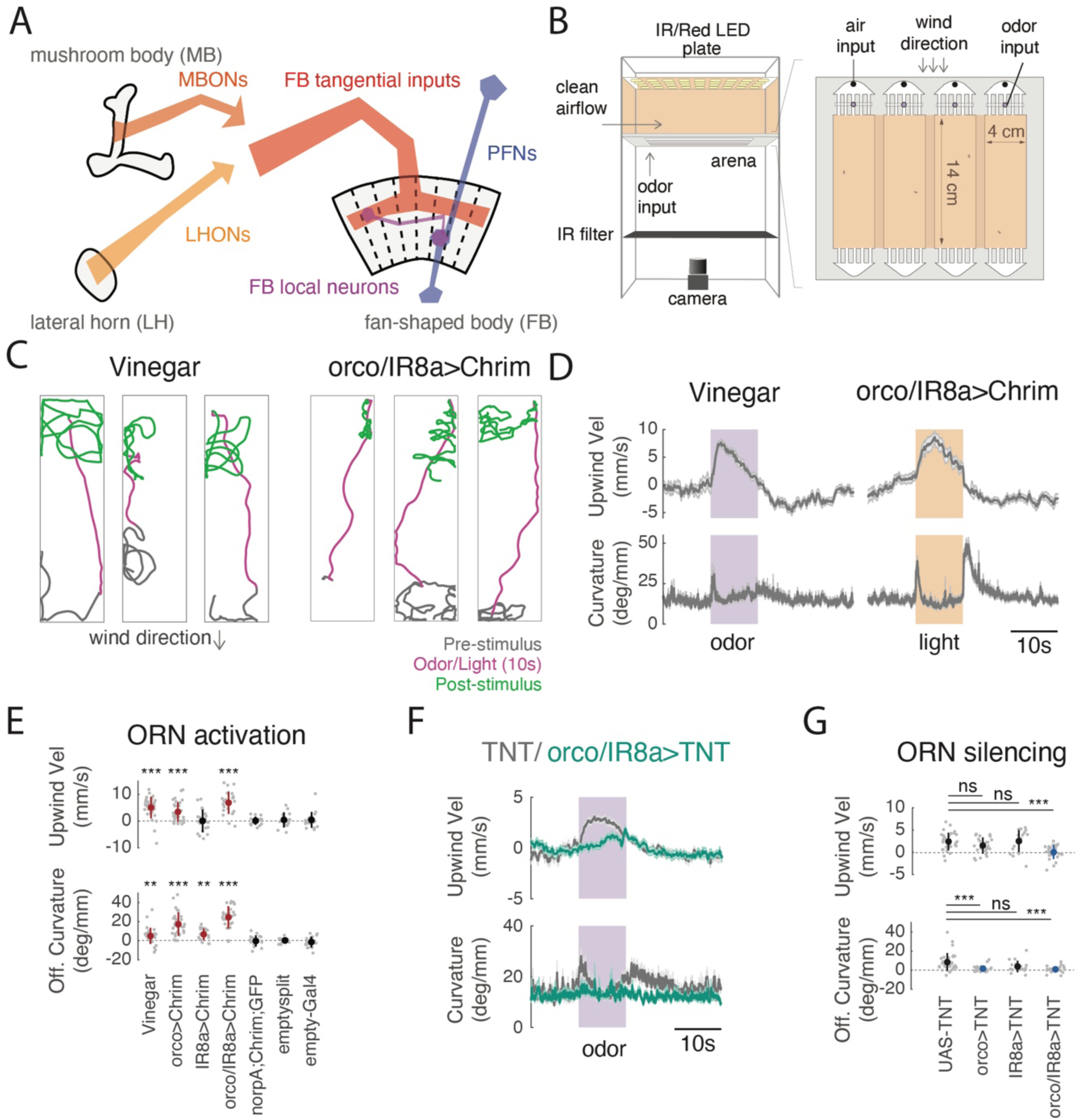
A behavioral paradigm for investigating odor-evoked wind navigation. **A)** Diagram of brain regions and neuronal classes investigated in this study. The mushroom body (MB) is required for olfactory learning while the lateral horn (LH) is thought to perform innate olfactory processing. The fan-shaped body (FB) is part of the central complex, and is thought to play a role in spatially-oriented navigation. Output neurons of the MB and LH (MBONs and LHONs) provide input directly or indirectly to FB tangential inputs (Li et al. 2020, Scheffer et al. 2020, Hulse et al. 2021). The FB also receives input from columnar cells (PFNs) that have been shown to strongly encode wind direction information (Currier et al. 2020). FB local neurons receive input both from FB tangential inputs and from columnar PFNs (Hulse et al. 2021). **B)** Schematic of top and side view of the behavioral apparatus showing IR illumination (850nm), red activation light (626nm, 26µW/mm^2^), imaging camera, behavioral chambers, and air and odor inputs. **C)** Navigation behaviors evoked by odor and optogenetic stimulation of olfactory receptor neurons. Example walking trajectories in response to apple cider vinegar (left, vinegar, 1%) and optogenetic activation of orco+ and IR8a+ ORNs (right), before (gray), during (magenta), and after (green) 10s of odor (left) or light (right). Wind was constant at 11.9 cm/s. **D)** Time course of upwind velocity and curvature (angular/forward velocity) in response to odor or optogenetic stimulation, averaged across flies (mean±SEM, vinegar N= 26 flies, ORN activation N= 24 flies). shaded area: stimulation period (10s 1% vinegar (purple) or 10s light (orange) respectively). **E)** Upwind velocity and OFF curvature (average change from baseline for single flies) in response to stimulation for each genotype/condition. Mean±STD overlaid; red indicates a significant increase. Vinegar, orco>Chrimson, and orco/IR8a>Chrimson stimulation all drove significant increases in upwind velocity (0-5 s after stimulus ON, Wilcoxon signed rank test, vinegar: p= 1.0997e-05, orco: p=3.8672e-05, orco,ir8a: 2.6691e-05) and OFF curvature (0-2 s after stimulus OFF, Wilcoxon signed rank test, vinegar: p=1.9619e-04, orco: p=1.1742e-06, orco,ir8a: p=2.3518e-05). Light activation of flies carrying only the parental effector (norpA;UAS-Chrimson), empty-GAL4>Chrimson, or empty-split GAL4>Chrimson did not increase upwind velocity (Wilcoxon signed rank test, parent: p= 0.7174, empty-split: p=0.6874, empty-gal4: p=0.6698) or OFF curvature (Wilcoxon signed rank test, parent: p= 0.7960, empty-split: p=0.3144 ,empty gal4: p=0.7354). IR8a>Chrimson stimulation did not increase upwind velocity (p=0.3507) but did increase OFF curvature (p=4.4934e-04). **F)** Time course of upwind velocity and curvature in response to odor in flies with all vinegar-sensitive ORNs silenced (orco/IR8a>TNT, mean±SEM, N=26, teal) versus control (UAS-TNT, N= 31, gray). **G)** Upwind velocity and OFF curvature for silencing experiments, quantified as in **E**. Blue overlay represents significant decrease compared to control (UAS-TNT). (Mann-Whitney U test compared to UAS-TNT control, upwind velocity: orco: p= 0.10627, ir8a: p=0.40095, orco,ir8a: p=0.00010917, OFF curvature: orco: p=3.2758e-05, ir8a: p=0.037135, orco,ir8a: p=4.2482e-05) All comparisons were corrected using the Bonferroni method (see Methods).

In contrast, the fan-shaped body (FB), a part of the *Drosophila* navigation center called the central complex (CX), has been recently shown to encode wind direction (Currier et al. 2020). Columnar inputs to the fan-shaped body (FB), known as PFNs, represent wind direction as a set of orthogonal basis vectors, and receive input from the lateral accessory lobe (LAL) via LNa neurons (Currier et al. 2020). Wind direction signals are strongest in PFNs targeting ventral layers of the FB (PFNa, p, and m in the hemibrain connectome, Scheffer et al. 2020). PFNs targeting more dorsal layers (PFNd and v) encode both optic flow (Lyu et al. 2020) and self-motion during walking (Lu et al. 2020) in a similar vector format. PFNs of all types show little sensitivity to odor stimuli (Currier et al. 2020), suggesting that this pathway mostly encodes flow and self-motion information independent of odor.

A distinct set of FB inputs, known as FB tangential cells, are anatomically downstream of the MB (Li et al. 2020, Scaplen et al. 2020), however most of these have not been functionally characterized. To date, few olfactory inputs to the FB have been described (Homberg, 1985). Large lesions to the FB disrupt visual navigation (Strauss and Heisenberg 1993), and activation of subsets of FB neurons can produce oriented locomotor behaviors in cockroaches (Martin et al. 2015). Recent theoretical and experimental work suggests that FB circuitry is optimized for encoding vectors and specifying navigational goals (Hulse et al. 2020, Goulard et al. 2021, Lyu et al. 2021, Lu et al. 2021), but experimental evidence for goal encoding in the FB is still sparse. Numerous studies have explored the role of the FB in path integration (Stone et al. 2017, Hulse et al. 2021), visual navigation (Sun et al. 2020), and landmark-guided long-distance dispersal (Dacke et al. 2019, Honkanen et al. 2019, Leitch et al. 2021). However, few studies have investigated the role of this region in olfactory navigation (Sun et al. 2021).

Here we used an optogenetic wind-navigation paradigm, together with calcium imaging, connectomic analysis, and computational modeling to ask how the MB/LH and FB work together to promote olfactory navigation behavior. We show that a subset of attraction-promoting MB and LH neurons evoke upwind movement when activated. However, calcium imaging indicates that these neurons do not strongly encode wind direction, suggesting that integration of odor and wind information occurs elsewhere. We next performed a large behavioral screen of FB inputs, finding that several groups of FB tangential inputs, but not columnar PFNs, drive upwind movement. Imaging revealed that these neurons also encode odor but not wind direction. Finally, we identify a specific type of FB local neuron, called hΔC, that receives input from both wind-sensitive PFNs and from odor-sensitive FB tangential cells. We show that these neurons encode an odor-gated wind direction-tuned signal, and promote navigation in a reproducible direction when sparsely activated in a fly-specific pattern. Silencing hΔC neurons impairs the ability of flies to maintain upwind orientation throughout an odor stimulus. Based on motifs from the fly connectome, we develop a computational model showing how different patterns of activity in hΔC neurons can promote navigation either upwind (under natural odor and wind activation) or in a reproducible arbitrary direction (during sparse optogenetic activation). Taken together, our data support an emerging model of the FB, in which spatial direction cues and non-spatial context cues enter this region through distinct anatomical pathways, and are integrated by local neurons to specify goal-directed navigation behaviors.

## Results

### An optogenetic paradigm to investigate the neural circuit basis of upwind orientation

To investigate the neural circuit basis of wind-guided olfactory navigation, we developed an optogenetic activation paradigm. We modified a set of miniature wind tunnels (Fig. 1B, Alvarez-Salvado et al. 2018) to present temporally-controlled red light stimuli as walking flies navigated in a laminar wind flow. To validate this assay, we asked whether optogenetic activation of olfactory receptor neurons (ORNs) with Chrimson could produce behavioral phenotypes similar to those observed with an attractive odor, apple cider vinegar (vinegar). We found that broad activation of olfactory receptor neurons using either the orco, or the orco and IR8a co-receptor promoters together (Larsson et al 2004, Silbering et al. 2011) resulted in robust navigation behaviors similar to those observed with vinegar (Fig. 1C,D, Fig. S1A). In response to either odor or light, flies ran upwind, generating an increase in upwind velocity. Following odor or light OFF, flies initiated a local search, characterized by increased curvature (Fig. 1C,D). Neither behavior was observed in the absence of the orco-GAL4 or orco/IR8a-GAL4 driver, or when we expressed Chrimson under an empty-GAL4 or empty split-GAL4 driver (Fig. 1E, Fig. S1B). Silencing both orco and IR8a-positive ORNs using tetanus toxin abolished both upwind and search responses to odor (Fig. 1F, G). Thus, optogenetic activation can substitute for odor in producing both upwind orientation and OFF search, and ORNs are required for these behavioral responses to odor.

Vinegar activates a subset of both orco+ and IR8a+ glomeruli (Jung et al, 2015). Although the behavioral phenotypes evoked by vinegar and by optogenetic activation of ORNs were similar, they exhibited some subtle differences. Vinegar produced a stronger upwind response than optogenetic activation of orco+ ORNs in the same flies (Fig. S1A). However, the OFF search behavior evoked by optogenetic activation orco-GAL4, or of orco/IR8a-GAL4, was more robust than that evoked by vinegar (Fig. 1C,D, Fig. S1A). Moreover, activation of orco/IR8a+ ORNs in the absence of wind produced OFF search without upwind orientation (Fig. S1C). These results indicate that upwind orientation can be evoked independently of OFF search, and suggest that these two behaviors are driven by distinct but overlapping populations of olfactory glomeruli. Activation of single ORN types known to be activated by vinegar (Jung et al, 2015), did not generate significant upwind orientation or OFF search (Fig. S1D). In addition, silencing of orco+ or IR8a+ ORNs alone did not abolish upwind orientation, but did reduce OFF search (Fig. 1G, Fig. S1E). These data indicate that groups of ORNs must be activated together to promote upwind orientation and that substantial silencing of most olfactory neurons is required to abolish upwind movement in response to vinegar.

### A subset of LHONs and MBONs drive wind orientation and encode a non-directional odor signal

We next asked whether activation of central neurons in the LH and MB could similarly produce wind navigation phenotypes. We activated several groups of LHONs and MBONs that were previously shown to produce attraction or aversion in quadrant preference assays (Aso et al. 2014, Dolan et al. 2019). We found that several of these neuron groups drove robust upwind movement when activated (Fig 2A,B, Fig. S2A). Neurons promoting upwind movement included the cholinergic LHON cluster AD1b2 (labeled by LH1396, LH1538, and LH1539 (Fig. 2A), and the cholinergic MBON lines MB052B (labeling MBONs 15-19), MB077B (labeling MBON12), and MB082C (labeling MBONs 13 and 14, Fig. 2B). AD1b2 drivers and MB052B also elicited significant increases in OFF curvature when activated (Fig. 2A,B), while activating individual MBONs within MB052B (MBONs 15-19) did not drive significant upwind movement (Fig. S2B). Silencing single MBON or LHON lines that drove navigation phenotypes did not abolish upwind movement in response to odor (Fig. S2C) consistent with models suggesting that odor valence is encoded by population output at the level of the MB (Owald and Waddell, 2015), and with our findings at the periphery that very broad silencing is required to eliminate behavioral responses to vinegar.

**Fig. 2:**
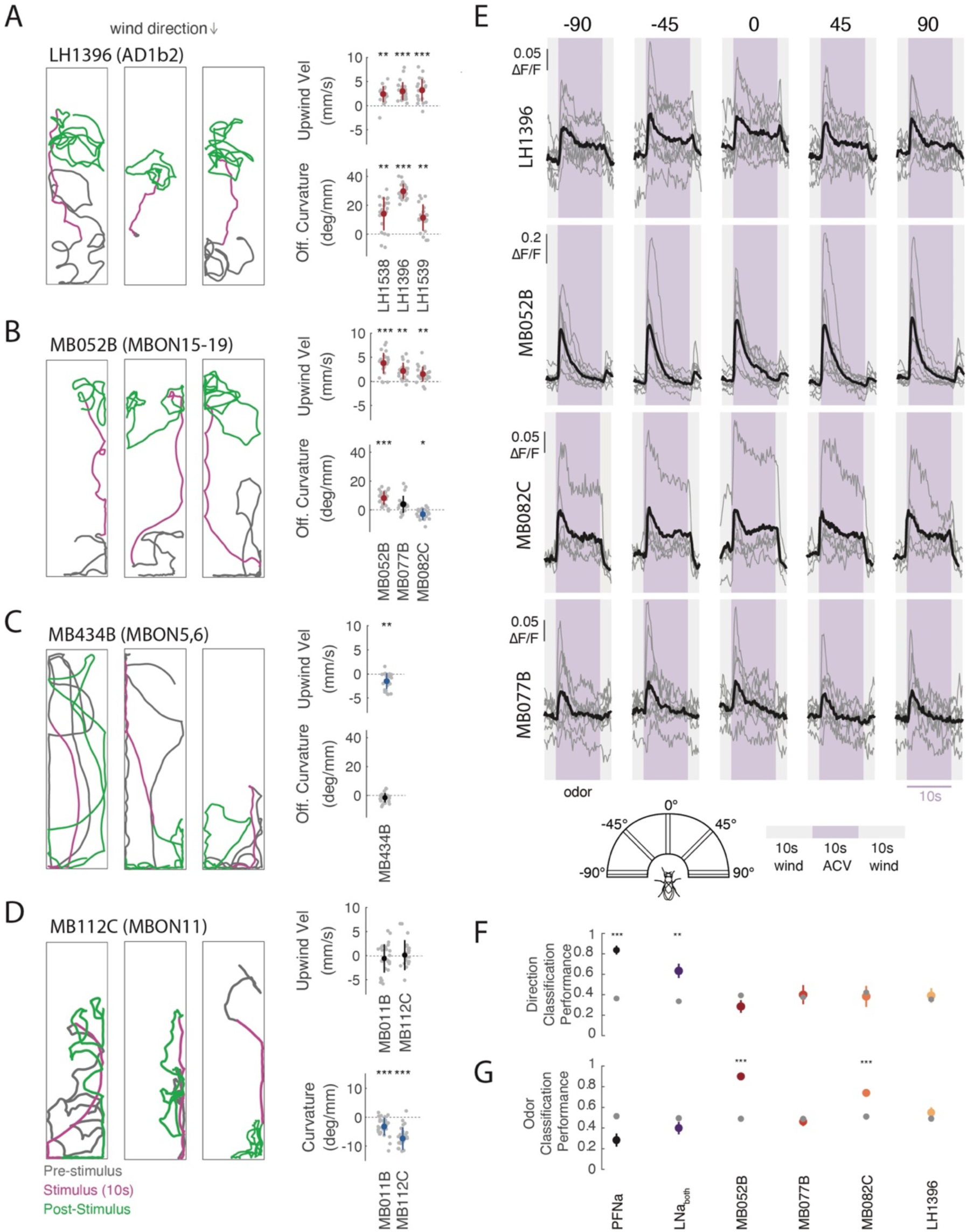
LH and MB output neurons promote wind navigation behavior but encode odor independent of wind direction. **A)** Optogenetic activation of AD1b2 LHONs drives upwind movement and OFF search. Left: Example behavioral trajectories in response to optogenetic activation of AD1b2 LHONs labelled by the line LH1396 (left). Right: Upwind velocity and OFF curvature (quantified as in Fig. 1D) for three lines that label AD1b2 LHONs (LH1538, LH1539, LH1396). All three lines significantly increase both upwind velocity and OFF curvature (Wilcoxon signed rank test, upwind: p= 2.4548e-04, 7.9941e-05, 1.8215e-05; OFF-curvature: p= 3.6712e-04, 1.7699e-04, 1.8215e-05 respectively) **B)** Optogenetic activation of attraction-promoting MBONs drives upwind movement and OFF search. Left: Example behavioral trajectories in response to optogenetic activation of MBONs15-19 labelled by the line MB052B (left). Right: Upwind velocity and OFF curvature quantification for three cholinergic MB lines: MB052B, MB077B, and MB082C. Each line labels distinct MBONs. All increase upwind velocity (Wilcoxon signed rank test, p= 1.0997e-05, 1.2267e-04, 2.0378e-04) while MB052B increases OFF curvature (p= 6.2811e-06), MB077B does not (p= 0.0046) and MB082C reduced OFF curvature (p= 0.0018). **C)** Activation of aversion-promoting MBONs promotes downwind movement. Left: Example behavioral trajectories in response to optogenetic activation of glutamatergic MBONs 5 and 6, labeled by the line MB434B (left). Right: Upwind velocity and OFF curvature for MB434B. MB434B significantly decreases upwind velocity (Wilcoxon signed rank test p= 2.2806e-04) but does not OFF curvature (p= 0.0258). **D)** MBONs promoting straighter trajectories. Example behavioral trajectories in response to optogenetic activation of the GABAergic MBON11, labeled by the line MB112C (left). Right: Curvature during stimulus (from 2-5s after stimulus ON) for MB112C, and MB011B. MB112C and MB011B significantly reduce curvature during the stimulus (Wilcoxon signed rank test, MB112C: p= 3.5150e-06, MB011B: p=3.407e-05). **E)** Upwind-promoting LHONs and MBONs encode odor independent of wind direction. Calcium responses (ΔF/F) measured in four lines that all drove upwind movement. Responses were measured in LH dendritic processes of LH1396 (N = 8 flies), in output processes of MB052B (N = 9 flies), MB082C (N=5), MB077B (N=8). All responses measured using GCaMP6f in response to odor (10% vinegar, purple) and wind (gray) delivered from 5 directions (schematic). Gray traces represent individual flies, black traces represent mean across flies. No significant differences in response as a function of direction were observed for either LH1396, MB052B, MB082C or MB077B (ANOVA: F(4,35)=0.35, p=0.8408, F(4,40)=0.3, p=0.8794, F(4,25)=0.2, p=0.9989, F(4,35)=0.67, p=0.6166). **F)** Performance of a wind direction (left, center, right) tree classifier trained on the first 5s of the odor period. Gray dots represent a classifier trained with the same data and shuffled labels. Student’s t-test PFNa p= 3.5063e-09, LNa p= 5.3035e-04, MB052B p= 0.1044, MB077B p= 0.7911, MB082C p= 0.7116, LH1396 p= 0.6229. **G)** Performance of an odor versus wind classifier trained on the first 5s of wind or odor. Gray dots represent a classifier trained with the same data and shuffled labels. Student’s t-test: PFNa p= 0.0657, LNa p= 0.3231, MB052B p= 2.7204e-08, MB077B p= 0.4104, MB082C p= 4.2903e-04, LH1396 p= 0.0909. All comparisons for behavioral data, and classifier performance were corrected using the Bonferroni method (see Methods).

We also identified MBONs that produced other navigational phenotypes. For example, the glutamatergic (inhibitory) MBON line MB434B (labeling MBONs 5 and 6), which was previously shown to produce aversion (Aso et al. 2014), generated downwind movement in our paradigm (Fig. 2C). Moreover, two MBON lines produced straightening (reduced curvature) in our paradigm (Fig. 2D) but no change in movement relative to wind (Fig. 2D, Fig. S2A): the GABAergic line MB112C (labeling MBON 11), which evoked attraction in quadrant assays, and the glutamatergic line MB011B (labeling MBONs 1,3,4), which evoked aversion (Aso et al. 2014). Overall, these results indicate that LH/MB outputs can drive coordinated “suites” of locomotor behavior that promote attraction or aversion in different environments. Several LHONs and MBONs redundantly drive upwind movement, key to attraction in windy environments, while other MBONs drive straightening, which promotes attraction in odor gradients (Schulze et al. 2015) or in response to familiar visual stimuli (Ardin et al. 2016).

Are the LHONs and MBONs that drive upwind movement sensitive to wind direction, or do they encode a non-directional odor signal that is integrated with wind direction downstream? To answer this question, we used calcium imaging to measure responses to calibrated wind and odor stimuli delivered from 5 different directions (Fig. S2D). Across all four upwind-promoting lines (MB052B, LH1396, MB077B, and MB082C), we observed responses to vinegar, but no tuning for wind direction (Fig. 2E, S2E, ANOVA: F(4,35)=0.35, p=0.8408, F(4,40)=0.3, p=0.8794, F(4,25)=0.2, p=0.9989, F(4,35)=0.67, p=0.6166). We separately used electrophysiology to show that the α’3 compartment of the MB, which was previously shown to respond to airflow (Mamiya et al. 2008), does not encode wind direction (Fig. S2F). To rigorously test whether MB/LH responses carry wind direction information, we generated tree classifiers (see Materials and Methods) and asked them to decode if wind was presented from the left, right or center relative to the fly (Fig 2F) based on responses during the odor period. We found that classifiers trained on MB/LH responses performed no better than shuffled controls (Fig. 2F). In contrast, classifiers trained on the responses of wind pathway neurons (PFNa, or the difference between left and right LNa neurons, Currier et al. 2020) performed significantly better than shuffled controls. We trained a second classifier to discriminate between odor and wind ON (Fig. 2G). In this case, MB052B and MB082C performed significantly better than control, while neither wind pathway neuron showed significant discrimination. All neurons significantly discriminated odor from baseline (Fig. S2G). We also applied our wind direction classifier to other phases of the response, such as wind ON and OFF (Fig. S2H). LH1396, but no other neuron group or phase, showed some discrimination at wind ON. This was largely due to responses that were stronger in front of the fly than at the sides. Taken together, this analysis supports the idea that MB/LH neurons that promote upwind orientation largely encode odor presence independent of wind direction.

### Multiple tangential FB inputs promote upwind orientation and respond to odor

The FB is anatomically downstream of the MB and LH (Li et al. 2020, Scaplen et al. 2020) and has previously been shown to encode wind direction. We therefore asked whether inputs to the FB are likewise capable of driving movement relative to wind direction. We first confirmed that the FB is anatomically downstream of our neurons of interest by performing anterograde trans-synaptic tracing (Talay et al. 2017) on two of our lines that drove upwind movement (MB052B and LH1396) and observed signal in the dorsal layers of the FB in both cases (Fig. 3A).

**Fig. 3:**
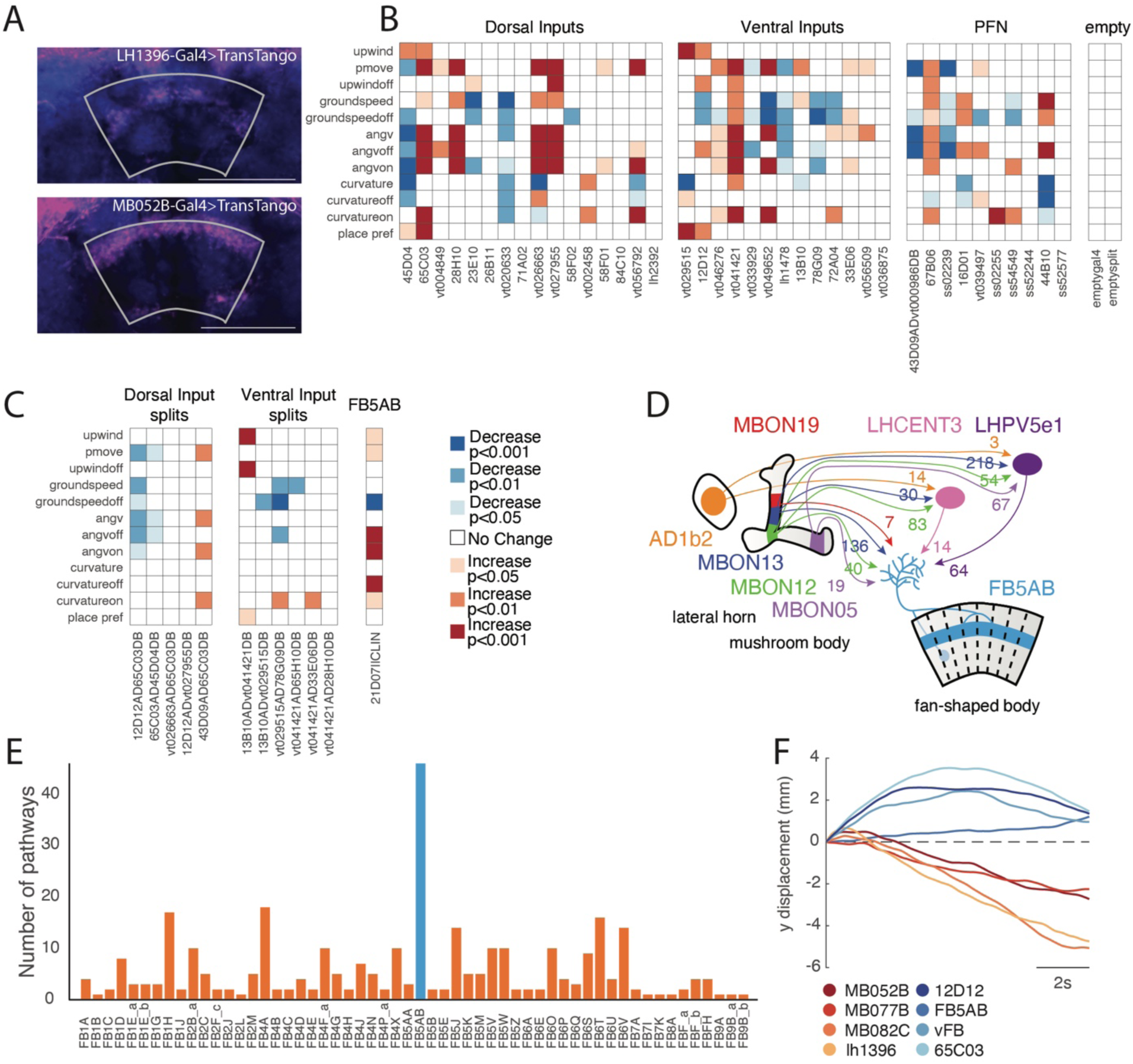
A set of FB tangential inputs promote wind navigation behavior. **A)** Trans-synaptic tracing reveals connections between upwind-promoting MB/LH neurons and FB tangential neurons. *Trans-tango* signal driven by LH1396-GAL4 (top) and MB052B-GAL4 (bottom). Trans-synaptic signal (magenta) was observed in horizontal layers of the dorsal FB in both cases. Neuropil is shown in blue. The FB is outlined in gray. Scale bar 50uM. **B)** Optogenetic activation results for FB inputs, including dorsal tangential inputs, ventral tangential inputs, columnar PFNs, and empty-GAL4 and empty-split-GAL4 controls. Two dorsal inputs and two ventral inputs drove significant increases in upwind velocity. Control lines drove no significant change in any measured behavioral parameter. See Materials and Methods for calculation of behavioral parameters. **C)** Optogenetic activation results for split GAL4 lines labeling dorsal and ventral tangential FB inputs, and for a line labeling FB5AB (21D07-GAL4||CLIN). One split-GAL4 line labeling ventral FB tangential inputs drove a significant increase in upwind velocity. **D)** Schematic showing feedforward connectivity onto FB5AB from three upwind-promoting MBONs (MBON 19, MBON 12, MBON 13), one upwind-promoting LHON (AD1b2), and one downwind-promoting MBON (MBON05). Pathways converge onto FB5AB directly or indirectly through LHCENT3 and LHPV5e1. Numbers represent the average synaptic weight between each cell type and the right-sided LHCENT3 (id: 487144598), LHPV5e1 (id: 328611004), or right-sided FB5AB (id: 5813047763). **E)** Number of parallel lateral horn pathways from vinegar-responsive projection neurons (estimated from ORN responses in Jung et al., 2015) to each FB tangential input neuron. Pathways consist of two synapses: the first between the projection neuron and lateral horn neuron, and the second between the lateral horn neuron and the FB tangential input neuron. Blue bar represents the number of pathways converging onto FB5AB. **F)** Upwind displacement responses to optogenetic activation of FB tangential input lines outlast the stimulus while responses to activation of MB/LH lines do not. Timecourses of average relative y-displacement (arena position) across flies, following stimulus OFF for each line. Individual fly’s average positions across trials were set to 0 and relative change in position for 10s following stimulus OFF was averaged across flies for each genotype.

To ask whether the FB plays a role in wind-guided navigation, we performed an activation screen of 40 lines labeling FB input neurons, including dorsal and ventral FB tangential inputs, and columnar PFNs, as well as additional Cx neurons (Fig. 3B, Fig. S3). We performed this screen using genetically blind flies (see Materials and Methods) and in the presence of teashirt-Gal80 (Clyne and Miesenbock, 2008) to reduce potential Chrimson expression in the ventral-nerve cord (VNC) (see Methods). We found that 4 lines labeling FB tangential inputs, but no lines labeling PFNs, generated significant movement upwind (Fig. 3B, S4A). Two dorsal FB input lines that were previously shown to promote sleep (23E10 and 84C10, Donlea et al. 2014) did not produce any wind-oriented movement in our assay, although we did observe a decrease in groundspeed in 23E10. FB tangential lines driving upwind phenotypes targeted both dorsal and ventral layers of the FB (Fig. 4A). We attempted to refine these lines by making split-Gal4 drivers from combinations of these hemidrivers, but only one of these, labeling a set of ventral FB tangential inputs, also drove an upwind phenotype (Fig. 3C). Most split-Gal4 drivers labeled only a very small number of neurons.

**Fig. 4:**
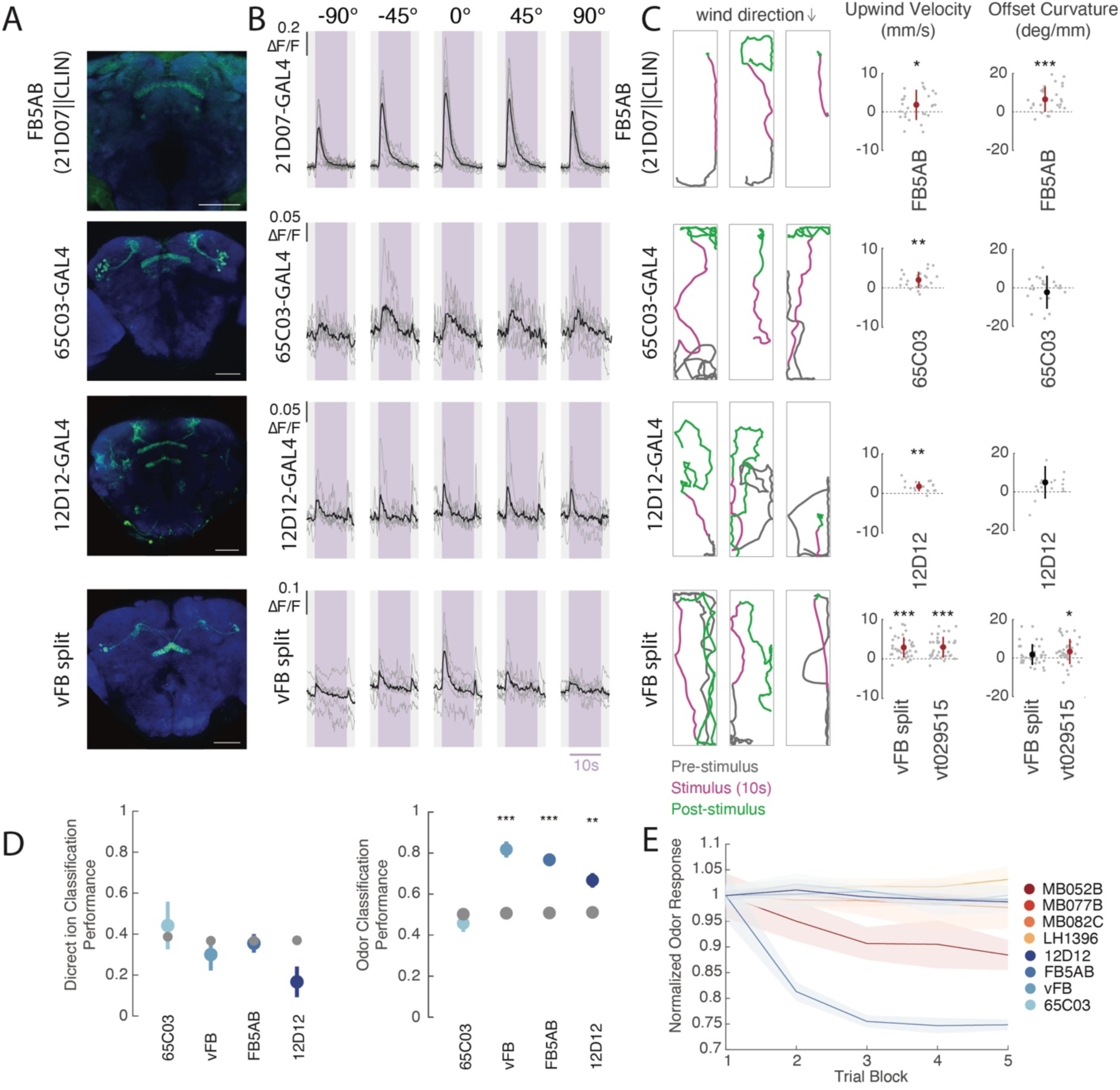
Multiple FB tangential inputs respond to attractive odor and drive upwind movement. **A)** Confocal images of lines that showed FB responses to vinegar and drove upwind movement when activated. Each image shows stain for mVenus expressed with UAS-Chrimson in flies of the same genotype used for activation experiments (abbreviated genotypes shown at left). **B)** Calcium responses in each line shown at left in response to odor delivered from 5 different directions (as in Fig. 2E). Shaded purple region indicates odor period. Gray lines represent individual flies and black represents the mean across flies (21D07-GAL4: N = 9, 65C03-GAL4: N = 7, 12D12-GAL4: N = 6, vFB split: N = 6 flies). No significant difference in response magnitude was observed between directions (ANOVA: 21D07-GAL4: F(4,40)=2.14, p=0.0938, 65C03-GAL4: F(4,30)=0.68, p=0.6096, 12D12-GAL4: F(4,25)=0.13, p=0.971, vFB split: F(4,25)=0.64, p=0.6358). **C)** Example behavioral trajectories and quantification of upwind velocity and OFF curvature in each line shown at left. For FB5AB only light intensity was 34µW/mm^2^. For all drivers, upwind velocity was quantified over 0-10s after odor ON. Right: Mean±STD overlaid upwind velocity, OFF curvature (Wilcoxon signed rank test: 21D07||CLIN: p =0.0306, 8.1448e-05, 65C03-GAL4: p=4.1850e-04, 0.5841, 12D12-GAL4: p=1.8218e-04, 0.0055 vFB split: p=3.4153e-07, 0.1155, VT029515-GAL4 p=4.3255e-07, 0.0024). **D)** Performance of tree classifiers for wind direction (left) and odor presence (right) for FB tangential inputs. Gray dots represent classifiers trained with the same data and shuffled labels. Left: Performance of wind direction (left, center, right) classifier trained on the first 5s of odor period. Student’s t-test 65C03 p=0.6450, vFB p=0.4100, FB5AB p=0.7882, 12D12 p=0.0177. Right: Performance of odor versus wind classifier trained on first 5s of wind and odor ON. Student’s t-test: 65C03 p=0.0203, vFB p=1.0934e-06, FB5AB p=2.8375e-04, 12D12 p=0.0383. **E)** Decay of fluorescence response to odor over trial blocks. The response to each trial block was calculated as the average odor response to 5 consecutive trials (each from one of the directions), averaged across all flies of a genotype. Response magnitude was normalized by the average response to the first block for each line. Shaded area represents standard error across flies. All comparisons for behavioral data and classifier performance were corrected using the Bonferroni method (see methods)

In addition to the lines identified through our screen, we identified the neuron FB5AB using the connectome (Fig. 3D,E, Fig, 4A). This single pair of neurons stood out as the only FB input that receives at least one direct synaptic input from each of the MBON lines with wind navigation phenotypes (Fig. 3D). In addition, FB5AB receives the largest number of di-synaptic LH inputs from vinegar-responsive glomeruli of any FB input neuron (Fig. 3E, see Methods). We identified a GAL4 line that labels FB5AB neurons (21D07). As 21D07 labels some neurons in the antennal lobe, a primary olfactory area, we used the cell class-lineage intersection (CLIN, Ren et al. 2016) technique to limit expression of Chrimson to neurons in the FB (see Methods). High intensity light activation of this driver, which weakly but specifically labels FB5AB, also drove upwind orientation (Fig. 3C, 4C).

Overall, upwind velocity responses to FB tangential input stimulation were more persistent than those evoked by optogenetic activation in the MB and LH, continuing to promote upwind displacement after light OFF, rather than evoking search behavior (Fig 3F, S4A). We characterized the neurotransmitter phenotypes of each of the hits from our screen and found that all were cholinergic, and thus excitatory (Fig, S4B). As in the MB and LH, silencing of individual FB input lines that drove upwind movement was not sufficient to block upwind movement in response to odor (Fig. S4C, Fig. 6H). Together these results support the hypothesis that patterns of population activity in FB tangential inputs can promote upwind movement.

We next sought to characterize the sensory responses of upwind-promoting FB tangential inputs by performing calcium imaging in response to wind and odor from different directions (Fig. 4B, S4D). We observed responses to vinegar in all but one line (45D04). No FB tangential line showed significant directional tuning (ANOVA: 21D07: F(4,40)=2.14, p=0.0938, 65C03: F(4,30)=0.68, p=0.6096, 12D12: F(4,25)=0.13, p=0.971, vFB split: F(4,25)=0.64, p=0.6358) nor were tree classifiers based on their odor responses able to distinguish between odor delivered from the left, right or center (Fig 4D, S4E). In contrast, tree classifiers trained to distinguish odor ON from wind ON performed better than shuffled controls for 3 of 4 lines (vFB split, FB5AB, and 12D12, Fig. 4D), while all performed better than shuffled control when trained to distinguish odor from baseline (Fig. S4F). The largest responses were observed in FB5AB (21D07), although these (but not other FB tangential line responses) decayed over trials (Fig 4E). In the line 65C03, we observed odor responses only from the dorsal layers of the FB, while in the line 12D12, we observed odor responses only from the ventral layers of the FB. Together these data identify a population of olfactory FB tangential inputs, targeting multiple layers of the FB, that respond to attractive odor and promote upwind movement.

### hΔC neurons encode a wind direction signal that is modulated ON by odor

Our functional imaging data suggest that, like the upwind-promoting neurons in the MB/LH, FB tangential inputs are not directionally tuned for wind. In contrast, PFNa and LNa encode wind direction, but not the presence of vinegar (Fig. 2F). Together these data suggest that columnar and tangential inputs to the FB encode directional and non-directional information respectively. We therefore hypothesized that FB local neurons, which receive input from both columnar and tangential inputs, might integrate these two types of information.

To identify FB local neurons that integrate odor and wind direction signals, we used the hemibrain connectome (Scheffer et al. 2020) to search for neurons that receive input both from FB5AB and from the most wind-sensitive PFNs in our previous survey (PFNa, p, and m, Currier et al. 2020). This analysis revealed a population of 20 hΔC neurons that tile the vertical columnar structure of the FB (Fig. 5A, Hulse et al. 2021). Stains revealed these neurons to be cholinergic (Fig. S5A). Each hΔC neuron receives input from wind-sensitive PFNs at its ventral dendrites, and from FB5AB at its dorsal axons, where excitatory olfactory input might gate synaptic output (Fig. 5B,C).

**Fig. 5:**
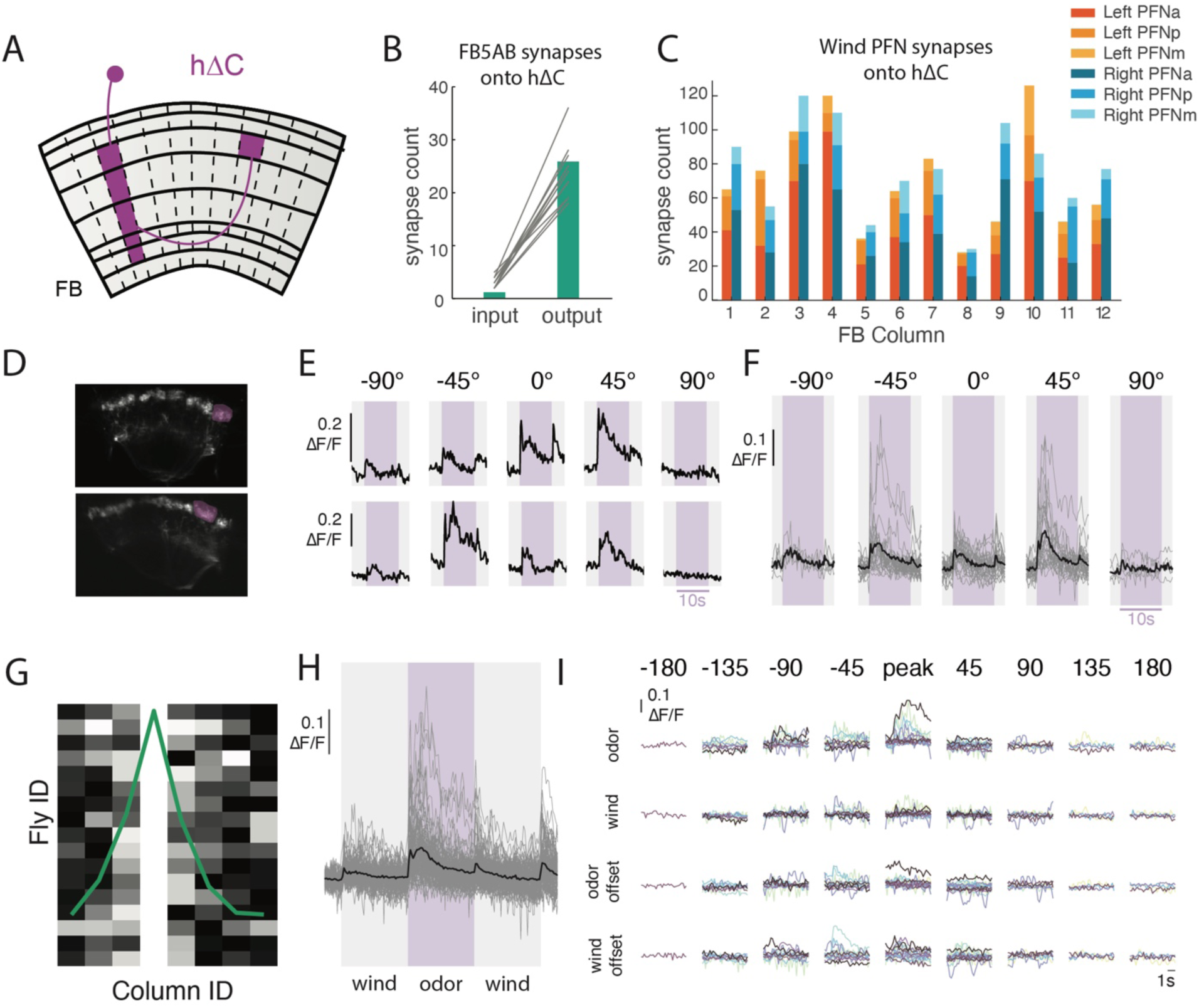
hΔC neurons exhibit odor-modulated wind direction tuning. **A)** Schematic of an individual hΔC neuron’s anatomy (purple). The 20 hΔC neurons that tile the FB each receive input in horizontal layers 2-6 in a single column, and project halfway across the FB, making outputs in horizontal layer 6. **B)** Number of FB5AB synapses onto the input and output tufts of hΔC for every FB5AB-hΔC pair. **C)** Number of synapses from left and right wind-sensitive PFNs (PFNa, PFNp, and PFNm, tuned to 45° left and right respectively) onto hΔC neurons, summed within columns. **D)** 2-photon image of tdTOM expressed with GCaMP6f under VT062617-GAL4 which labels hΔC neurons. Purple ROI drawn around output tuft of a single column for analysis. **E)** Odor responses of two example flies/columns showing directionally-tuned odor responses. **F)** Summary of all measured odor responses > 2STD above baseline across columns and flies. Gray traces represent individual columns and black traces represent mean across columns. **G)** Directional responses are restricted to nearby columns: maximally responsive direction for each fly, where data is phase shifted so maximum columns align at column 4. Each fly normalized to maximum column response. **H)** Summary of wind and odor responses for all measured responses > 2STD above baseline across columns and directions. Responses to odor are stronger than those to wind ON, odor OFF and wind OFF (n=87 responsive columns from N=16 Flies). **I)** Summary of wind and odor phase activity of maximally responsive column, aligned to maximally responsive direction. Data is shifted to be aligned to the maximal direction and each row represents different stimulus period. Each color represents a different fly (N=16).

To ask whether hΔC neurons respond to wind and odor signals, we identified a GAL4 line—VT062617— that labels hΔC neurons, and performed calcium imaging from the FB while presenting odor and wind from different directions. We observed calcium responses in the dorsal FB, where hΔC neurons form output tufts (Fig. 5D, S5B). In several examples, we observed odor responses that were strongest for a particular direction of wind (Fig. 5E). hΔC responses were strongest to odorized wind from +45° and -45° (Fig. 5F), although the relationship between wind direction and peak response was not consistent across flies (Fig. 5E). hΔC responses were typically localized to a few nearby columns (Fig. 5G), and we observed no consistent relationship between column location and preferred wind direction when pooling data across flies (Fig. S5C). Thus, the wind direction representation in hΔC neurons appears to be unique to each fly, similar to what has been observed for heading representations in compass neurons (Seelig and Jayaraman 2015), and distinct from the wind representation in PFNs, which is uniformly tuned to 45° ipsilateral (Currier et al. 2020).

We next examined hΔC responses as a function of stimulus phase. Across flies and columns, the strongest and most consistent responses occurred during the odor, although weaker responses also occurred at wind ON, odor OFF, and wind OFF (Fig. 5H). To examine whether wind tuning differed across these phases, we selected the maximally responsive column for each fly, and aligned responses to the peak direction of that response, then plotted responses to other phases of the stimulus relative to that wind direction. Responses in other stimulus phases tended to be in the same direction or nearby directions to the odor response (Fig. 5I). Taken together, our results suggest that hΔC neurons exhibit a bump of activity that depends on wind direction, but whose columnar position is unique for each fly. This bump is activated most strongly during odor, although it can also appear more weakly during other phases of the stimulus.

### hΔC neurons promote diverse navigation phenotypes and contribute to persistent upwind walking

Do hΔC neurons contribute to navigation behavior? To address this question, we activated hΔC neurons optogenetically using various spatial activation patterns, and examined the resulting behavioral trajectories (Fig. 6). Because our imaging data suggest that hΔC neurons show a bump of activity during odor and wind stimulation, we first asked what behavior was produced by sparsely activating hΔC neurons. We used SPARC2-I (Isaacman-Beck et al. 2020) to activate a random ∼15% of hΔC neurons in each fly while they walked in the presence of laminar wind. Activation of sparse subsets of hΔC neurons (Fig, 6A) caused many flies to re-orient and then walk in a specific reproducible direction (Fig. 6B,C). Each fly walked in a distinct direction, suggesting that this was specified by the particular pattern of hΔC neurons activated in that fly (Fig. 6A,B). Directions were biased towards the long axis of our arena, but equally distributed up- and downwind (Fig. 6C). Directional walking was not due to the arena walls, as fly behavior near the walls was excluded from our analysis (see Materials and Methods). To assess the significance of this directional walking, we compared the strength of orientation in hΔC>SPARC flies to empty-GAL4>SPARC flies (Fig. 6C,D, S6A). Empty-Gal4>SPARC flies showed significantly weaker orientation behavior (Fig. 6D). To determine whether the pattern of hΔC activation was related to walking direction, we dissected and stained each hΔC>SPARC fly. However, we observed no relationship between the anatomy of the activated columns, and the direction or strength of oriented walking (Fig S6A-E). We interpret these results to suggest that sparse activation in hΔC neurons can produce a reliable heading, and that this heading is encoded in fly-specific (not fixed) coordinates.

**Fig. 6:**
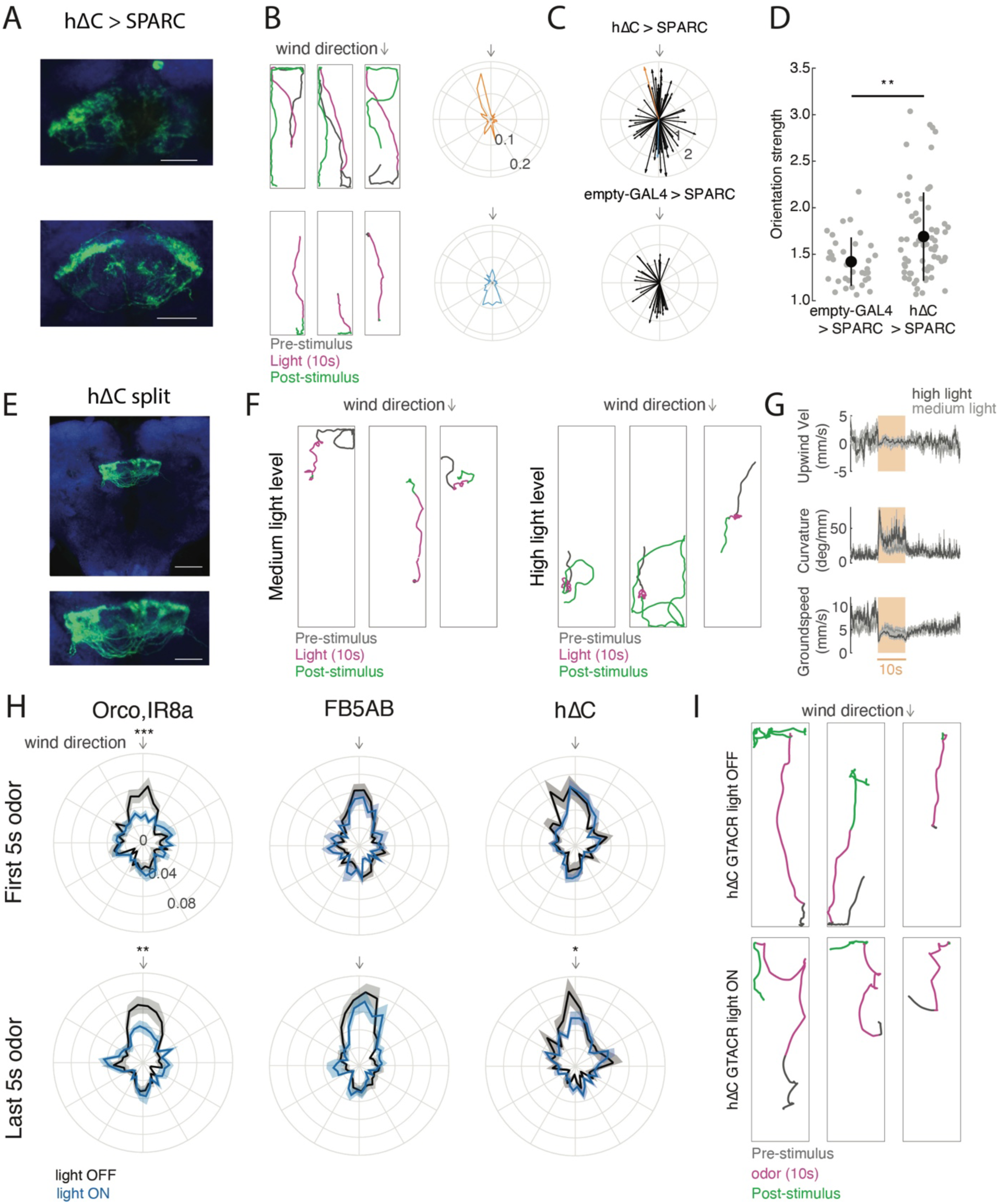
Role of hΔC neurons in navigation behavior. **A)** Anatomical and behavior data from two representative hΔC > SPARC flies. Left: confocal images of tdTomato expressed with UAS-SPARC2-I-Chrimson. Scale bar 25µm. **B)** Behavior of two representative hΔC > SPARC flies from **A**. Left: example behavioral trajectories of flies before, during, and after optogenetic activation. Right: orientation histograms for each fly for the period 2-6 s after light ON. Radial axis represents probability. **C)** Preferred orientations across all flies, where the vector direction corresponds to the preferred direction and the vector strength corresponds to the orientation index (see Materials and Methods, Fig. S6A). Top: hΔC > SPARC. Representative flies from A-B) are shown in orange and blue. Bottom: empty-GAL4 > SPARC. **D)** hΔC > SPARC flies show stronger oriented walking than empty-GAL4 > SPARC2 flies (see Fig. S6A for Methods, ranksum test p = 0.0035). **E)** Confocal image of mVenus expressed with UAS-Chrimson by the hΔC split-GAL4 line 19G02AD;VT062617DB. Scale bar 50µm (top), and 25µm (bottom). **F)** Example behavioral trajectories driven by hΔC split-GAL4 activation with 26 µW/mm^2^ light (left) or 34 µW/mm^2^ light (right) **G)** Upwind velocity, curvature, and groundspeed across all hΔC split-GAL4 flies before, during, and after optogenetic activation using 26 µW/mm^2^ light (light gray) or 34 µW/mm^2^ light (dark gray). **H)** Optogenetic inactivation of hΔC neurons using GtACR disrupts persistent upwind orientation. Each plot shows orientation histograms during light-evoked silencing (blue) compared to no-light control (black) for the first 5s of odor (top) and last 5s of odor (bottom). Shaded regions represent SEM. Optogenetic silencing of ORNs (Orco,IR8a) significantly reduces the probability of orienting upwind (+/- 10°) during both phases (p=9.2724e-04 early, 0.0090 late), while silencing of FB5AB does not (p=0.3848 early, p=0.3259 late). Silencing of hΔC neurons reduces upwind orientation only during the late phase (p=0.8512 early, p=0.0475 late). **I)** Example behavioral trajectories in hΔC > GtACR flies in response to odor. Top: blue light off (non-silenced). Bottom: blue light on (hΔC neurons silenced).

In addition, we asked whether broad activation of hΔC neurons could drive a behavioral phenotype. For these experiments, we generated a split-GAL4 line selectively labeling hΔC neurons, and expressed Chrimson throughout this population (Fig. 6E). Activation of this line caused unstable reorientation behaviors. At medium light levels (26µW/mm^2^), fly walking was interrupted by frequent left and right turns, while at higher light levels (34µW/mm^2^), flies displayed tight turns that were clockwise, counterclockwise, or alternating between clockwise and counterclockwise (Fig. 6F). Across flies, these behaviors were captured by an increase in curvature with no change in wind upwind velocity (Fig 6G). Similar increases in curvature were obtained using a variety of drivers for hΔC neurons (Fig. S6F). Therefore, uniform activation of hΔC neurons causes unstable reorientation behaviors.

Finally we asked whether hΔC activity is required for upwind orientation. In preliminary experiments, we found constitutive silencing of hΔC neurons to be lethal. We therefore used the light-activated chloride channel GtACR (Mohamed et al. 2017) to acutely silence hΔC neurons during odor presentation. We compared the effects of acutely silencing hΔC neurons to silencing of olfactory receptor neurons using orco/IR8a and FB5AB. Consistent with our constitutive silencing results (Fig. 1F,G), we found that acute silencing of olfactory receptor neurons impaired upwind orientation throughout the odor period (Fig. 6H). In contrast, silencing of FB5AB had no effect on upwind orientation, as we observed for other FB tangential inputs (Fig. 6H, Fig. S4C). In flies with hΔC neurons silenced, we observed normal upwind orientation early in the odor period, however trajectories then deviated significantly from upwind later in the odor period (Fig. 6H,I). We conclude that hΔC activity is required for persistent upwind fixation throughout the odor stimulus.

### A computational model of hΔC neurons’ contribution to navigation behavior

Our data suggest that hΔC activity contributes to persistent upwind orientation during odor and that sparse activation of hΔC neurons can promote navigation in a reproducible direction. To understand how different patterns of hΔC neuron activation can evoke diverse behavioral phenotypes, we developed a computational model. The elements of our model are all based on neurons and connection motifs (direct or indirect) found in the hemibrain connectome (Fig. 7A,C, Scheffer et al. 2020, Hulse et al. 2020), although we imagine that additional neurons and connections may play a role in natural olfactory navigation behavior.

**Fig. 7:**
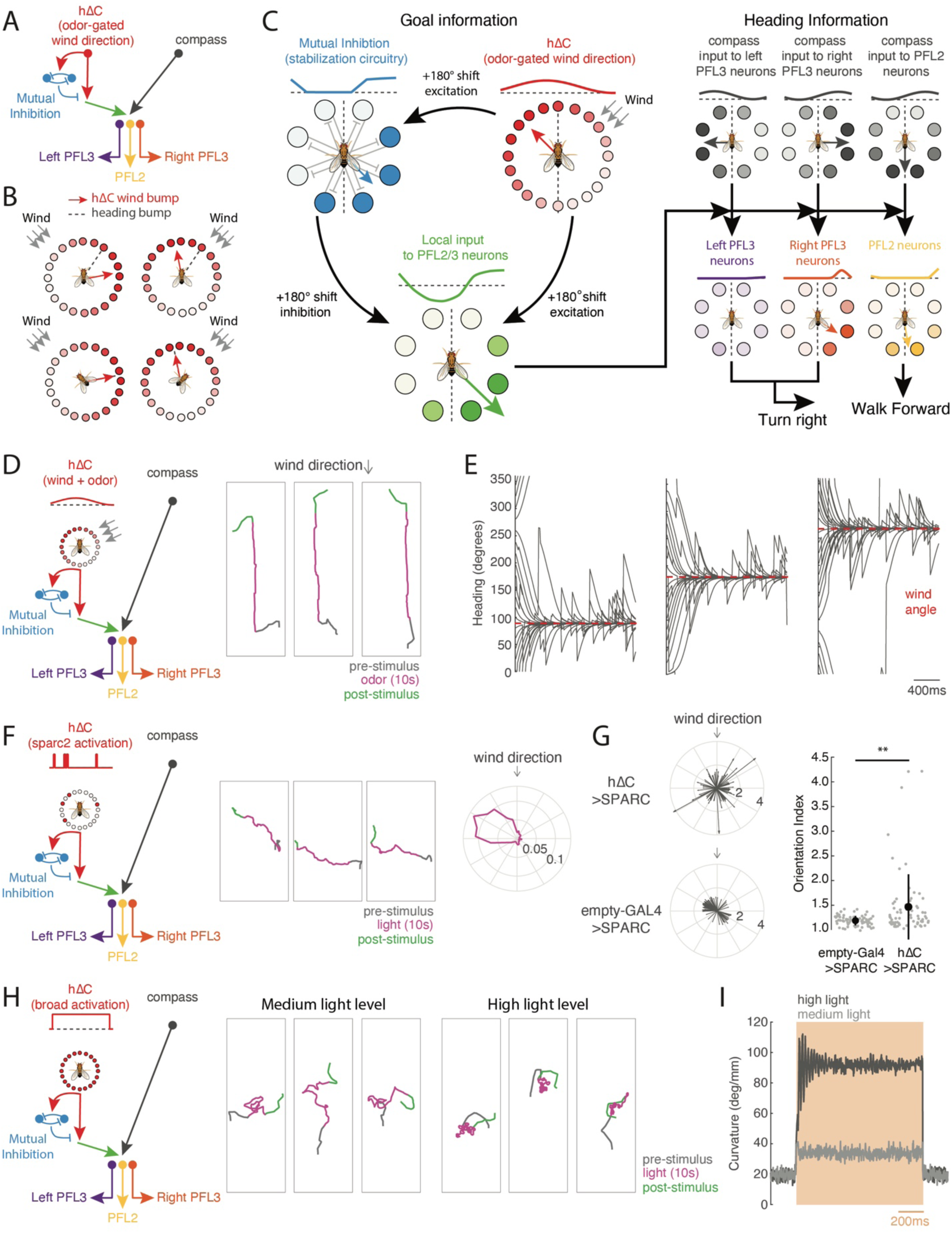
A model of FB circuitry that translates hΔC activity into goal-directed walking. **A)** Overview of cell types and information flow in an FB circuit model. Odor gates a wind direction signal in hΔC neurons (red), which is processed by a mutual inhibition circuit (blue) as well as fed forward to PFL output neurons (green). PFL neurons (purple, yellow, orange) integrate goal information from hΔC with heading information from compass neurons (black) to control turning (PFL3) and forward velocity (PFL2). **B)** Modeled allocentric wind representation in hΔC neurons, shown as both a bump of activity (circles) and as a vector (red arrow). Heading vector is displayed in gray. Leftward rotations of the fly cause rightward rotations of the heading vector, consistent with observed motion of the heading representation in compass neurons during movement of the visual field (Seelig and Jayaraman 2015). Note that the hΔC vector represents the wind direction in allocentric coordinates, i.e. the bump does not move as long as the wind comes from the same world direction (columns). In addition, the wind and heading vectors only align when the fly is pointed upwind (bottom row). When the fly is not pointed upwind, the wind vector is to the right of the heading vector for leftward wind and to the left of the heading vector for rightward wind (top row). An alternate wind representation is depicted in Fig. S7A. **C)** Detailed diagram of the model circuit showing the transformation of an hΔC wind representation into upwind movement by PFL3 and PFL2 neurons. For each step, activity is represented as a bump of activity across the FB (lines, with dotted line = 0 activity), a bump of activity across FB cells (circles), and as a vector. An odor-gated wind bump in hΔC neurons (red) is fed forward directly to PFL2/3 neurons (green) as well as indirectly via a mutual inhibition circuit (blue) that helps stabilize the activity pattern. 180° shifts in the output of each neuron group ensure that the wind bump in hΔC neurons is transformed into a stable bump in PFL2/3 neurons (green). PFL3 neurons (purple/orange) receive a heading bump from the compass system (black) that is shifted by 90° ipsilateral, as well as goal input from hΔC (green). When these bumps overlap, the inputs sum constructively, as shown here for right PFL3 neurons (orange) that drive right turns. When these bumps do not overlap, the inputs sum destructively, as shown here for the left PFL3 neurons that drive left turns. Total turning is driven by the sum of left and right PFL3 activity. PFL2 neurons (yellow) receive a heading bump from the compass system (black) that is shifted by 180°, together with goal input from hΔC (green). When these bumps overlap the inputs sum constructively to promote faster forward walking. This arrangement ensures that fly turns until the goal and heading bumps align, which happens when the fly faces upwind, and increases its speed when oriented upwind. **D)** Simulated circuit activity and trajectories when odor gates the expression of an allocentric wind bump as in **B**. Example trajectories (right) are shown for 3 model flies. In this simulation hΔC activity is only present during odor. **E)** Headings of simulated flies in response to odor with wind arriving from 90° (left), 180° (center), and 270° (right) and different initial headings. Note that the heading always converges to upwind despite turning noise which drives the fly off course. **F)** Simulated circuit activity during sparse optogenetic activation. In this simulation, 15% of hΔC neurons are activated for the duration of the light stimulus. The same neurons are activated in a single model fly, while different neurons are activated in different flies. Example trajectories (center) are shown for one model fly on three different trials. Note that the fly always converges to the same reproducible walking direction. Orientation histogram (right) calculated as in Fig. 6B. **G)** Preferred orientations across simulated hΔC > SPARC flies (top) and empty-GAL4 > SPARC flies (bottom) calculated as in Fig. 6C. Empty-GAL4 > SPARC flies were simulated by setting hΔC activity to zero. Simulated hΔC > SPARC flies have stronger orientation indices than simulated empty-GAL4 > SPARC flies (ranksum test p = 0.0032). **H)** Simulated circuit activity during broad optogenetic activation. In this simulation, every hΔC neuron is activated equally for the duration of the stimulus, at a medium (center) or high (right) level. **I)** Average curvature across simulated flies before, during, and after broad optogenetic activation using medium light (light gray) or high light (dark gray). Broad activation drives an intensity-dependent increase in curvature.

**Fig. 8:**
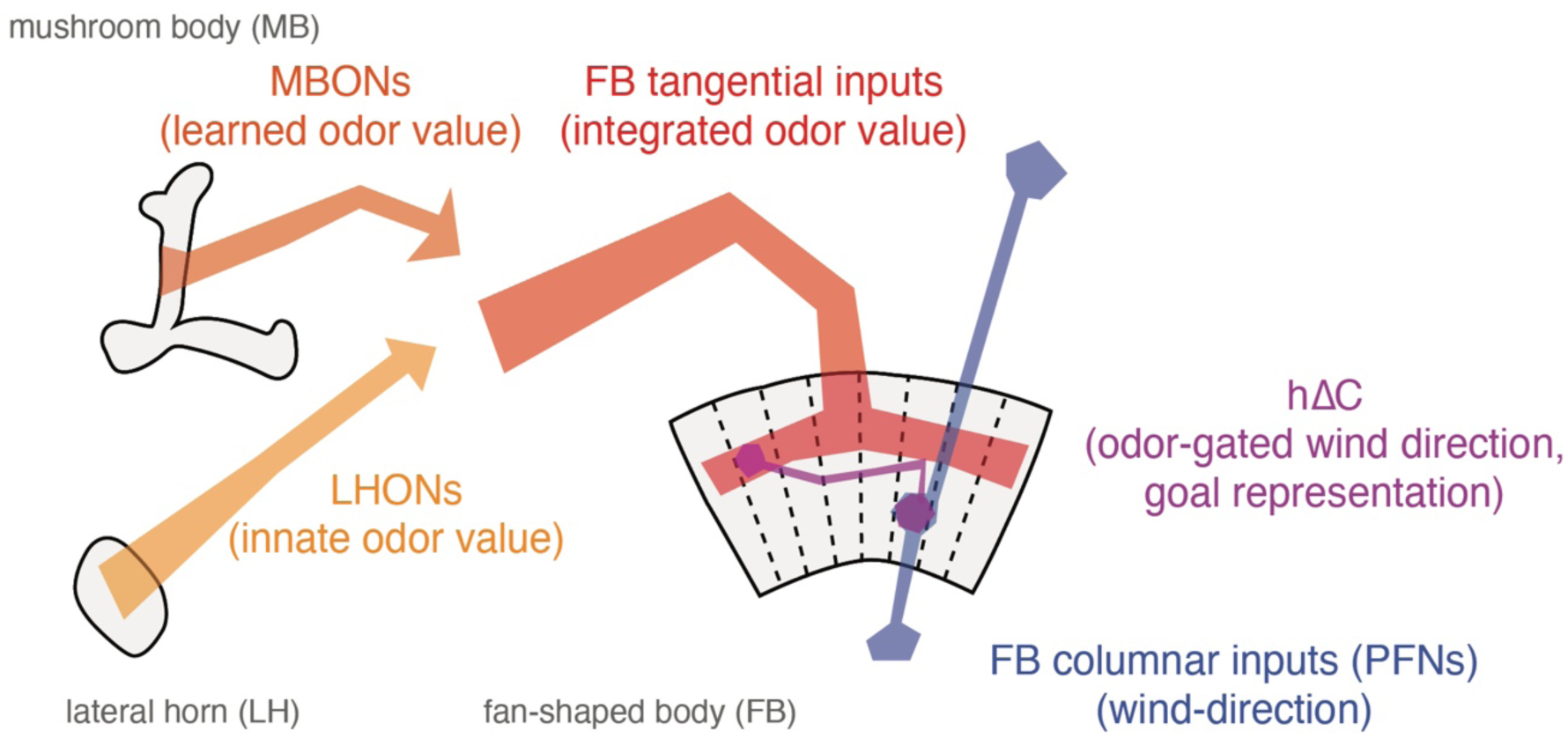
Conceptual model of sensory integration for olfactory navigation in the *Drosophila* central brain. Conceptual model of central olfactory navigation circuitry as suggested by this and previous studies. MBONs, LHONs, and FB tangential inputs promote wind navigation and encode odor information but not wind direction information. FB tangential inputs are a likely locus where learned and innate odor information may be integrated to drive behavior. In contrast, FB columnar inputs (PFNs) encode wind direction but not odor presence (Currier et al. 2020). hΔC neurons receive input both from directionally-tuned PFNs and odor-tuned FB tangential inputs, and encode a fly-specific wind direction signal that is modulated ON by odor. Sparse activation of hΔC neurons can drive movement in a reproducible direction and activity in these neurons is required for sustained upwind orientation during odor. Our data support a model in which columnar and tangential inputs to the FB encode directional and non-directional information respectively, and these two inputs are integrated by local neurons to specify navigational goals.

We first asked whether odor-gated wind direction information in hΔC neurons could be used to promote upwind movement. Although the nature of the wind representation in hΔC neurons is not entirely clear from our imaging experiments, our results are broadly consistent with a bump of activity that is tuned to wind-direction in a fly-specific manner, likely reflecting the fly-specific coordinates of the compass in each animal (Seelig and Jayaraman, 2015). Thus, we imagine that hΔC neurons might show a bump of activity whose location reflects the allocentric wind direction (Fig. 7B), as has previously been described for traveling direction signals in other FB local neurons (Lyu et al. 2021, Lu et al 2021). Alternatively, hΔC might show a bump of activity related to wind direction only in the frontal hemisphere, which could be computed based on the known frontally-tuned responses of wind-sensitive PFNs (Fig. S7A,B). In either case, in our model this bump becomes active when the fly encounters odor, due to gating input from FB5AB and other FB tangential neurons, although it could in principle also be activated at other times.

Activity in hΔC neurons is then translated into locomotor commands by a set of previously described output neurons: PFL3 and PFL2 (Fig. 7C, see Materials and Methods). PFL3 neurons project unilaterally to the right or left LAL and are thought to drive turning (Rayshubskiy et al. 2020, Stone et al. 2017, Hulse et al. 2021, Sun et al. 2020, Goulard et al. 2021), while PFL2 neurons project bilaterally to the left and right LAL and are hypothesized to modulate forward walking speed (Hulse et al. 2021). PFL neurons compare the “goal” representation arriving from hΔC neurons (a bump of activity flipped by 180° due to hΔC neurons projecting to columns halfway across the FB) with a shifted representation of the fly’s current heading arriving from compass neurons (Stone et al. 2017, Hulse et al. 2021). The heading representation in PFL3 neurons is shifted by -90° for right PFL3s and +90° for left PFL3s. This allows PFL3 neurons to determine whether a left or right turn will bring the fly in line with the “goal” heading specified by the hΔC bump. For example, wind to the right of the fly will generate a wind bump that overlaps more with the right PFL3 heading bump than the left (Fig. 7C). This system will cause the fly to turn until its heading bump aligns with the hΔC wind bump. In parallel, PFL2 neurons cause the fly to accelerate when the “goal” heading from hΔC overlaps with the compass heading representation. This heading representation is shifted by 180° so it will overlap the wind bump when wind is directly in front of the fly. Thus, PFL2 neurons drive the fly to walk faster when it is pointed upwind.

We added one additional element to our model that has not been previously described. This is a second set of mutually-inhibitory FB local neurons (light blue in Fig. 7C) that receive input from hΔC neurons and also send information to the PFL output system. This motif is not observed in hΔC neurons but reciprocal connections between opposing columns can be seen in several other FB local neuron types downstream of hΔC, such as hΔA, hΔG, hΔH, and hΔM, which all send strong projections to PFL3. As we show below, this motif is required to account for the unstable reorientation behavior we observe during broad activation of hΔC neurons.

We first asked whether this simplified FB model could promote upwind movement when the hΔC wind bump is activated (Fig. 7D,E). We simulated the circuit described above using a network of firing rate-based neurons (see Materials and Methods), and simulated odor input by turning on the bump of wind activity in hΔC neurons. As predicted, the odor-gated wind bump in hΔC neurons generated stable upwind orientation that was robust to allocentric wind direction and initial heading (Fig. 7E). Flies also oriented upwind using a frontal wind representation in hΔC, although this was less reliable than using an allocentric representation (Fig. S7A-D). We did not observe local search after odor OFF in our simulations, consistent with our finding that FB tangential inputs do not drive this behavior.

We next asked whether the same model could recapitulate the directional walking we observed with sparse random hΔC activation. To simulate the sparse activation experiment, we replaced the hΔC wind bump with random activation of 15% of the hΔC population in a fly-specific pattern (Fig. 7F). Similar to our behavioral results, simulated flies reoriented and then walked in an arbitrary but reproducible direction, where the direction was stable for each fly. This result occurred because the random optogenetic input provided stable non-uniform input to the PFL3 neurons, and the maximum of this input (i.e., the FB columns opposite to the highest density of ‘active’ hΔC neurons) establishes a stable direction. These directions were distributed in 360° space (Fig. 7G) because our simulation lacked arena constraints. Oriented walking was significantly reduced when we simulated empty-GAL4 > SPARC controls by omitting hΔC activity from our simulation (Fig. 7G).

Finally we asked whether our model could reproduce the unstable reorientation we observed with broad hΔC activation (Fig. 7H). To simulate these experiments, we uniformly activated hΔC neurons at two different intensities, corresponding to the two light levels in our experiment. Like our behavioral results, low activation generated walking interrupted with left and right turns, while high activation generated tight turns that were clockwise, counterclockwise, or alternating between clockwise and counterclockwise. Consistent with our experiments, simulated flies displayed elevated curvature that depended on the light level (Fig. 7I). The unstable reorientation behaviors in our simulations stemmed from oscillations within the mutual inhibition layer of the circuit; in the absence of this layer, uniform activation did not alter turning statistics (Fig. S7E-G). Taken together, our experiments and modeling suggest that sensory representations in FB local neurons such as hΔC can provide a “goal” heading that promotes stable navigation towards an environmental target such as an upwind odor source.

## Discussion

### Distinct direction and context pathways for olfactory navigation

Wind-guided olfactory navigation is an ancient and conserved behavior used by many organisms to locate odor sources in turbulent environments. Despite large differences in the types of odors they seek, and in the physics of odor dispersal, very similar behaviors have been observed in pheromone-tracking moths (Kennedy et al, 1974), food-seeking crustaceans (Page et al. 2011), and ants seeking both food and their nest (Buehlmann et al. 2012). Thus, this behavior is required in many animals for survival and reproduction, and likely mediated by conserved neural circuits.

In a previous study, we identified inputs to the FB that encode wind direction (Currier et al. 2020). Neurons throughout this pathway, including both LNa neurons and PFNs, show activity that is strongly modulated by wind direction, but weakly modulated by odor. Other LN and PFN neurons have been shown to strongly encode optic flow direction (Stone et al, 2017, Lyu et al. 2020), another cue that signals self-motion and wind direction— particularly in flight. Taken together, current data thus suggest that the columnar pathway to the FB encodes information about self-movement and environmental flow useful for recognizing where the fly is relative to its environment.

In contrast, the present study identifies an olfactory pathway to the FB that encodes odor information largely independent of the wind direction from which it is delivered. We found that neurons in both the MB (a center for associative learning) and the LH (a center for innate olfactory and multi-modal processing) are capable of driving upwind movement. However, very few of these upwind-promoting neurons exhibited wind-direction tuning, particularly during the odor period. Thus, the outputs of these regions likely represent odor identity or value signals, as has been suggested by several recent models (Ardin et al., 2016, Cognini et al., 2018). We further characterized a large population of olfactory tangential inputs to the FB that likewise promote upwind movement and encode odor independent of wind direction. At least one of these neurons is anatomically downstream of upwind-promoting MB and LH neurons. These data are consistent with previous studies showing that other FB tangential inputs encode non-spatial variables such as sleep state (Donlea et al. 2014) and food choice (Sareen et al. 2021). Together, these data suggest that the pathway running from the MB/LH to the FB encodes non-spatial contextual information.

The segregation of direction and context inputs to the FB that we observe here shows some similarities to the organization of visual circuits in primates and of navigation circuits in rodents. In primate vision, processing of object location and object identity are famously thought to occur in distinct pathways (Mishkin et al. 1983). In the hippocampus, inputs from the medial entorhinal cortex encode spatial cues, while anatomically distinct inputs from the lateral entorhinal cortex encode non-spatial context cues, including odors (Hargeaves et al. 2005, Leitner et al. 2016). Thus, a segregation between the computation of spatial/directional information and context/identity information may be a general feature of central neural processing in both vertebrate and invertebrate brains.

What might be the advantage of this type of organization? One hypothesis is that it allows learning about non-spatial cues to be generalized to different spatial contexts. For example, during food-guided search, innate information about which odors signal palatable food must be integrated with learned odor associations (Hige et al. 2015b, Schlegel et al. 2021), as well as with internal state variables such as hunger (Sayin et al. 2019, Sareen et al. 2021). However, the fly may wish to generalize information learned while walking to search in flight, as has been demonstrated in bees (Chaffiol et al. 2005). In walking flies, wind direction can be computed directly from mechanosensors in the antennae (Bell and Kramer, 1979, Suver et al. 2019). In flight however, wind only transiently activates mechanosensors (Fuller et al. 2014), but then displaces the fly as a whole, leading to an optic flow signal opposed to the fly’s direction of movement (Kennedy et al. 1940, van Breugel et al. 2014). As different PFNs carry airflow and optic flow signals in a similar format (Lyu et al. 2020, Currier et al. 2020), this circuit may be well-poised to compute wind direction in both walking and flying flies. Separating the computation of stimulus value, in the MB/LH and tangential FB inputs, from the computation of wind direction, in PFNs, may allow flies to generalize stimulus associations learned in one context (walking) to another (flight).

### Specification of a goal direction by FB local neurons

How does the nervous system represent goals for navigation? When sensory cues are continuously available, no explicit goal representation may be required, and movement towards a goal such as food may be accomplished through chains of sensory-motor reflexes (van Bruegel et al. 2014, Schulze et al. 2015, Alvarez-Salvado et al. 2018). In contrast, when sensory cues are unavailable, such as during navigation towards a remembered shelter, the nervous system must build and store a representation of the goal location (Stone et al. 2017).

Navigation towards an odor source presents an interesting intermediate case. Food sources generally produce strong sensory cues that can be used to drive chained reflexes. However, the wind direction and odor cues produced by turbulent plumes are fluctuating, variable, and uncertain, meaning that memory and internal estimates of source position can make an important contribution to effective navigation (Vergassola et al. 2007, Masson 2013, Grünbaum and Willis, 2015). Understanding how nervous systems represent and use internal variables during this task may help us to design robust search algorithms for noisy environments.

Here we present evidence that hΔC neurons participate in specifying a goal heading during olfactory navigation. Silencing of hΔC neurons impairs persistent upwind orientation during odor, suggesting that the activity of these neurons is required to maintain upwind fixation. hΔC neurons receive input from an odor-tuned, upwind-promoting pathway through the MB/LH to FB tangential neurons. Our imaging results are consistent with a model in which hΔC neurons represent wind direction (likely through inputs from wind-sensitive PFNs) and modulate the strength of this representation during odor (perhaps through axo-axonic gating input from FB tangential neurons). Thus, hΔC neurons are well-poised to construct an internal representation of the goal direction during olfactory navigation. However, understanding the precise nature of wind, odor, and goal representations in hΔC neurons will require future imaging of population activity during ongoing navigation.

We also show here that sparse random activation of hΔC neurons can evoke locomotion in a reproducible direction, consistent with the idea that hΔC activity patterns can specify a goal heading. We were unable to determine from our data whether hΔC activation specifies a distance (and thus a vector) as well as a direction, although many trajectories from the same fly were of similar length. Curiously, we found that hΔC activity had to be non-uniform in order to evoke directional walking, but need not be organized as a “bump” of activity— any asymmetry in the response across columns was sufficient. Although we produced this activity through sparse optogenetic stimulation, similar asymmetries could be generated and stored as patterns of asymmetric synaptic input from FB tangential neurons onto FB local neurons, or as asymmetric synaptic weights between interconnected FB local neurons. Because the Drosophila FB contains at least 27 local neuron types targeted by ∼145 tangential neuron types (Hulse et al. 2020), this structure could provide a reasonably large capacity for storing diverse direction or vector memories.

To understand how hΔC activity might specify a goal heading, either upwind or in an arbitrary reproducible direction, we developed a computational model. This model is based on motifs found in the *Drosophila* connectome, such as the phase shifts between compass neurons and PFL neurons (Stone et al. 2017, Hulse et al. 2020) and mutual inhibition found in certain local neuron populations. However, the connection schemes used in our model are simplified compared to the real circuit. In our model, the architecture of the FB is used to compare a *goal* heading in hΔC neurons with the *current* heading represented in compass neurons, and then drive turning to minimize discrepancies between these directions. This computation is implemented by the convergence of hΔC input and compass neuron input onto PFL output neurons that control steering and forward velocity. Our steering model is similar to several recent models proposed for path integration and visual landmark-guided navigation (Stone et al. 2017, Hulse et al. 2020, and Goulard et al. 2021). However, our model differs in the proposed location and nature of the goal representation. In previous models, goals were stored in PFN activity (Stone et al. 2017), or in plastic synapses between compass neurons and PFLs (Goulard et al. 2021). Here the goal is instead stored in the dynamic activity pattern of a population of local neurons. One advantage of our model is that different local neurons can be rapidly switched on and off through tangential input, allowing the fly to rapidly update its goals depending on behavioral demands. A further advantage is the large number of local neurons and tangential inputs, which as noted above provide a substrate for learning, storing, and releasing multiple goal memories.

Although our results implicate hΔC neurons in the integration of odor and wind cues and the specification of a target walking direction, flies with hΔC neurons silenced were still able to initially turn upwind in response to odor, suggesting that other neurons play a role in this initial upwind turn. These might be additional FB local neuron types— we identified multiple FB tangential inputs that drive upwind movement and not all of these target hΔC. They may also include neurons that bypass the central complex. In addition to targeting the FB, MBONs also make direct connections to the lateral accessory lobe (LAL) a region implicated in motor control and steering, including odor-evoked steering behavior in moths (Namiki and Kanzaki, 2016, Rayshubskiy et al. 2020, Scaplen et al. 2021, Li et al., 2020). As the LAL also receives wind direction input (Okubo et al. 2019), it could form a second site of wind and odor integration. In this study, we found that activation of FB neurons produced more persistent wind orientation phenotypes than activation of MB/ LH output neurons. Thus, parallel pathways from olfactory centers to motor centers could regulate behavior on different timescales. FB representations of odor and wind might also allow a fly to adopt courses at particular angles to the wind, or to generate an internal estimate of the direction of the odor source. Determining how activity in these different pathways shapes behavior on different timescales, and in different spatial contexts, will provide additional insight into the organization of central circuits for navigation.

## Materials and Methods

### FLY STOCKS

All experimental flies (except trans-tango flies described below) were raised at 25° C on a standard cornmeal-agar medium. Flies were raised with a 12h light-dark cycle. We used the following genotypes and ages for the experiments shown in each figure. Parental genotypes and sources are listed in the Key Resources Table below. For optogenetic activation experiments, experiments were run in male norpA hemizygotes, which are genetically blind, to eliminate any possible innate visual responses to red light. All other flies used were female. We detected no difference in olfactory behavior of male versus female flies in our assay in a previous study (Alvarez-Salvado et al., 2018). For calcium imaging experiments we used older flies (5-21 days) to maximize indicator expression. For electrophysiology we used younger flies to minimize glial ensheathing which make it challenging to obtain clean patch recordings. Trans-tango flies were raised at 18°C and aged until they were 10-20 days old following recommended protocols (Talay et al., 2017).

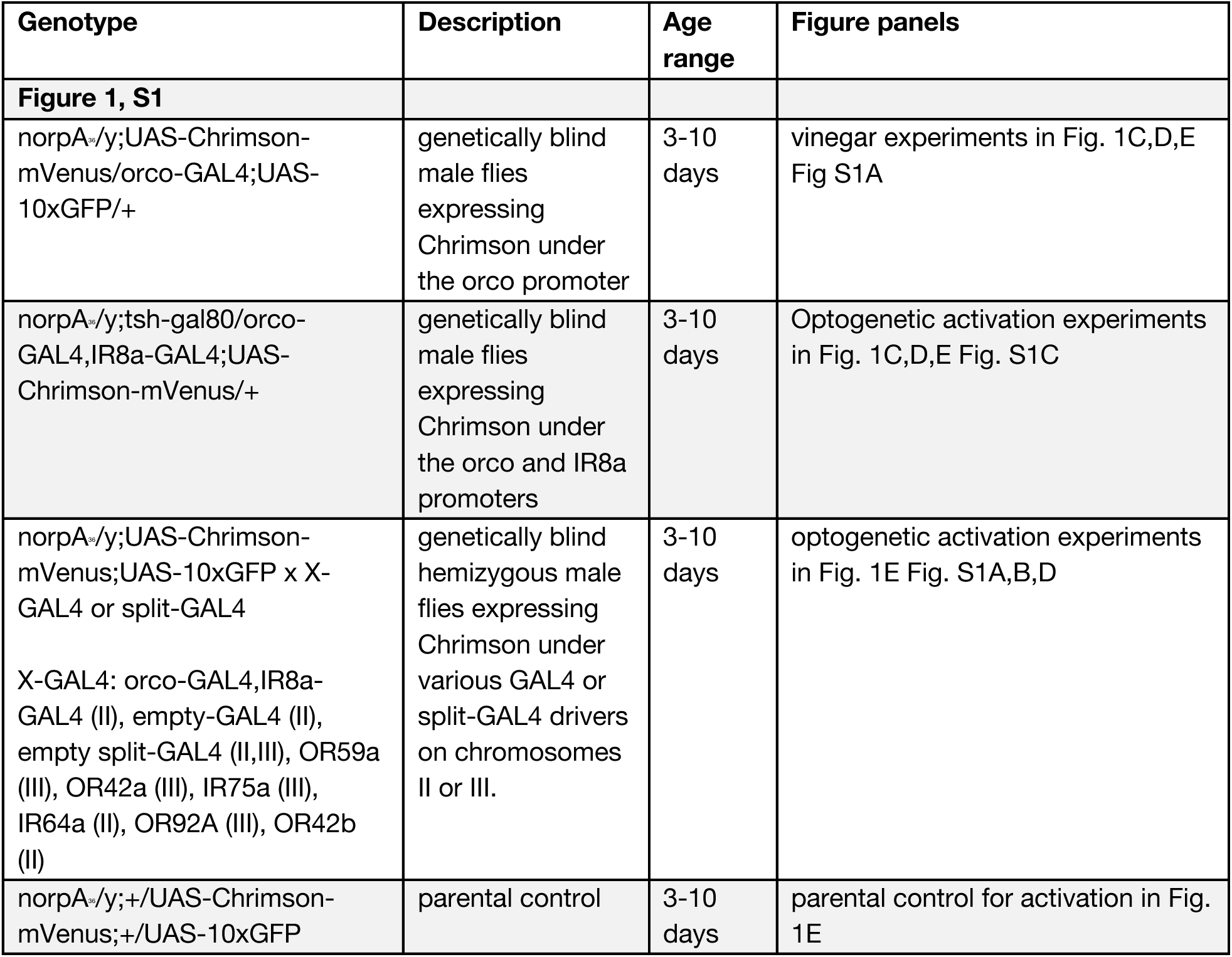
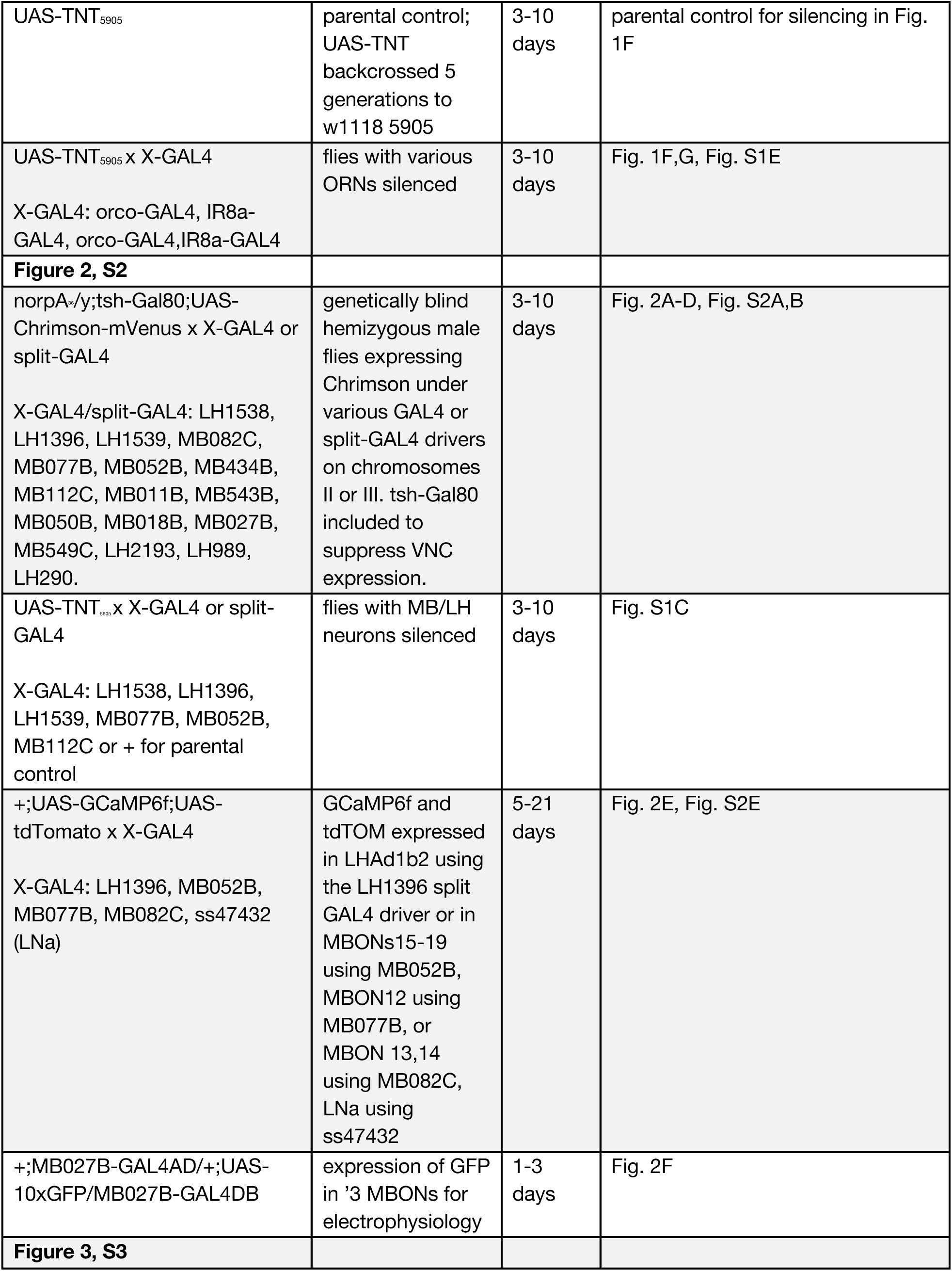
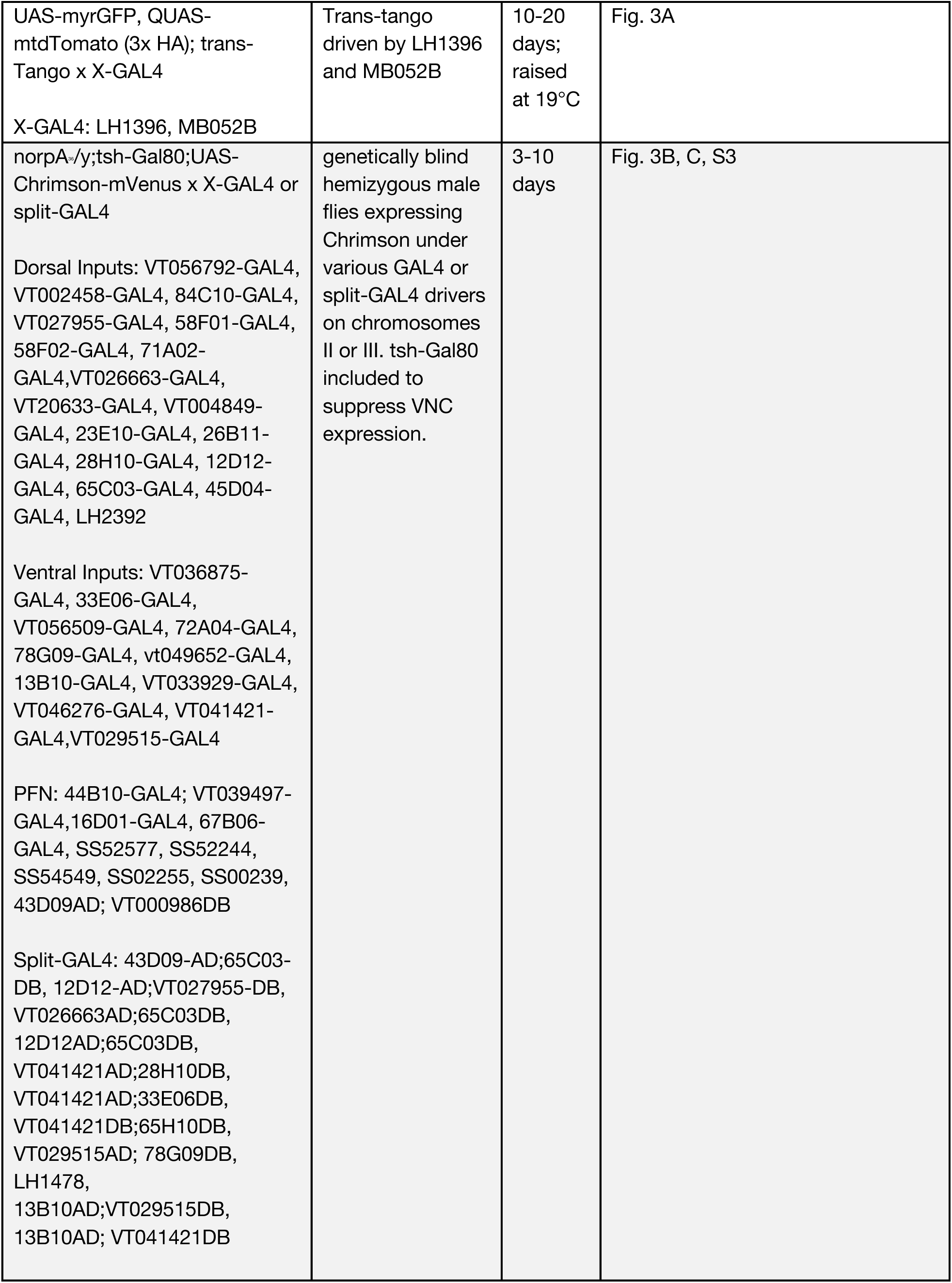
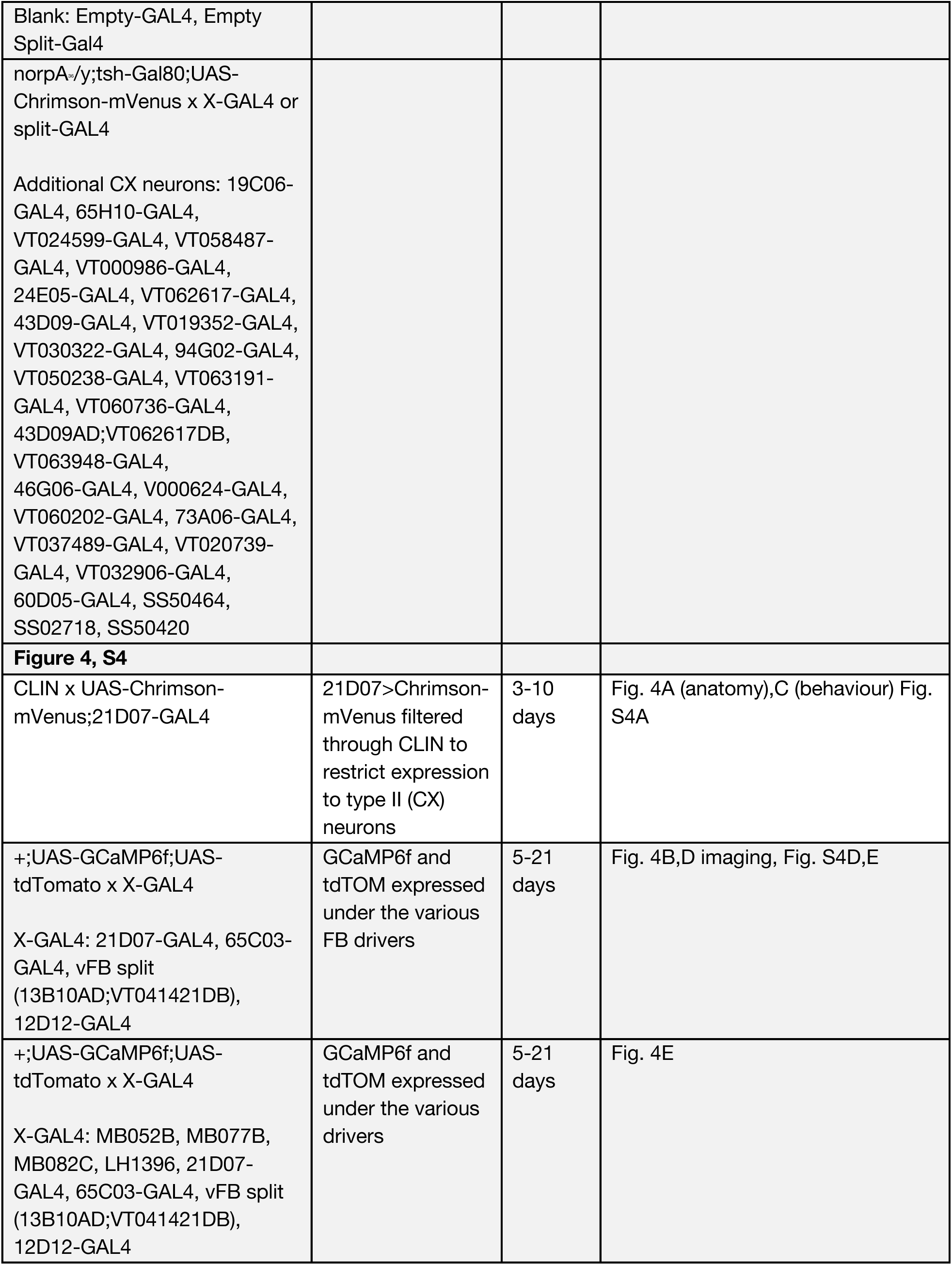
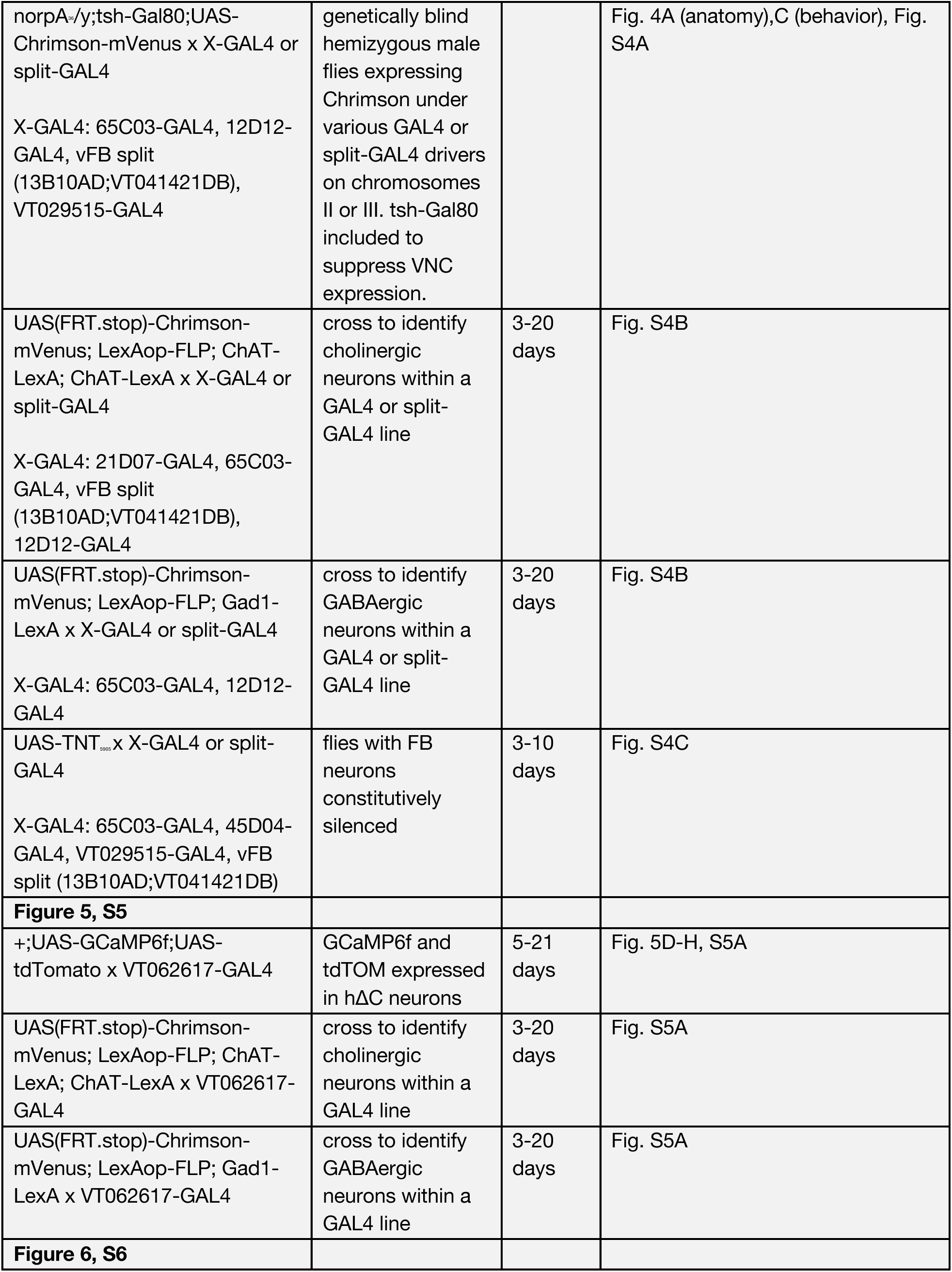
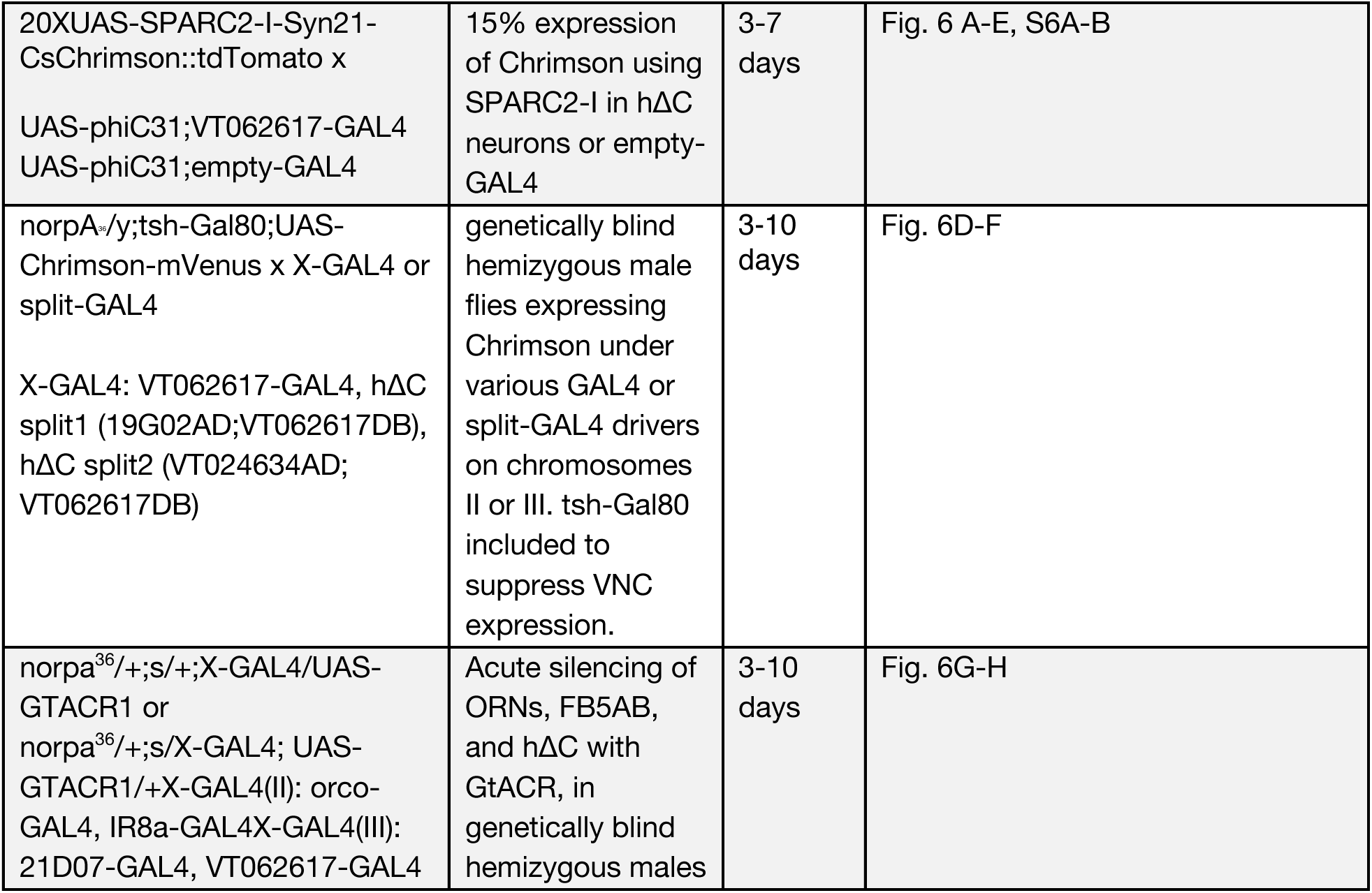

### BEHAVIORAL EXPERIMENTS

For behavioral experiments we used a modified version of the miniature wind tunnel setup described in Alvarez-Salvado et al. 2018. Briefly, flies were constrained to walk in a shallow arena with constant laminar airflow at 11.9 cm/s and tracked using IR LEDS (850nm, Environmental Lights) and a camera (Basler acA1920-155um). In experiments with odor, a 10s pulse of 1% apple cider vinegar (Pastorelli) was introduced through solenoid valves located immediately below the arena. For optogenetic activation experiments, we used red LEDs (626nm, SuperBrightLEDS), interleaved with the IR LEDs in a panel positioned 4.5cm above the arena. Light intensity was measured with a light meter (Thorlabs) and was 26 µW/mm^2^, measured at 626nm, for the majority of activation experiments. In flies expressing Chrimson using the CLIN technique the light level was 34 µW/mm (Aso et al., 2014) to compensate for low Chrimson expression in this line. For optogenetic silencing experiments, light panels with blue LEDs (470nm, Environmental Lights), were used. Light intensity for silencing experiments was 58 µW/mm, measured at 530nm. Wind and odor stimuli were calibrated using a hotwire anemometer (Dantec Instruments) and photo-ionization detector (miniPID, Aurora Systems) respectively.

Flies (males or mated females) were collected 2 to 7 days before the experiment and housed in time shifted boxes on 12h cycles. For optogenetic experiments, flies were placed on cornmeal-agar food supplemented with 50uL all-trans retinal (35mM stock: Sigma, R2500, dissolved in ethanol, stored at -20°C) mixed into ∼1 teaspoon of hydrated potato flakes. These vials were covered with aluminum foil except for a small window near the fly plug. All flies began the experiment (∼1.5-2h) between their subjective ZT1 and ZT4. 20-24h before the experiment we deprived flies of food by placing them in a polystyrene vial with damp shredded kimwipe. Flies were briefly anaesthetized by placing the vial on ice for ∼1min before loading into the arena. Flies were allowed 5-10 minutes to recover before the experiment began. Flies for genetic silencing were run in the dark, while optogenetic activation experiments were run with ambient lighting. During experiments with odor, flies received blocks of three trial conditions in random order: wind and odor (30s wind alone, 10s odorized air, 30s wind alone), wind alone (70s), or no wind (70s). During optogenetic activation experiments, flies received blocks of four trial conditions in random order: light stimulation with wind (30s wind, 10s wind and light, 30s wind), light stimulation alone (30s no stimulus, 10s light, 30s no stimulus), wind alone (70s wind), or blank trials (70s). During optogenetic silencing experiments, flies received blocks of four trial conditions in a random order: light stimulation with wind (25s wind, 15s light, 30s wind) odor stimulation with wind (30s wind, 10s odor, 30s wind), wind with overlapping light and odor, 25s wind, 15s light with 10s odor overlaid, 30s wind).

We analyzed behavioral data in a similar fashion as in Alvarez-Savaldo et al. 2018. X,Y coordinates and orientation were tracked in real time at 50Hz using custom Labview software (National Instruments). Data were further analyzed offline using custom MATLAB scripts. Coordinates and orientation were low-pass filtered at 2.5Hz using a 2-pole Butterworth filter. Any trials with tracking errors, or where the fly moved less than 25mm overall were discarded from further analysis. Any fly which moved on less than 5 trials for a condition was excluded as well. For all measured parameters (see below) time periods when the fly was stationary (moving at less than 1mm/s) were omitted. In trials with odor stimuli, we aligned the time courses of behavioral parameters to the actual time the fly encounters the odor based on their position is the arena, based on delays recorded by miniPID.

Behavioral parameters as a function of time were calculated as follows for each trial: Distance moved was calculated as the length of the hypotenuse between two pairs of coordinates at each frame. Groundspeed was calculated as the distance moved divided by the time interval of a frame (20ms). Upwind velocity was calculated as the change in Y-coordinates divided by the frame interval. Angular velocity was calculated as the absolute value of the change in unwrapped orientation divided by the time frame interval. Curvature was calculated as the filtered angular velocity divided by the filtered groundspeed. Probability of movement (pmove) was calculated by binarizing groundspeed with a threshold of 1 mm/s.

Quantification of average behavioral parameters represent average parameters across trials for each fly. All parameters were compared to a baseline period (10-25s in the trial) expect place preference where the baseline was taken from 25-30s into the trial (immediately before stimulus ON at 30s). We used the following time windows for analysis: probability of movement (pmove): 0-5s from stimulus ON, upwind velocity (upwind): 0-5s from stimulus ON for periphery, LH, MB, 0-10s from stimulus ON for FB, OFF upwind velocity (upwindoff): 0-2s after stimulus OFF, groundspeed: 2s-5s from stimulus ON, OFF groundspeed: 0-2s after stimulus OFF, angular velocity (angv): 2s-5s from stimulus ON, OFF angular velocity (angvoff): 0-2s after stimulus OFF, ON angular velocity (angvon): 0-1s from stimulus ON, curvature: 2s-5s from stimulus ON, OFF curvature (curvatureoff): 0-2s after stimulus OFF, ON curvature (curvature on): 0-1s from stimulus ON, place preference (placepref): 7.5s from stimulus ON to 2.5s after stimulus OFF. For display purposes we do not depict the significance values in tables (Fig 3. B,C, Fig S3) for upwind velocity for small magnitude increases (<1mm/s). We do not display values for curvature (both ON and OFF) if angular velocity is below <50deg/s and groundspeed is <4mm/s. We excluded these values because the curvature increase was due mostly to the drop in groundspeed and trajectories in these cases did not exhibit the characteristics of search. To examine the persistence of upwind displacement we looked at the change in displacement relative to position at stimulus OFF (Fig 3F). We averaged this across trials for individual flies, and averaged across flies for 10s following stimulus OFF for the depiction in the figure. To examine the orientation of flies during the odor period for optogenetic silencing experiments, we calculated the probability a fly had a given angular orientation across all trials while moving >1mm/s. We summed the probability that a fly was +/-10° from upwind, and compared this probability between odor and odor+light conditions using a ranksum test. For display we show polar histograms with 10° binning.

### SPARC2 BEHAVIOR AND ANATOMY

The orientation index and preferred direction of SPARC flies (VT062617-GAL4 or empty-GAL4) were determined by tracking the orientation of flies between 2 and 6 seconds after light ON. This range was selected to capture fly heading after flies made their initial turn, but before flies walked into the arena wall. Data points where the fly was walking slower than 1 mm/s or was positioned within 3 mm of the arena walls were excluded from analysis. We then computed a polar histogram of orientation values for all light periods in each fly. We computed the orientation index by taking the SVD of this histogram, projecting the data onto the two principal components (PC), and then calculating the ratio of the standard deviations along these two principal components. Histograms that show more pronounced orientation exhibit higher ratios. The preferred direction was determined by finding the direction along the first principal component with the highest value. Flies with fewer than 15 trials or fewer than 2000 orientation data points were excluded from analysis.

The Chrimson expression of hΔC > SPARC flies was analyzed using two methods: number of cell bodies and expression vector. The number of cell bodies was calculated by counting cell bodies near the FB. The expression vector was calculated by drawing an ROI around the dorsal FB (layer containing output tufts of hΔC neurons), splitting the FB into 12 columns, converting the normalized expression across these columns into a 360° polar plot, and then converting the polar plot representation into a vector. The vector length was compared to orientation index, while the vector direction was compared to preferred direction.

### CALCIUM IMAGING

For calcium imaging experiments, flies were cold-anaesthetized and mounted in a version of the fly holder described in Suver et al. 2019. The fly’s head was positioned in a keyhole shaped metal cutout (etchit) within a plastic holder. We attached the fly to the holder using UV glue (Riverruns UV clear Glue, thick formula), and stabilized the proboscis to the head/body, leaving the antenna free to move. We removed the front two legs to prevent interference with airflow stimuli. Under *Drosophila* extracellular saline (103 mM NaCl, 3 mM KCl, 5 mM TES, 8 mM trehalose dihydrate, 10 mM glucose, 26 mM NaHCO3, 1 mM NaH_2_PO_4_H_2_0, 1.5 mM CaCl_2_2H_2_O, and 4 mM MgCl_2_6H_2_O, pH 7.1-7.4, osmolarity 270-274 mOsm), we dissected away the cuticle at the back of fly’s head using fine forceps. We removed the trachea, airsacs, and muscle over the surface of the brain. Flies were starved for 18-24h prior to the experiment. External saline bubbled with carbogen (5% CO_2_, 95% O_2_) was perfused for the duration of the experiment.

2-Photon imaging was performed using a pulsed infrared laser (Mai Tai DeepSea, Spectraphysics) with a Bergamo II microscope (Thorlabs). Images were acquired through a 20x water immersion objective (Olympus XLUMPLFLN 20x) using ThorImage 3.0 software. The wavelength of the laser was set to 920 nm and power at the sample ranged from 13 to 66mW. Spectral separation of emitted photons was accomplished with two bandpass filters (red, tdTOM, 607/70nm, green, GCamp6f, 525/50nm) and detected by GaAsP PMTs. Imaging areas varied depending on the genotype but were between 47 x 47uM and 122 x 74uM. Imaging regions were identified first using the tdTOM signal under epifluorescence. Images were acquired at ∼5.0 frames per second. Across genotypes we excluded any flies where we were unable to obtain 5 trials of each direction, either due to fat migration or cell death. We excluded 2 flies from the 65C03-GAL4 data which showed rhythmic spike like activity and did not respond to any phase of our stimulus. For VT062617-GAL4 imaging we excluded 1/17 flies as no columns showed had an average response >2STD above baseline.

The majority of airflow and odor stimuli were delivered using a 5-direction manifold described in Suver et al. 2019 and Currier et al. 2020. Air was charcoal filtered, then passed through a flowmeter (Cole-Parmer), and proportional valves (EVP series, EV-05-0905; Clippard instruments laboratory,Cincinnati, OH) to direct air or odorized air at the fly from one of 5 directions. Airspeed was ∼25cm/s. For a small subset of additional experiments, we used a stepper motor (Oriental motor, PKP564FMN24A) and rotary union (DSTI, LT-2141) to rotate airflow and odor stimuli around the fly, while the fly remained stationary in the center of the arena (Currier et al., 2020). In these experiments we rotate the stimulus to the same 5 directions we could present with the manifold. Airspeed was 40 cm/s and presented through 3-way solenoid valves (Lee Company, LHDA1233115HA). We used a hotwire anemometer (Dantec Dynamics) to verify that airspeed was equivalent across directions and constant through the wind and odor phases of the stimulus. Odorant (apple cider vinegar, 10%) was diluted in distilled water on the day of the experiment. Each trial consisted of 5-10s without stimuli, 10s of wind alone, 10s of odorized wind, 10s of wind alone and 8-10s of no stimulus following the wind. We randomized the direction from which odor was presented in blocks of 5 trials and completed 5 blocks for each fly (25 total trials). We adjusted imaging position, Z-plane, gain, and power levels after each block as necessary. Of the flies included, we present data from all 25 trials here. MB052B, LH1396, 65C03-GAL4, 21D07-GAL4, vFB split, and VT062617-GAL4 were imaged with the stimulus manifold, while MB077B, MB082C and 12D12-GAL4 were imaged with the rotating stimulus setup. Stimuli were controlled through custom MATLAB and Python scripts.

Analysis of calcium data was performed as in Currier et al. 2020. We used the CalmAn MATLAB package, ImageJ, and custom MATLAB scripts to align and analyze data. We used the CalmAn package (Giovannucci et al. 2019) to implement the NoRMCorre rigid motion correction algorithm (Pnevmatikakis et al. 2017) on the red (tdTOM) time series and applied the same shifts to the green (GCaMP6f) times series. We drew regions of interest (ROIs) by hand on maximum intensity projections of the tdTom time series for the first trial. ROIs were applied to all trials, and had their position manually adjusted using imageJ if significant drift occurred between trials. ROIs were drawn around the following regions: for LH1396, the ROI was placed around the dendritic processes in the LH, for MB052B, MB082C and MB077B the ROIs were placed around the putative axonal processes in the protocerebrum. ROI location and imaging region was selected based on pilot experiments recording from different planes and ROIs. For tangential FB inputs (FB5AB, 65C03-GAL4, 12D12-GAL4 and vFB split) we imaged from the FB and placed our ROI around the layer innervated by each GAL4 line. We report imaging quantifications across the entire layer in the figures. In tangential inputs we did not observe obvious direction specific responses in different anatomical locations of the output layer. For VT062617-GAL4 imaging of hΔC neurons, we drew ROIs across 8 putative columns of the FB based on the glomerular structure that was observable in the tdTom signal. In some cases, the true number of columns was unclear and depended on the exact plane of imaging and positioning of the fly’s head. ImageJ ROIs were imported into MATLAB using ReadImageJROI (Muir et al. 2014). We calculated ΔF/F for the GCaMP6f time series by dividing the time series by the average fluorescence of the baseline period (first 5s of the trial, excluding the first sample due to shutter lag). For main text figures we present the average ΔF/F signal for individual flies. Mean traces were calculated by resampling to 5 samples per second if frame rate varied between experiments. Supplemental figure heat maps were normalized to maximum response across all trials within an individual fly.

### ELECTROPHYSIOLOGY

We performed whole cell patch clamp recordings as described previously (Suver et al. 2019, Currier et al. 2020). Mounting and dissection were similar to that described above for calcium imaging, except that we used hot wax rather than UV glue to fix the fly in place. In addition, we removed the sheath covering the brain using collagenase (5% in extracellular saline, Worthington Biochemical Corporation Collagenase Type 4) under positive pressure applied with a fine tipped electrode (5-10uM diameter). Cell bodies of interest were visualized with 10x cytoplasmic GFP using an LED source (Cairn Research MONOLED) and filter cube (U-N19002 AT-GFP/F LP C164404). Brains were visualized under 40x magnification (Olympus, LUMPLFLN40XW) using a camera (Dage-MTI, IR-1000) and an LCD monitor (Samsung, SMT-1734). Cell bodies were cleaned using external saline and positive pressure, as well as light negative pressure to remove cell bodies near our cell of interest.

For whole-cell patch clamp recordings, we pulled 6 to 10 M-Ohm glass pipettes made of thick-walled glass (World Precision Instruments 1B150F-3) using a Sutter Instruments P-1000 puller. Pipettes were polished using a pressurized micro-forge (Scientific Instruments, CPM-2). Our intracellular solution contained 140 mM KOH, 140 mM aspartic acid, 10 mM HEPES, 1 mM EGTA, 1 mM KCl, 4 mM MgATP, 0.5 mM Na3GFP, and 13 mM biocytin hydrazide (for visualization of neural processes). Current and voltage signals were amplified using either an A-M systems Model 2400 amplifier or a Molecular Devices Multiclamp 700B. Recordings acquired with the A-M systems amplifier were paired with additional preamplification using a Brownlee Precision 410 preamplifier. We controlled stimuli and hardware using custom MATLAB and Arduino software scripts. All electrophysiological recordings were acquired at 10kHz.

Stimuli were delivered using an olfactometer similar to the one described in Nagel and Wilson 2016. Charcoal-filtered air was passed through a flowmeter (Cole-Parmer, 0.3 L/min), and then split into two airstreams that passed over either odorant (10% vinegar) or water. These two airstreams were then passed through two three-way solenoid valves (Lee company, LFAA1201610H) that allowed a signal to switch which airstream (odor or water) was directed into the main airflow. Main airflow (1L/min) was delivered to the fly through a Teflon tube (4mm outer diameter, 2.5mm inner diameter). The Teflon tube was positioned <1mm from the head of the fly using a micro manipulator for each experiment using two cameras (Unibrain). The airflow delivery system was positioned on the right side of the fly, thus cells on the fly’s right were ipsilateral while those on the left were contralateral. Pulses of 2s, 10s or 20s of odor were presented to the fly. Only 10s of vinegar is displayed here.

We used custom MATLAB scripts to analyze electrophysiology data. To analyze membrane potential, we applied a 2.5Hz Butterworth filter to remove spikes. We averaged the baseline period of of 2s and subtracted this from the average time course for each fly for presentation purposes. We report the difference between the baseline period (wind only) and the average during the first 4s of the odor period. The resting potential varied between -34.6 mV and -25.2 mV for MBONs α’3. The average resting potential was -30.8mV. We recorded from a total of 12 MBONs, 6 on each side that met our criteria for quality of recording based on input to access ratio great than 5:1.

### IMMUNOHISTOCHEMISTRY

We performed immunohistochemistry as in previous reports (Suver et al. 2019, Currier et al. 2020). We fixed brains for 15 minutes in 4% paraformaldehyde (in 1X phosphate buffered saline, PBS). Next, we washed the brain three times in PBS and stored at 4°C until antibody staining (immediately or within 2 weeks). We incubated brains in a blocking solution containing 5% normal goat serum dissolved in PBST (1x PBS with 0.2% Triton-X) for 20-60 minutes. Brains were incubated at room temperature in a solution of primary antibodies (see below for exact components). We then washed brains three times in PBST and incubated brains in a secondary antibody solution at room temperature for 24h. We washed brains three times in PBST and then stored in PBS at 4°C until imaging. To mount brains for imaging we placed brains in vectashield (Vector Labs H-1000) and sealed with coverslips and nail polish. We imaged brains at 20x magnification of a Zeiss LSM 800 confocal microscope with a 20x objective (Zeiss W Plan-Apochromat 20x/1.0 DIC CG 0.17 M27 75mm). All brains were imaged at 1-1.25uM depth resolution. Final images are presented as maximum Z projections over relevant depths.

To visualize the expression of *Chrimson-mVenus* in driver lines used for optogenetic activation experiments, we dissected brains of females from the same cross as experimental males. We visualized 1-3 brains for each genotype (data not shown) to assess the breadth of expression under the Chrimson effector.

We used the following antibody mixes for the experiments listed: Chrimson/neurotransmitter stains: primary: chicken anti-GFP (1:50) & mouse anti-nc82 (1:50); secondary: anti-chicken Alexa488 (1:250) & anti-mouse Alexa 633 (1:250). Electrophysiology stains: primary: chicken anti-GFP (1:50) & mouse anti-nc82 (1:50); secondary: anti-chicken Alexa488 (1:250), anti-mouse Alexa 633 (1:250) and Alexa568-conjugated streptavidin (1:1000). Trans-tango and CLIN stains: primary: chicken anti-GFP (1:50), mouse anti-nc82 (1:50), & rabbit anti-dsRed (1:500); secondary: anti-chicken Alexa 488 (1:250), anti-mouse Alexa 633 (1:250) and anti-rabbit Alexa 568 (1:250). GABA stains: chicken anti-GFP (1:50), mouse anti-nc82 (1:50) & rabbit anti-GABA (1:100); anti chicken Alexa488 (1:250), anti-mouse Alexa 633 (1:250) & anti-rabbit Alexa 568 (1:250).

### CONNECTOMIC ANALYSIS

Data from the hemibrain connectome (Scheffer et al. 2020) were interpreted using neuprint explorer (neuprint.janelia.org, version d2a8f5785d73421096f7cdc09ad585e5). Further analysis and visualization were completed using custom MATLAB and Python scripts. For the analysis shown in Fig. 5B, the location of FB5AB synapses onto hΔC was determined by their x position. No filtering or constraints were applied in synapse counts for this panel. For the analysis in Fig. 4B, we counted all FB tangential neurons two synapses downstream of the following projection neurons: VM7d_adPN, VM7v_adPN, DM1_lPN, DM4_adPN, DM4_vPN, VA2_adPN, DP1l_adPN, DP1l_vPN, DL2d_adPN, DL2d_vPN, DL2v_adPN, DC4_adPN, DC4_vPN, DP1m_adPN, DP1m_vPN in which the intermediate neurons contained LH (they were lateral horn neurons) and where synaptic weights exceeded a weight of 3.

### MODELING

The FB steering circuit model implemented in this study contains three types of inputs - heading, wind, and odor; two intermediate layers corresponding to hΔC and mutual inhibition local neurons; and an output layer corresponding to left PFL3, right PFL3, and PFL2 neurons, which modulate angular velocity and walking speed. As a general layout, heading inputs are sent directly to PFL3 and PFL2 neurons, while odor and wind inputs are integrated by hΔC neurons, processed by the mutual inhibition layer, and then integrated by PFL3 and PFL2 neurons. The relative activity of left and right PFL3 neurons are used to modulate angular velocity, while the activity of PFL2 neurons is used to modulate walking speed.

All modeling was performed in Matlab.

#### Input layer: heading

Heading input into PFL3 and PFL2 neurons were simulated by phase-shifting heading-tuned one-cycle sinusoids:

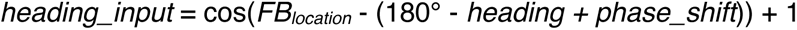

Where *FB_location_* refers to the spatial location within the FB (0° = left FB, 360° = right FB), *heading* refers to the heading direction of the fly, and *phase_shift* refers to the anatomical shift introduced by PFL neurons entering the FB. The phase shift parameters were taken directly from the connectome (Hulse et al. 2021) and were defined as the following: phase_shift_left_PFL3_ = -90°, phase_shift_right_PFL3_ = +90°, phase_shift_PFL2_ = +180.

#### Input layer: wind

Wind-tuned bumps of activity in hΔC neurons were simulated using one-cycle sinusoids that depended on the encoding strategy. For allocentric representation (Figure 7), the bump of activity in hΔC neurons follows the allocentric direction of wind and was defined by the following:

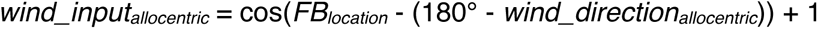

Where *wind_direction* refers to the allocentric wind direction. For frontal representation (Fig. S7) hΔC wind activity was constructed by phase shifting and summing the activity of a left and right population of wind-sensitive PFNs (Currier et al., 2020), each displaying a heading-tuned bump scaled by wind direction:

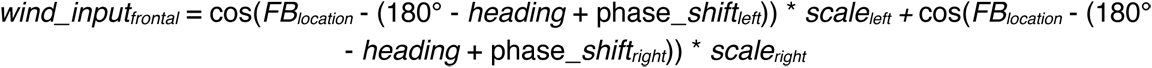

Where phase_*shift* refers to the anatomical shift introduced by PFN neurons entering the FB, and *scale* refers to the wind-tuned scaling of bump amplitude. The phase shifts differed for the two populations of PFNs and were defined as: phase_shift_left_ = +45°, phase_shift_right_ = -45 (Hulse et al., 2021). The scaling corresponded to the wind tuning of the PFNs, which differed between the left and right PFN populations (Currier et al., 2020):

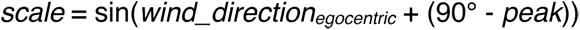

Where *wind_direction_egocentric_* is the egocentric wind direction and peak is the egocentric wind direction that maximally activates the PFN cell type (peak_left_ = -45°, peak_right_ = +45°).

#### Intermediate layer 1: hΔC neurons

The total activity of hΔC neurons was calculated as a summation of wind input and optogenetic input:

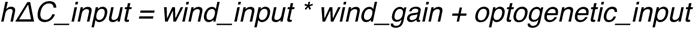

Where *wind_gain* is a binary variable equal to 1 only if odor is present and *optogenetic input* is a vector of sparse or broad optogenetic patterns. Sparse optogenetic patterns were generated by creating a 20-element vector, with each element stochastically set to 0 or 2.3 at a 15% probability, and broad optogenetic patterns were generated by creating a 20-element vector with every value set to the same value (low light = 0.75, high light = 2.3). These optogenetic values were chosen to best match simulated trajectories with real behavior.

The output of the hΔC population was calculated by phase-shifting the hΔC activity by 180°. To simplify the pathways connecting hΔC neurons with mutual inhibition circuits, PFL3 neurons, and PFL2 neurons, which all involve at least one additional cell type, the output of hΔC neurons was transformed from 20-neuron space into 8-column space and then thresholded using a sigmoid activation function:

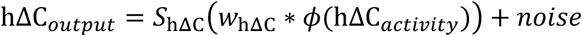

Where *S_hΔC_* is the hΔC sigmoid activation function with slope *k_hΔC_* and threshold *θ_hΔC_*, *w_hΔC_* is the feedforward transformation matrix, *ϕ* is the 180° phase shift function, and noise is exponentially filtered white noise. The matrix used for the space transformation was inspired by the connectivity between local neurons in the fan-shaped body. To construct this matrix, each neuron’s output was set as the size of one column, such that there were 20 overlapping column-sized outputs. The matrix element for column_i_ and neuron_j_ was then set as the amount of overlap between column_i_ and the output of neuron_j_, and the total input to column_i_ from all neurons was normalized to equal 1. The exact structure of *w_hΔC_* was not critical, as long as the matrix was normalized and maintained the approximate location of wind sinusoids in FB space. The filtered white noise was defined by the following equation (Shpiro et al. 2007):

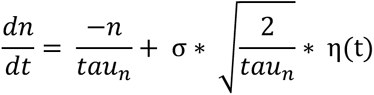

Where *n* is the noise variable, *tau_n_* is the noise time constant, σ*_n_* is the noise standard deviation, and η(t) is white noise with zero mean and unit variance.

#### Intermediate layer 2: mutual inhibition local neurons

hΔC columnar output was then relayed into an inhibitory recurrent network, consisting of 8 inhibitory neurons with mutual connections between neurons located in opposite columns. Neurons in this network also contained a slow adaptation parameter, a, that curtailed input when neural activity was high. This system was described using the following differential equations:

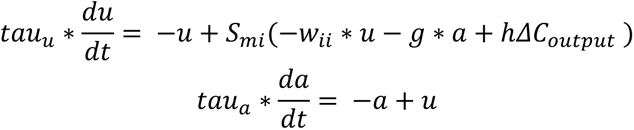

Where *u* is an 8-element vector corresponding to the activity of each inhibitory neuron, *a* is an 8-element vector corresponding to the adaptation of each inhibitory neuron, *w_ii_* is the recurrent weight matrix connecting the inhibitory neurons, *g* is the adaptation weight, *hΔC_output_* is the columnar output of the hΔC neurons, *tau_u_* is the time constant for neural activity, *tau_a_* is the time constant for adaptation, and *S_mi_* is the mutual inhibition sigmoid activation function with slope *k_mi_* and threshold *θ_mi_*. The equations and parameters used to model the mutual inhibition circuit were taken from Shpiro et al., 2007.

The output of the mutual inhibition population was calculated by phase-shifting *u* by 180°. This output was subtracted from the hΔC output to create the total local output sent to PFL3 and PFL2 neurons.

#### Output layer: PFL2 and PFL3 neurons

The activity of left PFL3, right PFL3, and PFL2 neurons was calculated by summing the excitatory output of hΔC neurons, inhibitory output of the mutual inhibition circuit, and phase-shifted excitatory output of the heading system, and then converting this input into activity using the following dynamic system:

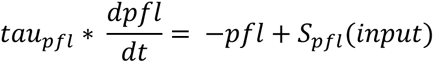

Where *tau_pfl_* is the time constant for PFL neurons and *S_pfl_* is the PFL sigmoid activation function with slope *k_pfl_* and threshold *θ_pfl_*.

#### Output layer: navigation modulation

Turning was calculated by comparing the activity of right PFL3 neurons and left PFL3 neurons and adding a noise term:

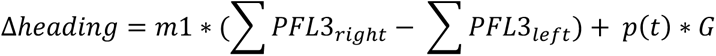

Where *m* is the coupling constant between PFL3 activity and turning, *p*(*t*) is a binary Poisson variable with λ rate, and *G* are angular velocity values drawn from a Gaussian distribution with zero mean and σ_turn_ variance. The random turn equations and parameters were taken from Álvarez-Salvado et al., 2018, while the coupling constant was selected to best match behavioral trajectories.

Groundspeed was calculated by summing the activity of PFL2 neurons according to the following equation:

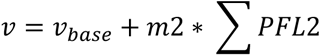

Where *v_base_* is the baseline speed and m*2* is the coupling constant between PFL2 activity and speed. The baseline speed was taken from Álvarez-Salvado et al., 2018, while the coupling constant was selected to best match behavioral trajectories.

#### Output layer: simulated trajectories

The model simulates heading and groundspeed over time in response to wind/odor or optogenetic stimuli. These values were converted into movement in x and y using:

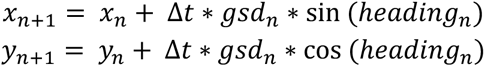

Where, Δ*t* is the time step between data points, *gsd_n_* is the current groundspeed, and *heading_n_* is the current heading.

#### Other

The sigmoid activation function used for hΔC neurons, the mutual inhibition circuit, and PFL neurons was the following:

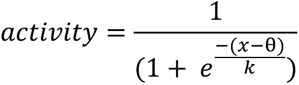

Where *x* is the input, *θ* is the activation threshold, and *k* is the activation slope.

All dynamic systems were computationally approximated using Euler’s method of integration with a time step of 0.05.

Note: many parameters were manually fit to match simulation data with behavioral trajectories. While not explored in this study, many of these parameters (e.g., activation thresholds) are flexible and can be adjusted together to keep the steering circuit functioning across a range of sinusoid amplitudes.

**Table #.**
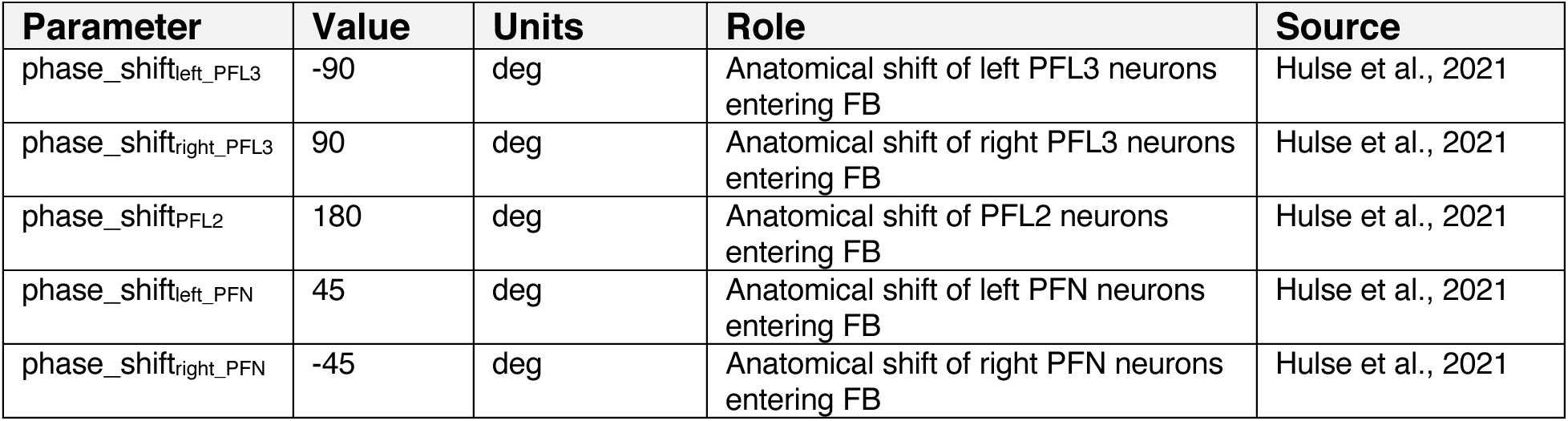
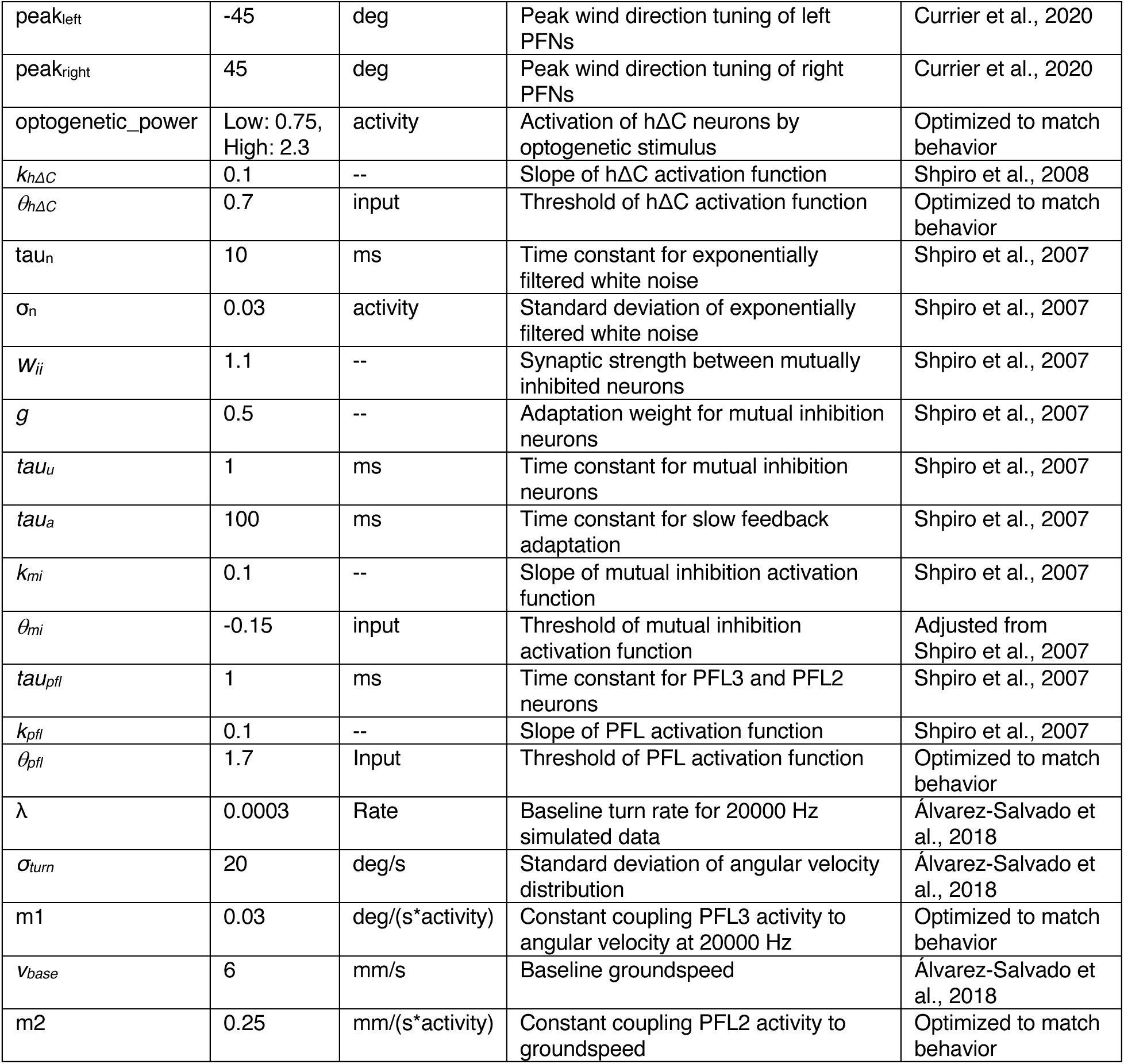
Values of steering circuit model parameters with their units, roles, and sources listed.

### QUANTIFICATION AND STATISTICAL ANALYSIS

#### Behavior

Based on previous analysis of behavioral data (Alvarez-Salvado et al. 2018, Suver et al. 2019) we assumed behavioral data was not normally distributed. We applied non-parametric statistics to compare results and corrected for multiple comparisons using the Bonferroni method. We used the two-sided Wilcoxon signed rank test (MATLAB signrank) to compare the average baseline value of each parameter to the average of the parameter over a window of interest. Between genotype comparisons (genetic silencing experiments) were made between baseline subtracted parameter values using the two-sided Mann-Whitney U test (MATLAB ranksum). For behavioral time courses we display standard error around the mean. For summary plots we present standard deviation around the mean. Bonferroni corrections were applied based on the number of genotypes tested labelling similar neuron types (i.e. 6 in Fig. 1D, 21 for all dorsal FB inputs in Fig. 3B,C) or for the number of comparisons made to control in genetic silencing experiments (3 in Fig. 1F).

#### Imaging and physiology

We used parametric tests for calcium imaging and electrophysiological data. When testing for significant differences in odor responses across directions we averaged individual trials across flies, and performed a one-way ANOVA across pooled trials across flies (MATLAB anova1). When assessing differences between wind ON responses and odor responses we calculated mean wind and odor responses in 10s windows following ON. We compared wind and odor periods using a two-tailed paired student’s t-test (MATLAB ttest, Fig. S4E, Fig. S5A). Similarly, we performed two-tailed paired student’s t-test when comparing wind and odor responses in MBON electrophysiology experiments and an two-tailed unpaired student’s t-test (MATLAB ttest2) when comparing odor responses between ipsilateral and contralateral MBONs. To assess the decay (Fig 4D) of the fluorescence response to odor over time, we computed the average response to the odor presentation period across flies for five trial blocks, including one trial of each direction. We averaged the response across all flies imaged, and normalized by the mean response in trial block one. Shaded regions depict standard error across flies around the mean.

#### Classification training

We trained classification tree models on calcium response data to assess if an observer could correctly identify if wind was coming from the left, right or head on relative to the fly. We pooled the data for 45 and 90 degrees left and right for this analysis. We trained these models on the activity during odor ON (5s, Fig. 2F, Fig. 4D) and during wind ON and OFF (5s, Supp Fig 2H, Supp Fig 4E). Model were trained using two predictors, the fly identity, and the Z-score of the calcium response during the specified window. We took the Z-score of the response to standardize the data between electrophysiology firing rates (PFNa) and calcium responses (all other neurons). As we recorded both left and right LNa neurons simultaneously we asked how well the difference between the activity of the two performed. We trained classification trees (Matlab’s fitctree) to identify wind direction and tested model performance using 10-fold cross validation error (Matlab’s KfoldLoss). We then compared this error to classification trees that were trained on the same data with shuffled labels, with 50 random iterations of shuffled labels.

We also designed classifier to assess if an observer could distinguish between wind ON activity and odor ON activity. We used the same parameters in the direction classifier but pooled all data across all wind directions. We performed the same analysis to assess if an observer could distinguish between baseline activity and odor ON activity in Supp. Fig 2F and Supp. Fig 4F.

### KEY RESOURCES TABLE

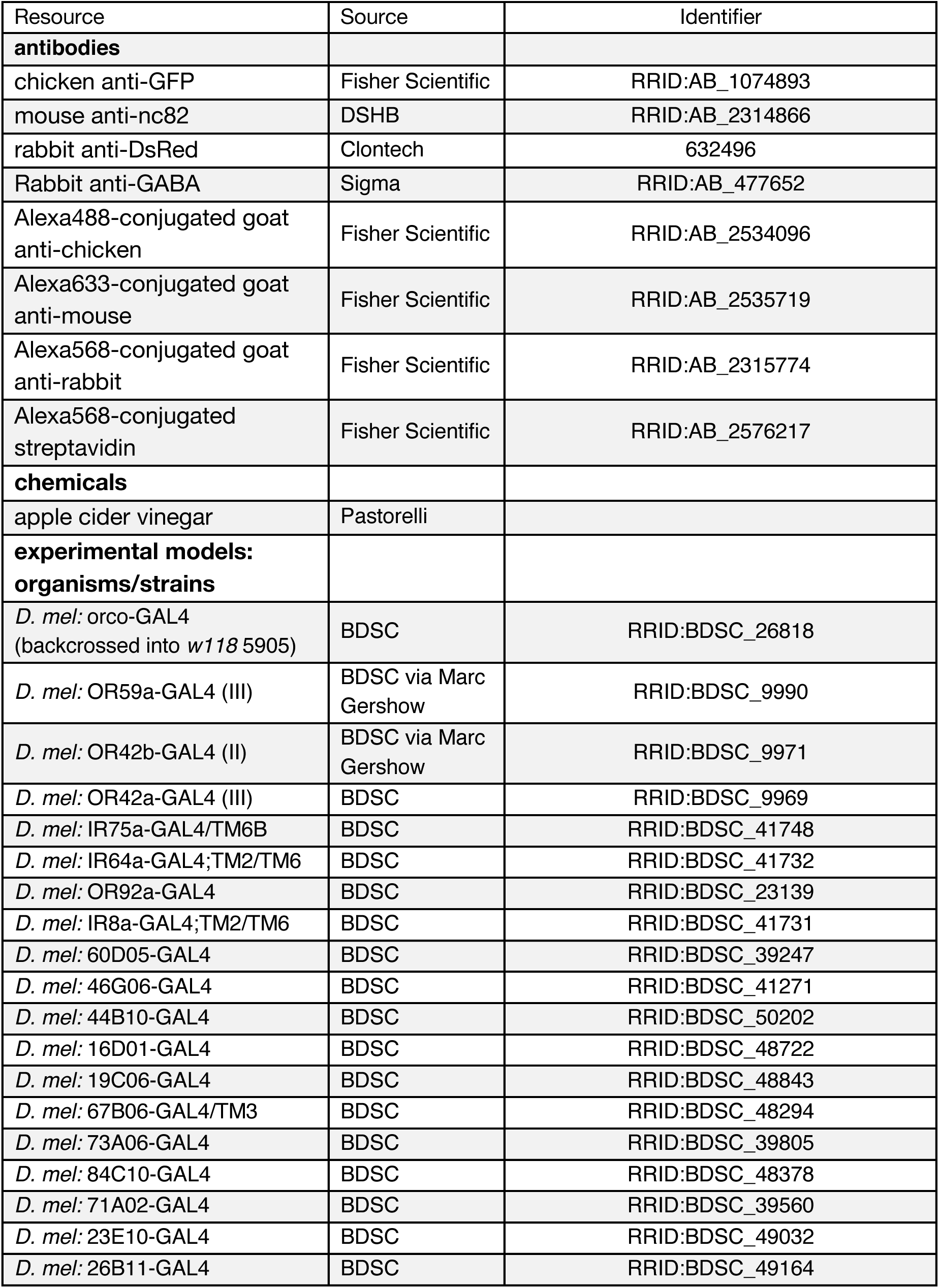
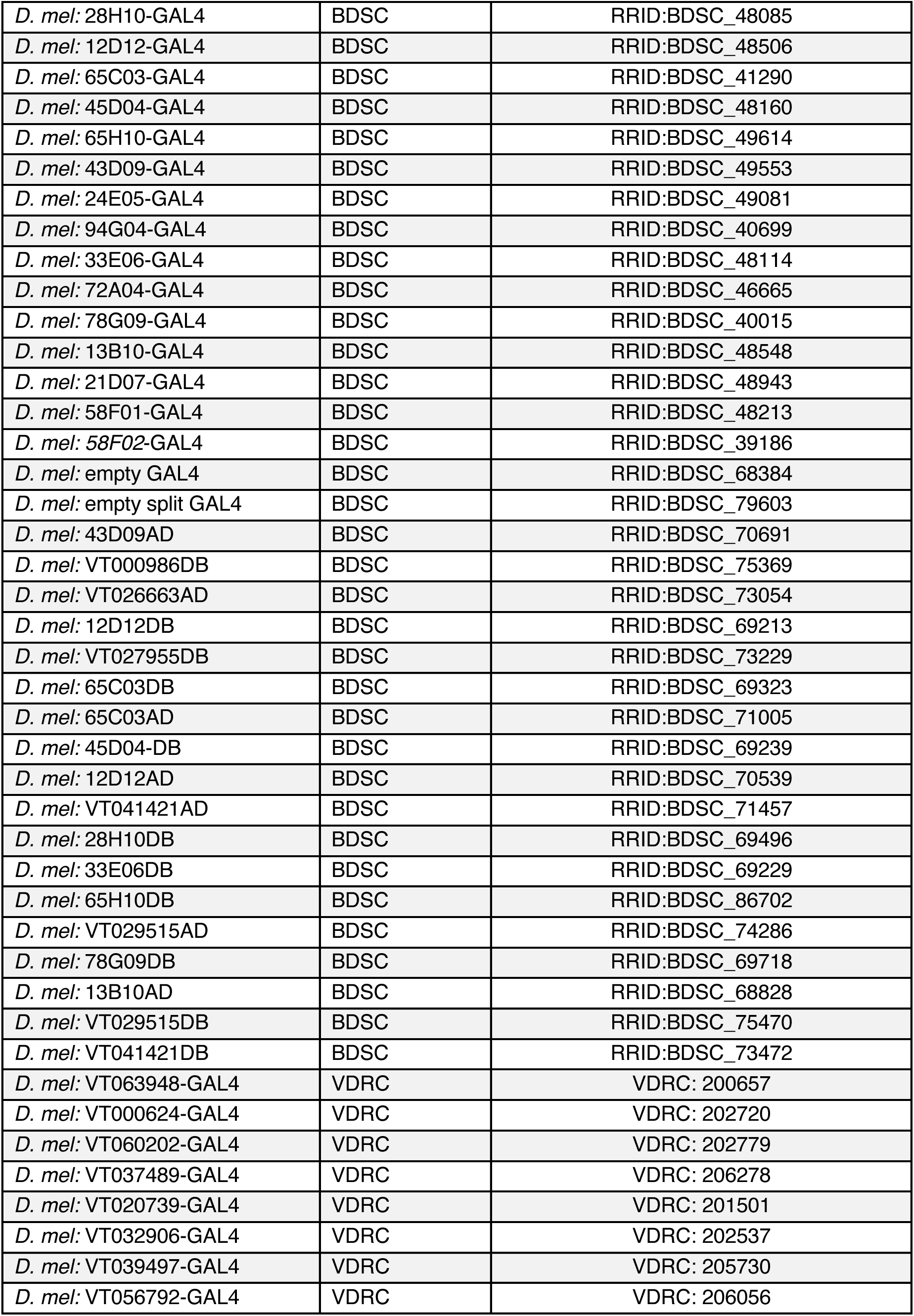
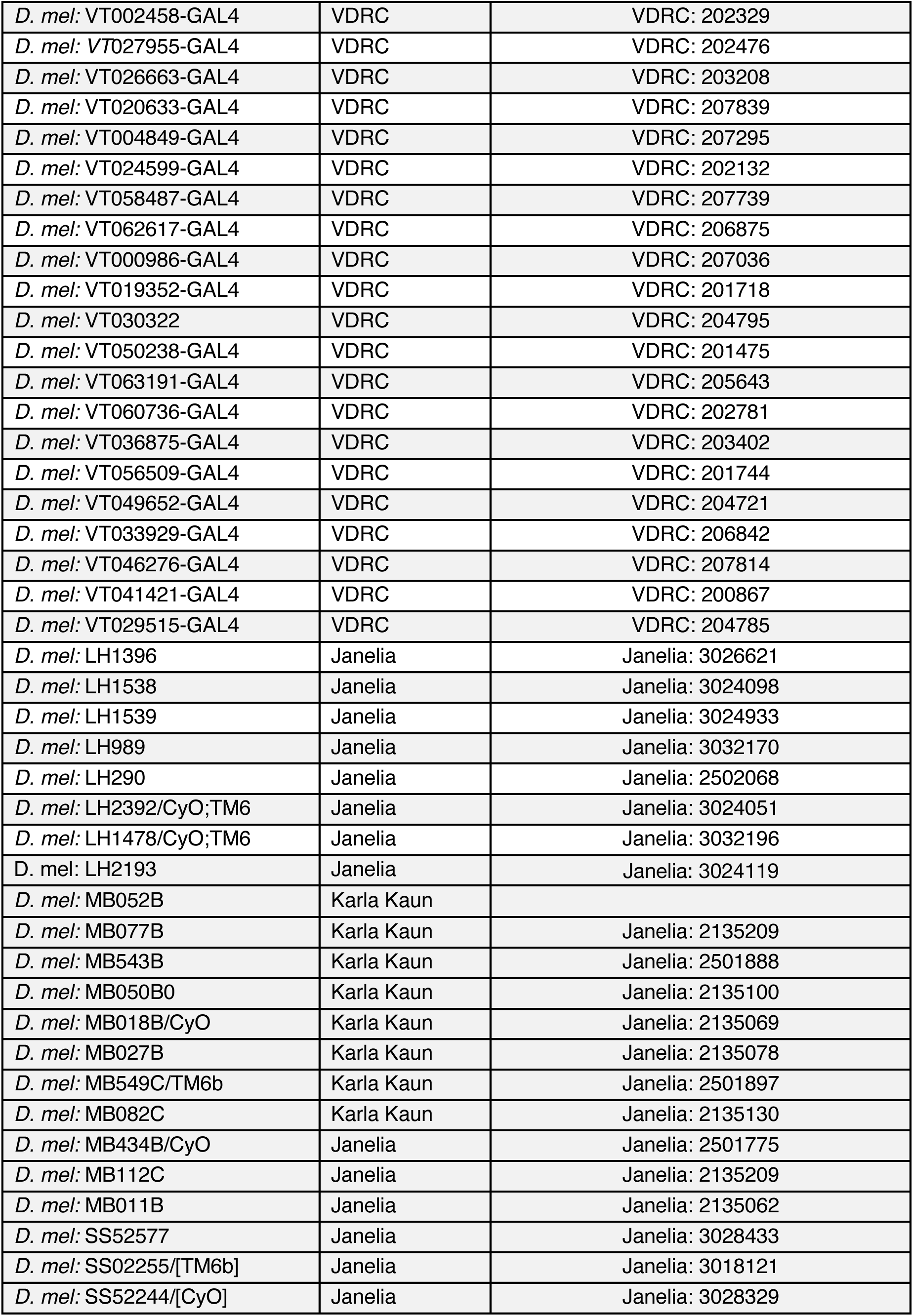
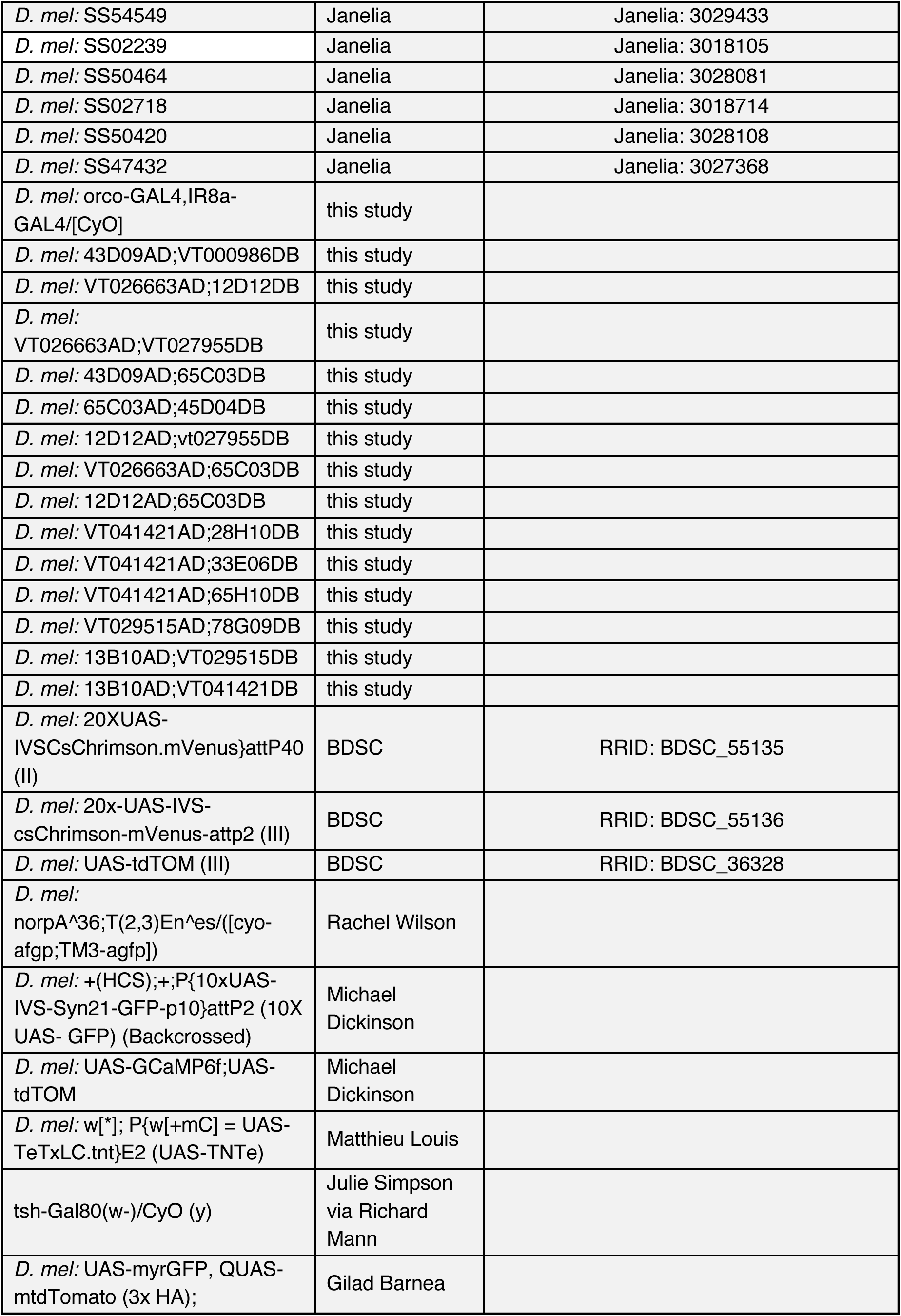
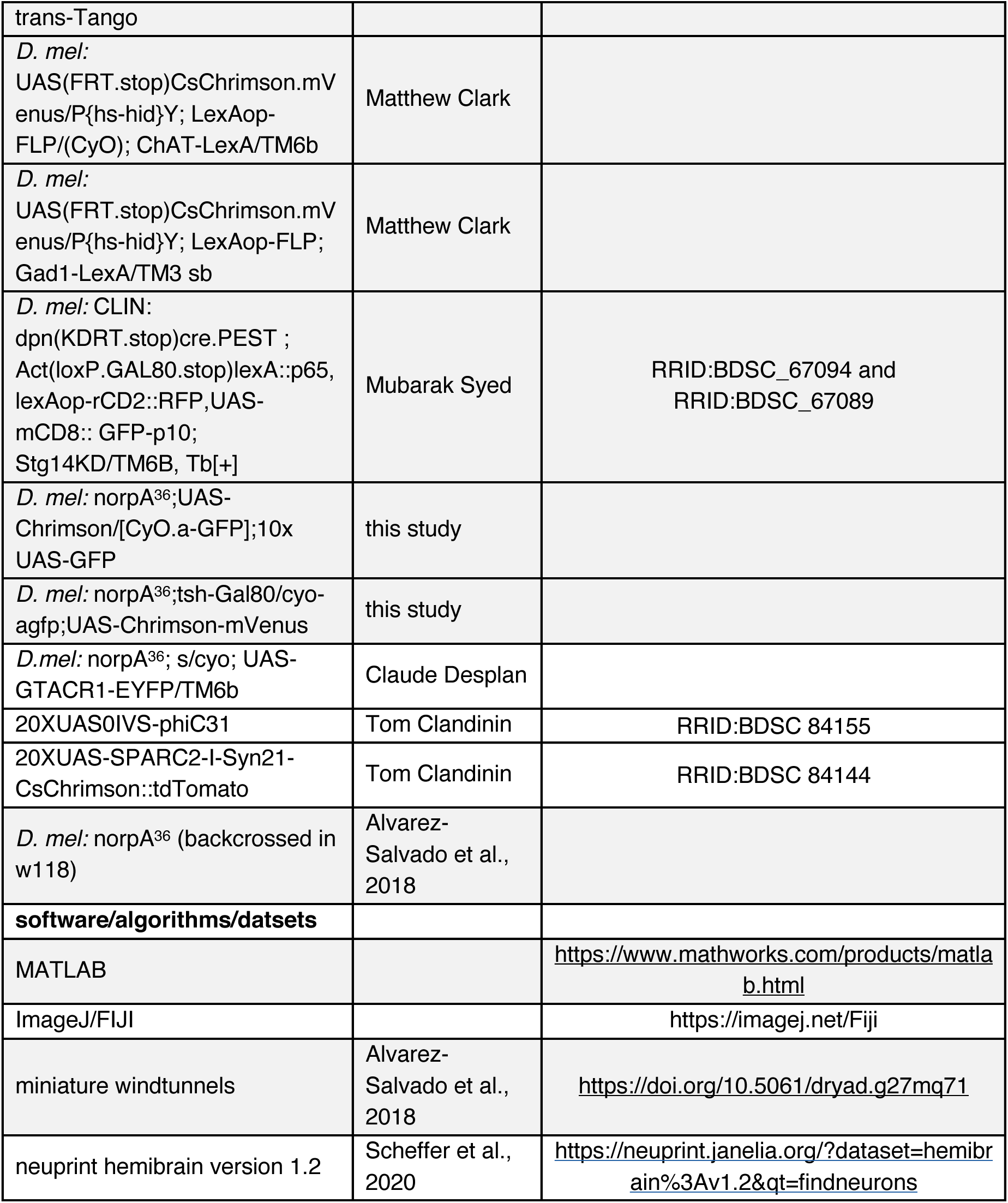

## Data Availability

All data generated during the study will be available on Dryad at accession number 149533.

## Code availability

All original code is available on Github at https://github.com/nagellab/Mathesonetal2021. Any additional information required to reanalyze the data reported in this paper is available from the Lead Contact upon request.

## Materials availability

No new transgenes were created for this study. Transgenic stocks are available on request from the Lead Contact.

## Acknowledgements

The authors would like to thank Marc Gershow, Karla Kaun, Tom Clandinin, Calude Desplan, Michael Dickinson, Michael Reiser, Matthew Clark, Matthieu Louis, Richard Mann, Gilad Barnea, Rachel Wilson, Tzumin Lee, Janelia Flylight, and the Bloomington and Vienna Drosophila Resource Centers for fly stocks. We thank Michael Long, Elizabeth Hong, Floris van Breugel, David Schoppik, and members of the Nagel and Schoppik labs for helpful input on the manuscript. This work was supported by R01DC017979, RF1NS127129, NSF Ideaslab (IOS-1555933) and NSF Neuronex grants, as well as a McKnight Scholar Award to K.I.N. M.H.S was supported by an NSF CAREER award IOS-204720. T.A.C was supported by a Dean’s Dissertation Fellowship from NYU. A.M.M. was supported by a Diversity Supplement to R01DC017979.

## Author contributions

K.I.N, A.M.M.M, and A.J.L. conceived the project. A.M.M.M performed the majority of behavioral, imaging, electrophysiology, and anatomy experiments, and generated new genetic stocks. A.J.L. performed SPARC experiments with A.M.M., performed connectomic analysis, and developed the computational model with input from K.I.N. and A.M.M.M. A.M.L. contributed to behavioral experiments. T.A.C. performed a subset of electrophysiology experiments. M.H.S. provided the genetic strategy to isolate CX neurons. K.I.N, A.M.M.M, and A.J.L wrote the paper with input from the other authors.

## Competing Interests

The authors declare no competing interests.

---

**Fig. S1:**
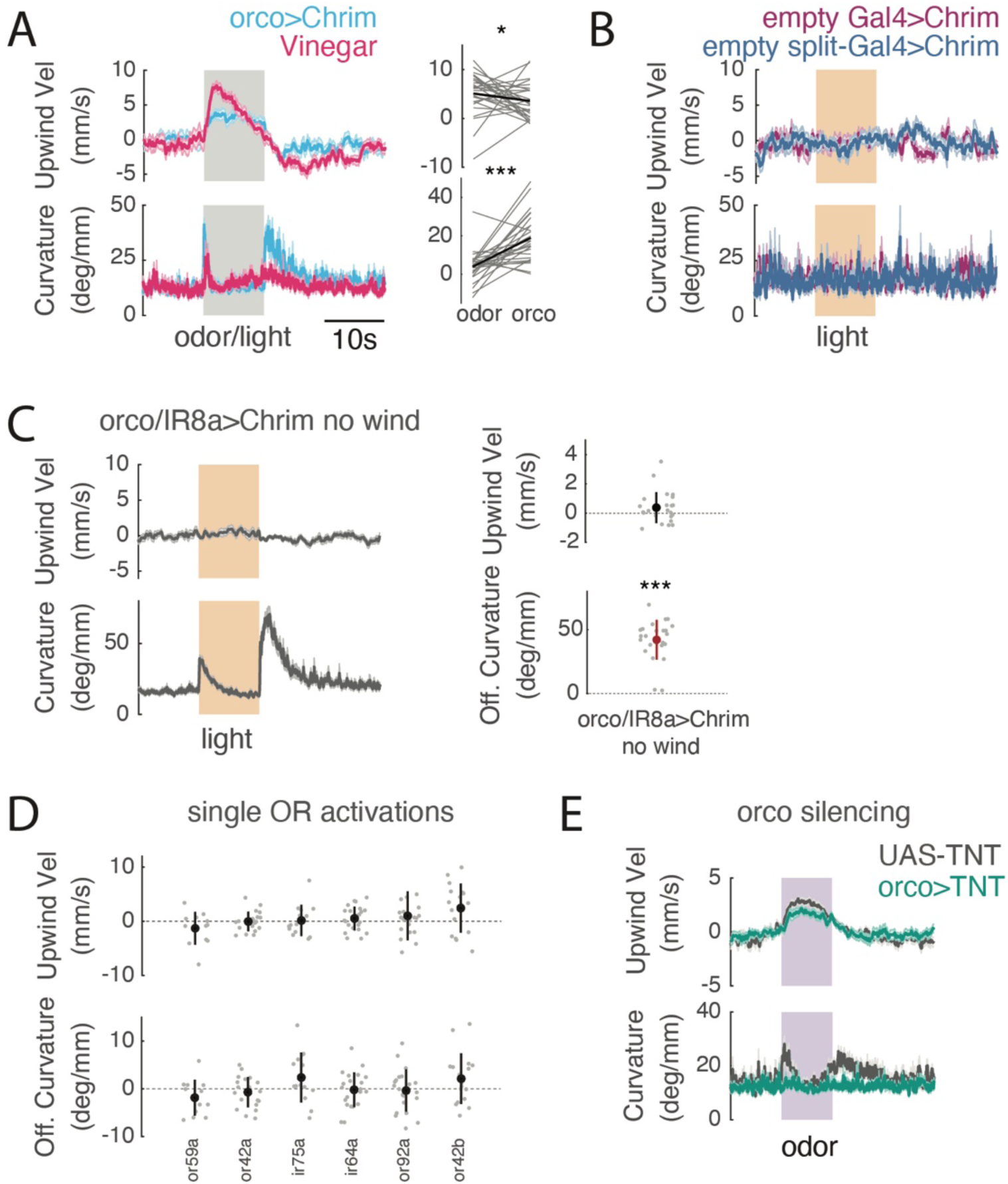
Additional data on ORN encoding of olfactory navigation behavior. **A)** Left: Upwind velocity and curvature time courses (mean±SEM) in the same flies (N = 26 flies) exposed to both 1% vinegar (pink) and optogenetic stimulation of orco+ ORNs (blue) in interleaved trials. Right: Comparison of average upwind velocity (0-5s after stimulus ON) and OFF curvature (0-2s after stimulus OFF) for each fly. Within each fly, vinegar evoked greater upwind velocity than orco stimulation (Wilcoxon signed rank test, p=0.0319) but weaker OFF curvature than orco stimulation (Wilcoxon sign rank test, p=1.3216e-04). **B)** Upwind velocity and curvature time courses (mean±SEM) for control flies: empty-GAL4>Chrimson (purple, N= 19) and empty split-GAL4>Chrimson (blue, N= 14). No significant change in either parameter during stimulation (quantified in Figure 1D). **C)** Right: Upwind velocity and curvature time courses (mean±SEM) for orco/IR8a>Chrimson flies in the absence of wind (N= 24 flies, see also Alvarez-Salvado et al., 2018 and Suver et al., 2019). Left: quantification as in Fig. 1D. No increase in upwind velocity (Wilcoxon signed rank test, p= 0.0191), but OFF curvature increases significantly increases after stimulation (Wilcoxon signed rank test, p= 1.8215e-05). **D)** Upwind velocity and OFF curvature (average change from baseline for single flies, mean ±STD overlaid) for each single ORN that responds to vinegar (Jung et al., 2015). No significant increase in upwind velocity (Wilcoxon sign rank test, p= 0.2334, 0.7089, 0.9032, 0.3304, 0.3754, 0.0442) or OFF curvature (Wilcoxon sign rank test, p= 0.1763, 0.3317, 0.0785, 0.7151, 0.6639, 0.1488). **E)** Timecourse of upwind velocity and curvature for flies with orco+ ORNs silenced using orco>TNT (N= 25 flies, teal) versus UAS-TNT controls (N= 31 flies, gray). All comparisons for behavioral experiments Bonferroni corrected for multiple comparisons.

**Fig. S2:**
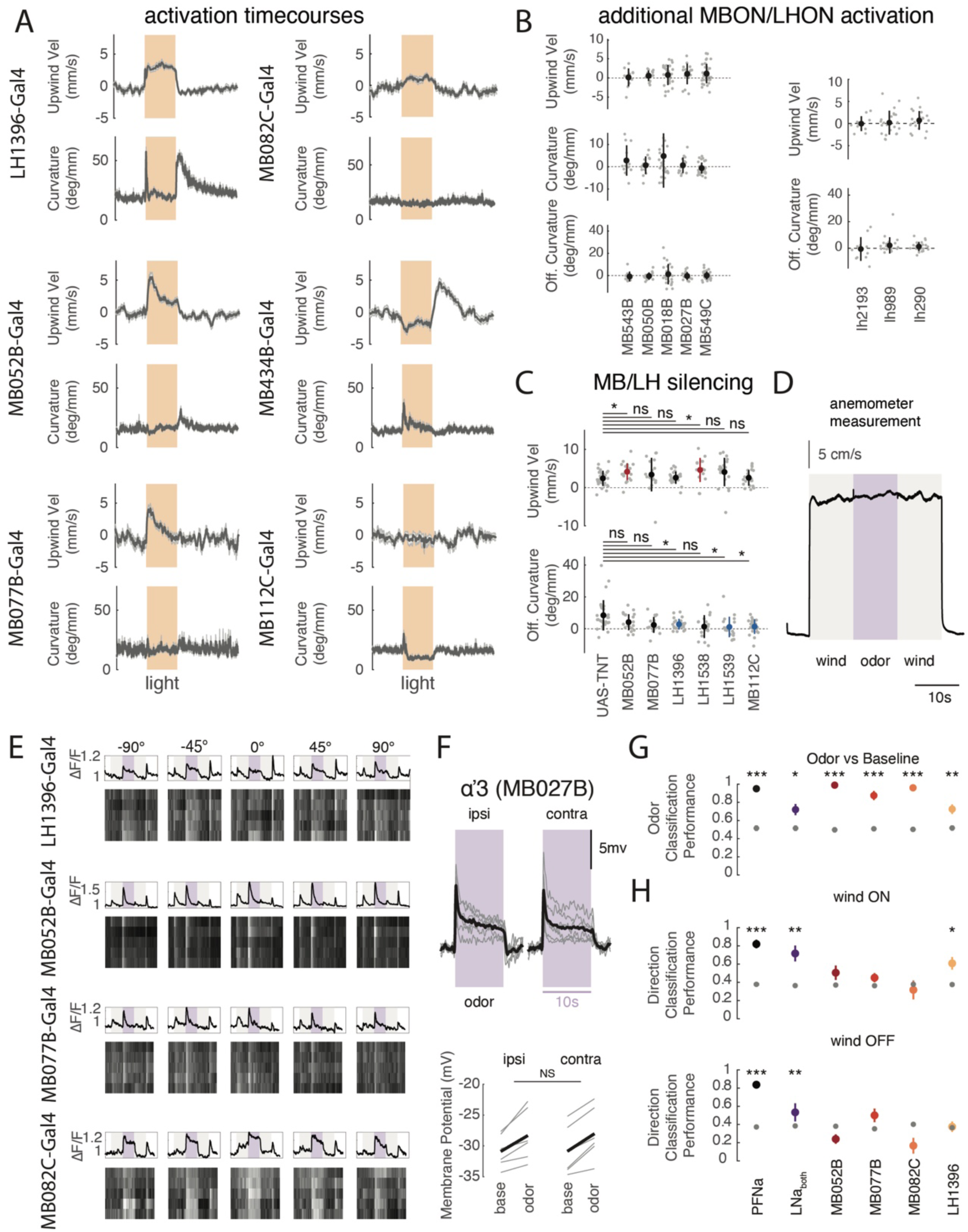
Additional data on LH and MB responses and behavior. **A)** Upwind velocity and curvature time courses (mean±SEM) for LH1396>Chrimson (N =24 flies), MB052B>Chrimson (N =27), MB077B>Chrimson (N=21), MB082C>Chrimson (N=24), MB0434B>Chrimson (N =24), and MB112C>Chrimson (N =29). **B)** Upwind velocity, curvature during stimulus, and OFF curvature for individual MBONs labeled by MB052B (left) and for additional LHONs (right), mean±STD overlaid. MBONs: upwind: MB543B: p=0.5693, MB050B: p=0.1726, MB018B: p=0.1363, MB027B: p=0.1477, MB549C: p=0.0360, curvature: MB110C: p=0.0027, MB543B: p=0.3804, MB083C: p=3.9023e-04, MB050B: p=0.5016, MB018B: p=0.2029, MB027B: p=0.5695, MB549C: p=0.0742 OFF curvature: MB543B: p=0.2061, MB050B: p=0.3910, MB018B: p=0.7533, MB027B: p=0.4380, MB549C: p=0.7807. LHONs: upwind: LH2193: p=0.9658, LH989: p=0.9095, LH290: p=0.0975, OFF curvature: LH2193: p=0.7002, LH989: p=0.1997, LH290: p=0.1443. **C)** Upwind velocity and OFF curvature (quantified as in Fig. 1D) when LHONs and MBONs are silenced. (Mann Whitney U test compared to UAS-TNT control, upwind velocity: MB052B: p=0.0036862, MB077B: p=0.17444, LH1396: p=0.6529, LH1538: p=0.0036449, LH1539: p=0.0090304, MB112C: p=0.83555) OFF curvature: MB052B: p=0.063702, MB077B: p=0.012398, LH1396: p=0.0027408, LH1538: p=0.013958, LH1539: p=0.00070953, MB112C: p=0.0011212) **D)** Average stimulus delivered to the fly from 3 trials from each of the 5 directions as measured by a hotwire anemometer (average across directions). There is no change in windspeed between wind alone and wind + odor. **E)** Single fly examples of calcium responses to wind and odor in LH1396, MB052B, MB077B, and MB082C. Top boxes show average traces for single flies across 5 trials from each direction. Heat maps below depict responses for individual trials. **F)** Membrane potential responses in α’3 MBONs (labeled by MB027B) to 10% vinegar presented from 90° ipsilateral or contralateral to the recorded neuron. Black trace represents mean across flies while gray traces represent individual flies (N=6 cells per hemisphere, each from 1 fly). Odor significantly increases membrane voltage both ipsilaterally and contralaterally (paired student t-test p= 0.0018, 0.0159) and is not different between sides (unpaired student t-test p=0.7561). **G)** Performance of a tree classifier at decoding odor versus baseline activity, trained on 5s of baseline versus first 5s of odor ON. Gray dots represent a classifier trained with the same data and shuffled labels. Student’s t-test, PFNa p= 3.2874e-10, LNa p= 0.0040, MB052B p= 7.9669e-18, MB077B p= 7.7200e-07, MB082C p= 2.3966e-12, LH1396 p= 0.0010. **H)** Performance of tree classifiers at decoding wind direction (left, center, right). Top: classifier trained on the first 5s of wind ON. Student’s t-test, PFNa p= 2.3499e-08 LNa p= 9.0101e-04 , MB052B p= 0.1095 , MB077B p= 0.1122, MB082C p= 0.5650 , LH1396 p= 0.0039. Bottom: trained on the 5s following wind OFF. Student’s t-test, PFNa p= 2.5701e-14, LNa p= 0.1572, MB052B p= 0.0161, MB077B p= 0.0670, MB082C p= 0.0157, LH1396 p= 0.8713. Gray dots represent classifiers trained with the same data and shuffled labels. All error bars on classifiers represent SEM and all comparisons for behavioral experiments and classifier performance are Bonferroni corrected for multiple comparisons.

**Fig. S3:**
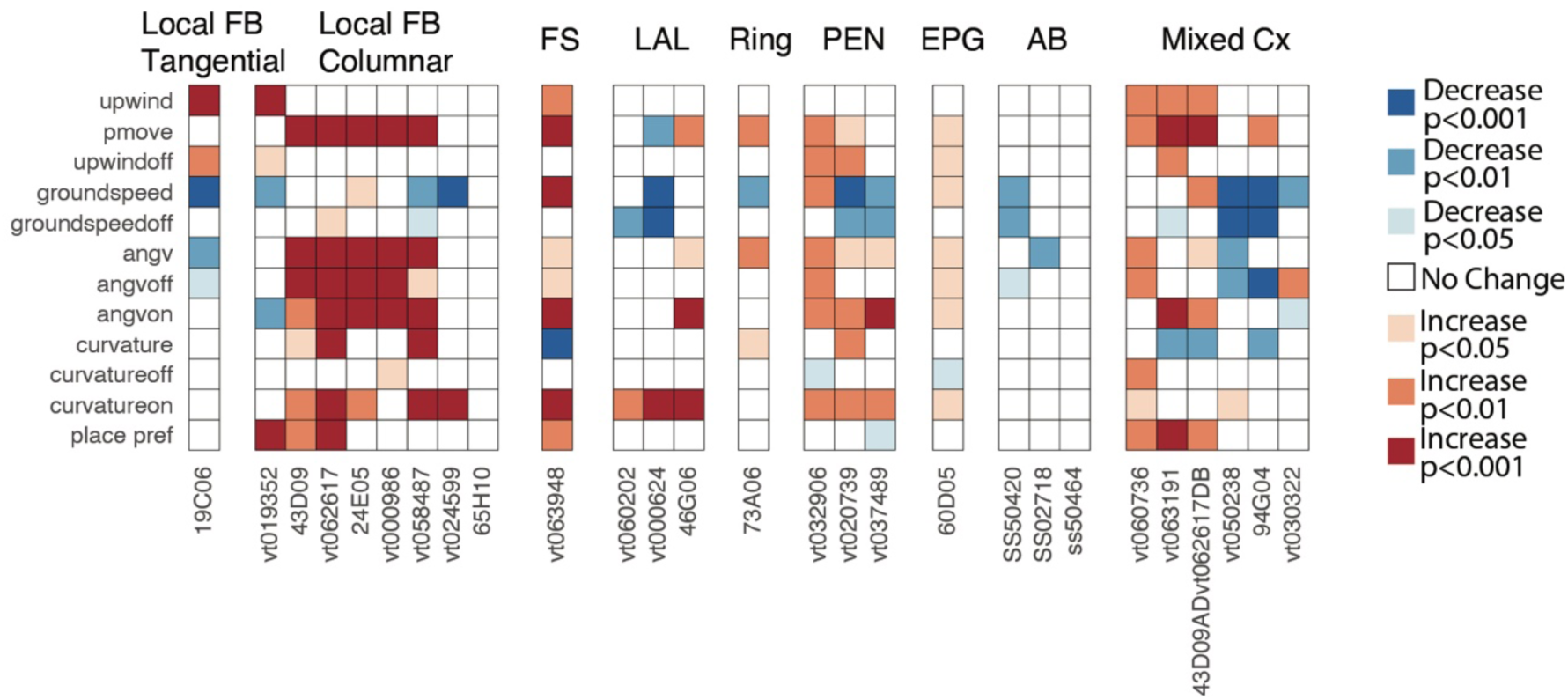
Additional data from optogenetic screen of CX neurons. Optogenetic activation data for additional central complex neuron types (Local FB tangential neurons, Local FB columnar neurons, FS neurons, LAL innervating neurons, Ring neurons, PEN neurons, EPG neurons, AB neurons, and drivers that labelled more than one Cx type prominently (mixed)). One local tangential FB neuron line (19C06-GAL4), one local columnar FB neuron line (VT019352-GAL4), and three mixed Cx lines (VT060736-GAL4, VT063191-GAL4, the split line 43D09-AD; VT062617-DB) and one FS neuron line (VT063948-GAL4) produced significant increases in upwind velocity. No LAL, Ring, PEN, or AB lines produced significant changes in upwind velocity, nor did activation of EPG compass neurons (60D05-GAL4). Several FB local columnar neuron lines drove increases in movement probability (Pmove) and angular velocity (angv): 43D09-GAL4, VT062617-GAL4, 24E05-GAL4, VT000986-GAL4, VT058487-GAL4. See Materials and Methods for calculation of behavioral parameters.

**Fig. S4:**
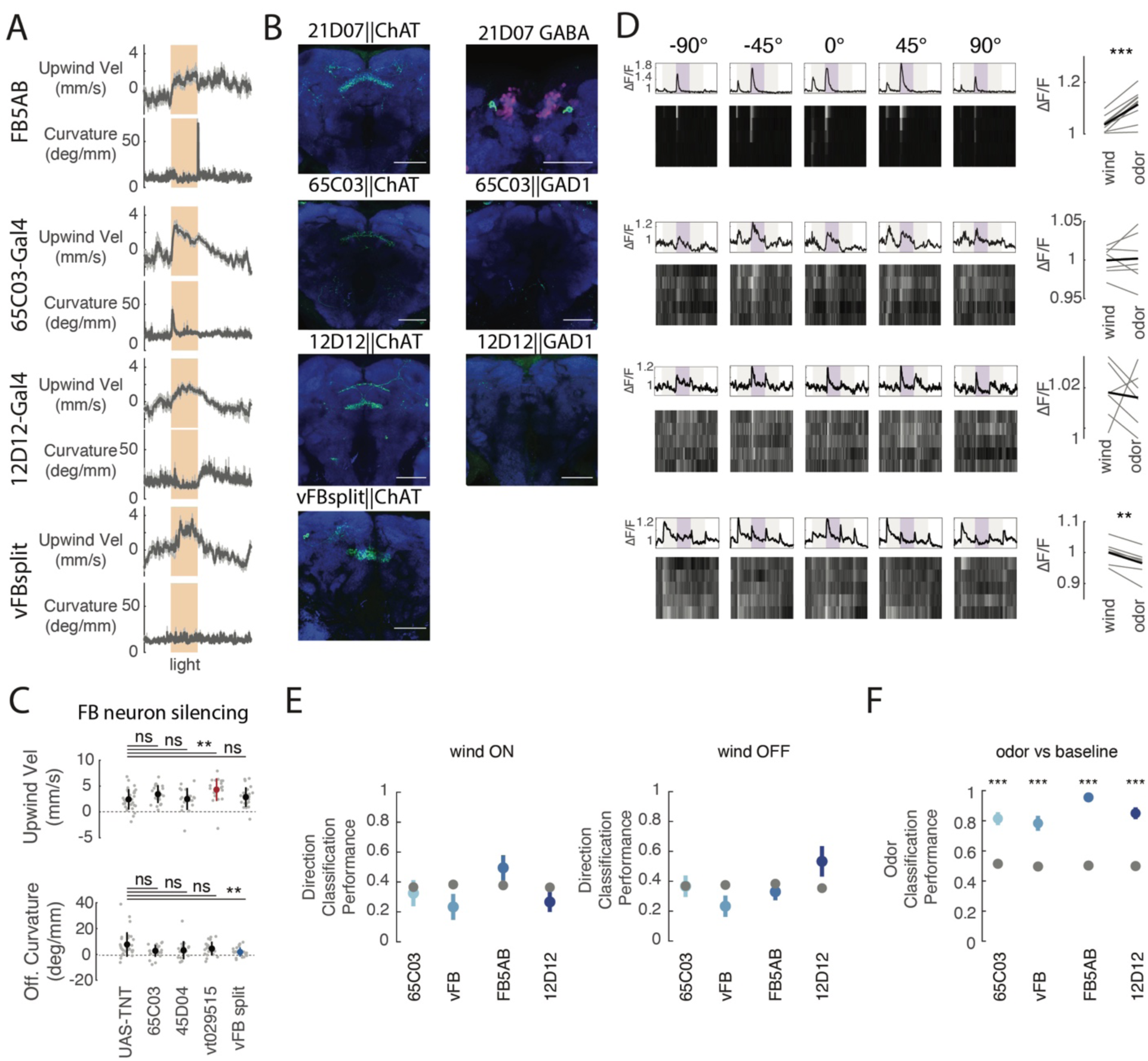
Additional data on odor-responsive, upwind-promoting FB tangential neurons. **A)** Timecourses of upwind velocity and curvature averaged across flies for all FB tangential input lines that showed significant responses to vinegar (mean±SEM), 21D07||CLIN (N=27), 65C03 (N=24), 12D12 (N=38), vFB split (N=27)). **B)** Odor-responsive FB tangential inputs are cholinergic and not GABAergic. Left column: Flies expressing *chrimson-mVenus* under the control of each driver using a ChAT-LexA,LexAOP-FLP strategy (green, see Materials and Methods). Right column: 21D07-GAL4: same genotype as left, but co-stained with anti-GABA (magenta). There is no colocalization of GABA and Chrimson-mVenus. Flies expressing *Chrimson-mVenus* using a Gad1-LexA, LexAOP-FLP strategy (green) do not label FB neurons in 65C03-GAL4 or 12D12-GAL4. Scale bars represent 50µM. **C)** Top: Effects of silencing FB neurons on behavioral responses to odor (10s 1% vinegar). Mean±STD overlaid. (Mann Whitney U test comparing to UAS-TNT control: upwind: 65C03: p=0.048027, 45D04: p=0.65028, vt029515: p=0.00086718, vFB split p=0.60472, OFF curvature: 65C03: p=0.044317, 45D04: p=0.087747, vt029515: p=0.31643, vFB split: p=0.0021892). **D)** Left: Example calcium responses from FB tangential inputs in individual flies. Top row depicts average ΔF/F for each direction across 5 trials, heat maps below depict responses in individual trials. Right: Calcium response to wind alone vs vinegar. Gray lines represent individual flies and black lines represent genotype means. Significant increase for 21D07-GAL4 (paired student t-test: p=3.9090e-04). No significant increase for 65C03 or 12D12 (paired student’s t-test, p=0.7790, p=0.7782) and significant decreases for vFB split (paired student t-test; p = 0.0028) **E)** Performance of tree classifiers at decoding wind direction (left, center, right). Left: classifier trained on the first 5s of wind ON. Student’s t-test: 65C03 p=0.6504, vFB p=0.1088, FB5AB p=0.1859, 12D12 p=0.1650. Right: classifier trained on the 5s following wind OFF. Student’s t-test 65C03 p=0.9802, vFB p=0.0646, FB5AB p=0.3656, 12D12 p=0.0995. Due to stimulus presentation differences between preps, PFNa data here is shown relative to wind ON as a control. Gray dots represent classifiers trained with the same data and shuffled labels. **F)** Performance of a tree classifier at decoding odor versus baseline activity, trained on 5s of baseline versus first 5s of odor ON. Gray dots represent a classifier trained with the same data and shuffled labels. Student’s t-test: 65C03 p=1.6787e-06, vFB p=2.0903e-05, FB5AB p=5.7008e-13, 12D12 p=5.8079e-08. All error bars on classifiers represent SEM and all comparisons for behavioral experiments and classifier performance are Bonferroni corrected for multiple comparisons.

**Fig. S5:**
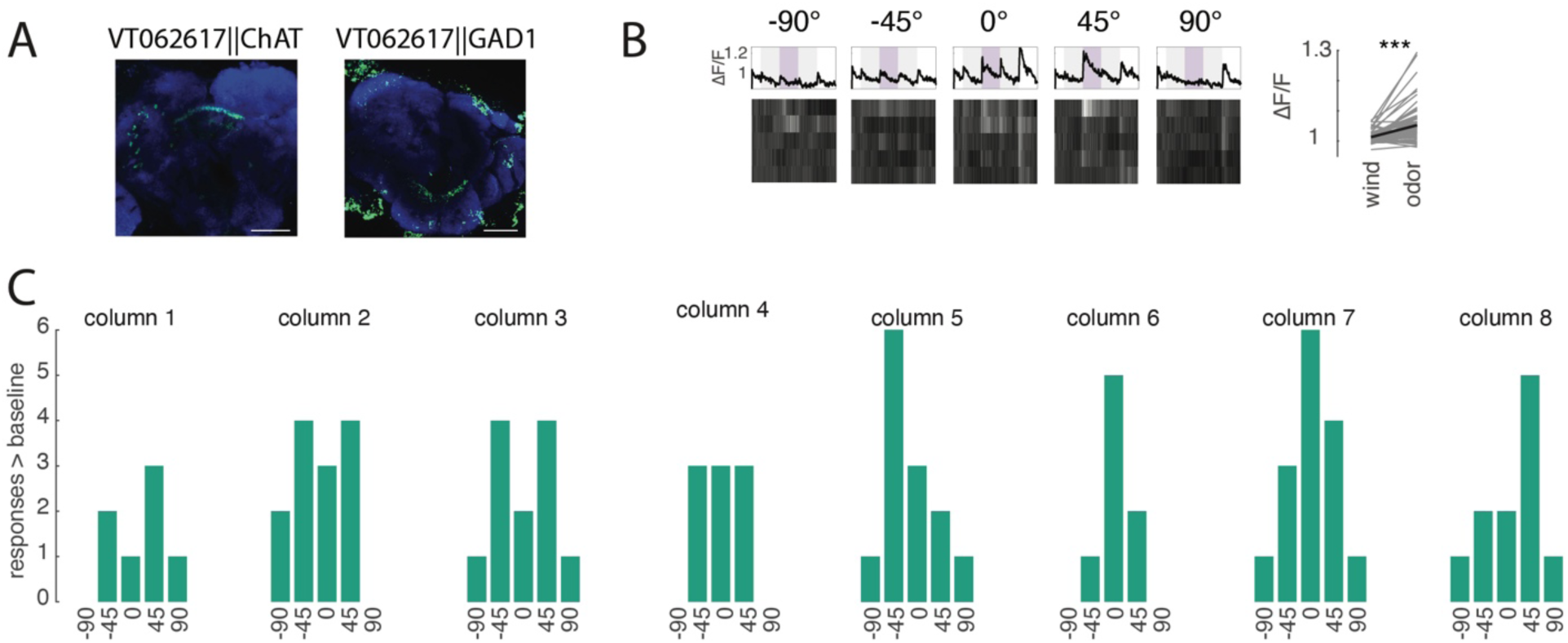
Additional data on hΔC responses. **A)** hΔC neurons are cholinergic. Left column: Flies expressing *chrimson-mVenus* under the control of VT062617-GAL4 using a ChAT-LexA,LexAOP-FLP strategy (green, see Materials and Methods). Right column: Flies expressing *Chrimson-mVenus* using a Gad1-LexA, LexAOP-FLP strategy (green) do not label FB neurons in VT062617-GAL4. Scale bars represent 50µM. **B)** Calcium response examples from an individual fly for hΔC (left). Top row depicts average ΔF/F for each direction across 5 trials, heat maps below depict responses in individual trials. Right: Calcium response to wind alone (5s after wind ON) vs vinegar (5s after odor ON). Gray lines represent average increases of individual active columns. Odor response is significantly larger than wind response (paired Student t-test: 4.7885e-11). **C)** Directional responses are not correlated with anatomical column. Histograms depict counts of directional responses during the odor period (first 5s) for each column >2STD above baseline period. Columns 1-8 correspond to fly’s right to left.

**Fig. S6:**
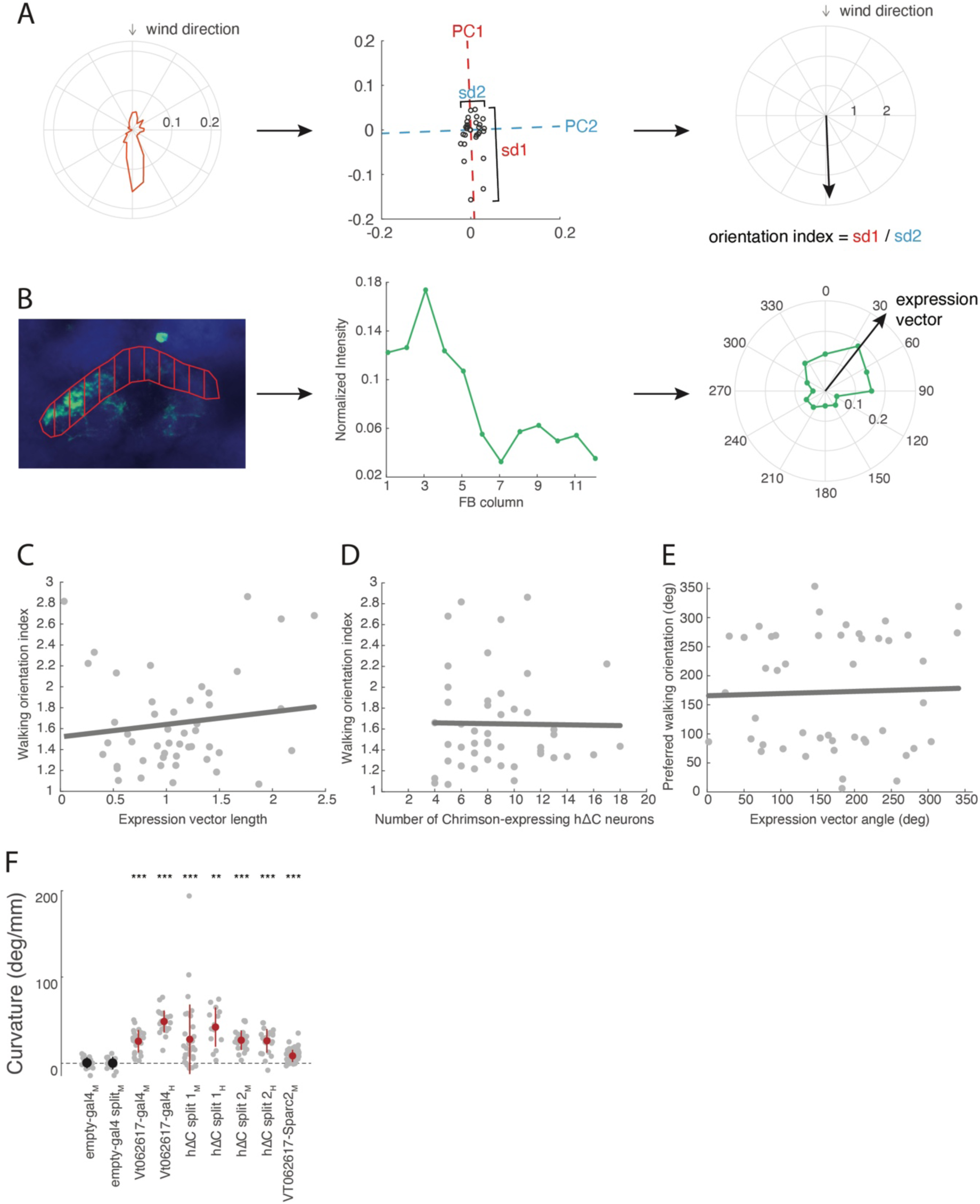
Additional data on hΔC activation. **A)** Strategy for calculating the walking orientation index and preferred walking direction of SPARC flies. Orientation data from 2-6 s after light ON were converted into an orientation histogram (left). We performed PCA on this histogram in Cartesian coordinates (center), then computed the orientation index as the ratio of the standard deviation along PC1 to the standard deviation along PC2 (right). We computed the preferred walking direction as the direction of PC1. **B)** Strategy for calculating the expression vector of hΔC > SPARC flies. We divided the output tufts of hΔC neurons into 12 columns (left), and computed normalized fluorescence across these columns (middle). We then converted the normalized expression profile into polar coordinates and summed to create an expression vector (right). **C)** The orientation index of hΔC > SPARC flies is not related to the expression vector length (correlation: *p* = 0.3607, *r^2^* = 0.1349). **D)** The orientation index of hΔC > SPARC flies is not related to the total number of Chrimson-expressing hΔC neurons (correlation: *p* = 0.9257, *r^2^* = 0.0002). **E)** The preferred direction of hΔC > SPARC flies is not related to the expression vector angle (correlation: *p* = 0.8263, *r^2^* = 0.0011). **F)** Curvature during optogenetic activation (mean ± STD) of empty GAL4 and various hΔC lines with medium (M: 26 µW/mm^2^) or high (H: 34 µW/mm^2^) light power. Empty-Gal4: p=0.5732, empty split p=0.7869, VT062617-Gal4 (medium) p= 1.2290e-05, VT062617-Gal4 (high) p=1.2290e-05, hΔC split1 (medium) p=1.8517e-05, hΔC split1 (high) p=1.2207e-04, hΔC split2 (medium) p=5.6061e-06, hΔc split2 (high) p=2.3518e-05, VT062617-SPARC2 (medium) p=2.2783e-12.

**Fig. S7:**
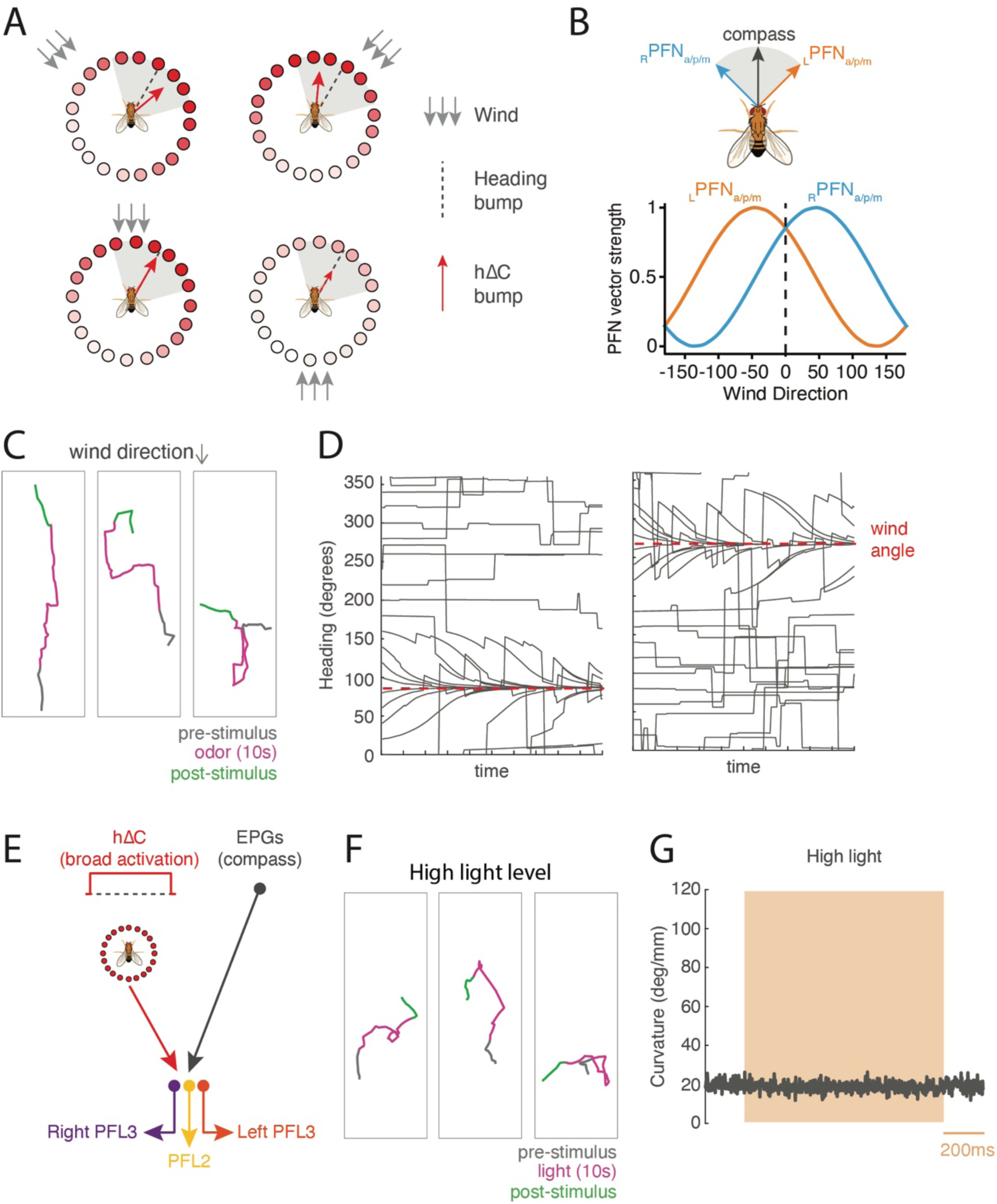
Additional model simulations. **A)** Alternate model wind representation in hΔC neurons, that can be built from two frontally tuned PFN populations as described in Currier et al. 2020. Wind direction is represented as a bump of activity (circles) and as a vector (red), while heading direction is represented as a vector (gray). As in the allocentric representation (Fig. 7B), the wind vector is to the right of the heading vector for leftward wind and to the left of the heading vector for rightward wind. However, in this scheme, wind vectors are contained within a 90° window centered around the heading vector, and wind from the rear produces a bump of reduced amplitude. **B)** Hypothesized wind representations wind-tuned PFNs (PFNa/p/m) based on physiology from Currier et al. 2020 and anatomy from Hulse et al. 2021. PFNs receive a shifted heading bump from the compass such that left and right hemisphere PFNs exhibit bumps offset by 45° for left PFNs and -45° for right PFNs (top). PFNs in each hemisphere are maximally activated by wind arriving from 45° ipsilateral (−45° for left PFNs, +45° for right PFNs) such that the length of the vectors increase or decrease across wind directions (bottom). **C)** Behavioral trajectories of simulated flies using the frontal wind representation in **A** before, during, and after odor. Note that flies go upwind so long as they are initially orientated within +/- 90° of upwind. **D)** Heading of simulated flies using the frontal wind representation in **A** for two different wind directions and several different initial headings. Note that flies go upwind so long as they are initially orientated within +/- 90° of upwind, or reach one of these orientations through random turns. **E)** Simulated circuit activity during broad optogenetic activation without mutual inhibition. In this simulation, every hΔC neuron is activated equally for the duration of the stimulus but the mutual inhibition layer is omitted. **F)** Behavioral trajectories of simulated flies lacking the mutual inhibition circuit during broad hΔC activation. Random turn rate is the same before, during, and after stimulation. **G)** Average curvature simulated flies lacking the mutual inhibition circuit during broad optogenetic activation using high light. Without the mutual inhibition circuit, broad activation of hΔC neurons does not drive an increase in curvature.

## Notes

### Competing Interest Statement

The authors have declared no competing interest.

## References

Álvarez-Salvado, E., Licata, A.M., Connor, E.G., McHugh, M.K., King, B.M., Stavropoulos, N., Victor, J.D., Crimaldi, J.P. and Nagel, K.I., 2018. Elementary sensory-motor transformations underlying olfactory navigation in walking fruit-flies. Elife, 7, p.e37815.

Ardin, P., Peng, F., Mangan, M., Lagogiannis, K. and Webb, B., 2016. Using an insect mushroom body circuit to encode route memory in complex natural environments. PLoS computational biology, 12(2), p.e1004683.

Aso, Y., Sitaraman, D., Ichinose, T., Kaun, K.R., Vogt, K., Belliart-Guérin, G., Plaçais, P.Y., Robie, A.A., Yamagata, N., Schnaitmann, C. and Rowell, W.J., 2014. Mushroom body output neurons encode valence and guide memory-based action selection in Drosophila. Elife, 3, p.e04580.

Bates, A.S., Schlegel, P., Roberts, R.J., Drummond, N., Tamimi, I.F., Turnbull, R., Zhao, X., Marin, E.C., Popovici, P.D., Dhawan, S. and Jamasb, A., 2020. Complete connectomic reconstruction of olfactory projection neurons in the fly brain. Current Biology, 30(16), pp.3183–3199.

Bell, W.J. and Kramer, E., 1979. Search and anemotactic orientation of cockroaches. Journal of Insect Physiology, 25(8), pp.631–640.

Buehlmann, C., Hansson, B.S. and Knaden, M., 2012. Path integration controls nest-plume following in desert ants. Current Biology, 22(7), pp.645–649.

Cardé, R.T. and Willis, M.A., 2008. Navigational strategies used by insects to find distant, wind-borne sources of odor. Journal of chemical ecology, 34(7), pp.854–866.

Celani, A., Villermaux, E. and Vergassola, M., 2014. Odor landscapes in turbulent environments. Physical Review X, 4(4), p.041015.

Chaffiol, A., Laloi, D. and Pham-Delègue, M.H., 2005. Prior classical olfactory conditioning improves odour-cued flight orientation of honey bees in a wind tunnel. Journal of experimental biology, 208(19), pp.3731–3737.

Clyne, J.D. and Miesenböck, G., 2008. Sex-specific control and tuning of the pattern generator for courtship song in Drosophila. Cell, 133(2), pp.354–363.

Cognigni, P., Felsenberg, J. and Waddell, S., 2018. Do the right thing: neural network mechanisms of memory formation, expression and update in Drosophila. Current opinion in neurobiology, 49, pp.51–58.

Crimaldi, J.P. and Koseff, J.R., 2001. High-resolution measurements of the spatial and temporal scalar structure of a turbulent plume. Experiments in Fluids, 31(1), pp.90–102.

Currier, T.A., Matheson, A.M. and Nagel, K.I., 2020. Encoding and control of orientation to airflow by a set of Drosophila fan-shaped body neurons. Elife, 9, p.e61510.

Dacke, M., Bell, A.T., Foster, J.J., Baird, E.J., Strube-Bloss, M.F., Byrne, M.J. and El Jundi, B., 2019. Multimodal cue integration in the dung beetle compass. Proceedings of the National Academy of Sciences, 116(28), pp.14248–14253.

David, C.T., Kennedy, J.S. and Ludlow, A.R., 1983. Finding of a sex pheromone source by gypsy moths released in the field. Nature, 303(5920), pp.804–806.

De Belle, J.S. and Heisenberg, M., 1994. Associative odor learning in Drosophila abolished by chemical ablation of mushroom bodies. Science, 263(5147), pp.692–695.

Dolan, M.J., Frechter, S., Bates, A.S., Dan, C., Huoviala, P., Roberts, R.J., Schlegel, P., Dhawan, S., Tabano, R., Dionne, H. and Christoforou, C., 2019. Neurogenetic dissection of the Drosophila lateral horn reveals major outputs, diverse behavioural functions, and interactions with the mushroom body. Elife, 8, p.e43079.

Donlea, J.M., Pimentel, D. and Miesenböck, G., 2014. Neuronal machinery of sleep homeostasis in Drosophila. Neuron, 81(4), pp.860–872.

Fuller, S.B., Straw, A.D., Peek, M.Y., Murray, R.M. and Dickinson, M.H., 2014. Flying Drosophila stabilize their vision-based velocity controller by sensing wind with their antennae. Proceedings of the National Academy of Sciences, 111(13), pp.E1182–E1191.

Goulard, R., Buehlmann, C., Niven, J.E., Graham, P. and Webb, B., 2021. A unified mechanism for innate and learned visual landmark guidance in the insect central complex. PLoS computational biology, 17(9), p.e1009383.

Grünbaum, D. and Willis, M.A., 2015. Spatial memory-based behaviors for locating sources of odor plumes. Movement ecology, 3(1), pp.1–21.

Hargreaves, E.L., Rao, G., Lee, I. and Knierim, J.J., 2005. Major dissociation between medial and lateral entorhinal input to dorsal hippocampus. science, 308(5729), pp.1792–1794.

Hige, T., Aso, Y., Rubin, G.M. and Turner, G.C., 2015. Plasticity-driven individualization of olfactory coding in mushroom body output neurons. Nature, 526(7572), pp.258–262.

Hige, T., Aso, Y., Modi, M.N., Rubin, G.M. and Turner, G.C., 2015. Heterosynaptic plasticity underlies aversive olfactory learning in Drosophila. Neuron, 88(5), pp.985–998.

Honkanen, A., Adden, A., da Silva Freitas, J. and Heinze, S., 2019. The insect central complex and the neural basis of navigational strategies. Journal of Experimental Biology, 222(Suppl_1), p.jeb188854.

Homberg, U., 1985. Interneurones of the central complex in the bee brain (Apis mellifera, L.). Journal of insect physiology, 31(3), pp.251–264.

Hulse, B.K., Haberkern, H., Franconville, R., Turner-Evans, D.B., Takemura, S., Wolff, T., Noorman, M., Dreher, M., Dan, C., Parekh, R. and Hermundstad, A.M., 2021. A connectome of the Drosophila central complex reveals network motifs suitable for flexible navigation and context-dependent action selection. Elife, 10:e66039.

Huoviala, P., Dolan, M.J., Love, F.M., Myers, P., Frechter, S., Namiki, S., Pettersson, L., Roberts, R.J., Turnbull, R., Mitrevica, Z. and Breads, P., 2020. Neural circuit basis of aversive odour processing in Drosophila from sensory input to descending output. BioRxiv, p.394403.

Jung, S.H., Hueston, C. and Bhandawat, V., 2015. Odor-identity dependent motor programs underlie behavioral responses to odors. Elife, 4, p.e11092.

Kennedy, J.S. and Marsh, D., 1974. Pheromone-regulated anemotaxis in flying moths. Science, 184(4140), pp.999–1001.

Larsson, M.C., Domingos, A.I., Jones, W.D., Chiappe, M.E., Amrein, H. and Vosshall, L.B., 2004. Or83b encodes a broadly expressed odorant receptor essential for Drosophila olfaction. Neuron, 43(5), pp.703–714.

Leitner, F.C., Melzer, S., Lütcke, H., Pinna, R., Seeburg, P.H., Helmchen, F. and Monyer, H., 2016. Spatially segregated feedforward and feedback neurons support differential odor processing in the lateral entorhinal cortex. Nature neuroscience, 19(7), pp.935–944.

Le Moël, F., Stone, T., Lihoreau, M., Wystrach, A. and Webb, B., 2019. The central complex as a potential substrate for vector based navigation. Frontiers in psychology, 10, p.690.

Li, F., Lindsey, J.W., Marin, E.C., Otto, N., Dreher, M., Dempsey, G., Stark, I., Bates, A.S., Pleijzier, M.W., Schlegel, P. and Nern, A., 2020. The connectome of the adult Drosophila mushroom body provides insights into function. Elife, 9, p.e62576.

Lu, J., Westeinde, E.A., Hamburg, L., Dawson, P.M., Lyu, C., Maimon, G., Druckmann, S. and Wilson, R.I., 2020. Transforming representations of movement from body-to world-centric space. bioRxiv.

Lyu, C., Abbott, L.F. and Maimon, G., 2020. A neuronal circuit for vector computation builds an allocentric traveling-direction signal in the Drosophila fan-shaped body. bioRxiv.

Mamiya, A., Beshel, J., Xu, C. and Zhong, Y., 2008. Neural representations of airflow in Drosophila mushroom body. PLoS One, 3(12), p.e4063.

Martin, J.P., Guo, P., Mu, L., Harley, C.M. and Ritzmann, R.E., 2015. Central-complex control of movement in the freely walking cockroach. Current Biology, 25(21), pp.2795–2803.

Masson, J.B., 2013. Olfactory searches with limited space perception. Proceedings of the National Academy of Sciences, 110(28), pp.11261–11266.

Mishkin, M., Ungerleider, L.G. and Macko, K.A., 1983. Object vision and spatial vision: two cortical pathways. Trends in neurosciences, 6, pp.414–417.

Mohamed, G.A., Cheng, R.K., Ho, J., Krishnan, S., Mohammad, F., Claridge-Chang, A. and Jesuthasan, S., 2017. Optical inhibition of larval zebrafish behaviour with anion channelrhodopsins. BMC biology, 15(1), pp.1–12.

Murlis, J., Elkinton, J.S. and Carde, R.T., 1992. Odor plumes and how insects use them. Annual review of entomology, 37(1), pp.505–532.

Namiki, S. and Kanzaki, R., 2016. Comparative neuroanatomy of the lateral accessory lobe in the insect brain. Frontiers in Physiology, 7, p.244.

Okubo, T.S., Patella, P., D’Alessandro, I. and Wilson, R.I., 2020. A neural network for wind-guided compass navigation. Neuron, 107(5), pp.924–940.

Owald, D., Felsenberg, J., Talbot, C.B., Das, G., Perisse, E., Huetteroth, W. and Waddell, S., 2015. Activity of defined mushroom body output neurons underlies learned olfactory behavior in Drosophila. Neuron, 86(2), pp.417–427.

Page, J.L., Dickman, B.D., Webster, D.R. and Weissburg, M.J., 2011. Getting ahead: context-dependent responses to odorant filaments drive along-stream progress during odor tracking in blue crabs. Journal of Experimental Biology, 214(9), pp.1498–1512.

Rayshubskiy, A., Holtz, S.L. , D’Alessandro, I. , Li, A.A., Vanderbeck, Q.X., Haber, I.S., Gibb, P.W., and Wilson, R.I 2020. Neural circuit mechanisms for steering control in walking *Drosophila* bioRxiv 2020.04.04.024703;

Ren, Q., Awasaki, T., Huang, Y.F., Liu, Z. and Lee, T., 2016. Cell class-lineage analysis reveals sexually dimorphic lineage compositions in the Drosophila brain. Current Biology, 26(19), pp.2583–2593.

Sareen, P.F., McCurdy, L.Y. and Nitabach, M.N., 2021. A neuronal ensemble encoding adaptive choice during sensory conflict in Drosophila. Nature communications, 12(1), pp.1–13.

Sayin, S., De Backer, J.F., Siju, K.P., Wosniack, M.E., Lewis, L.P., Frisch, L.M., Gansen, B., Schlegel, P., Edmondson-Stait, A., Sharifi, N. and Fisher, C.B., 2019. A neural circuit arbitrates between persistence and withdrawal in hungry Drosophila. Neuron, 104(3), pp.544–558.

Scaplen, K.M., Talay, M., Fisher, J.D., Cohn, R., Sorkaç, A., Aso, Y., Barnea, G. and Kaun, K.R., 2021. Transsynaptic mapping of Drosophila mushroom body output neurons. Elife, 10, p.e63379.

Scheffer, L.K., Xu, C.S., Januszewski, M., Lu, Z., Takemura, S.Y., Hayworth, K.J., Huang, G.B., Shinomiya, K., Maitlin-Shepard, J., Berg, S. and Clements, J., 2020. A connectome and analysis of the adult Drosophila central brain. Elife, 9, p.e57443.

Schlegel, P., Bates, A.S., Stürner, T., Jagannathan, S.R., Drummond, N., Hsu, J., Capdevila, L.S., Javier, A., Marin, E.C., Barth-Maron, A. and Tamimi, I.F., 2021. Information flow, cell types and stereotypy in a full olfactory connectome. Elife, 10, p.e66018.

Schulze, A., Gomez-Marin, A., Rajendran, V.G., Lott, G., Musy, M., Ahammad, P., Deogade, A., Sharpe, J., Riedl, J., Jarriault, D. and Trautman, E.T., 2015. Dynamical feature extraction at the sensory periphery guides chemotaxis. Elife, 4, p.e06694.

Seelig, J.D. and Jayaraman, V., 2015. Neural dynamics for landmark orientation and angular path integration. Nature, 521(7551), pp.186–191.

Shpiro, A., Curtu, R., Rinzel, J. and Rubin, N., 2007. Dynamical characteristics common to neuronal competition models. Journal of neurophysiology, 97(1), pp.462–473.

Silbering, A.F., Rytz, R., Grosjean, Y., Abuin, L., Ramdya, P., Jefferis, G.S. and Benton, R., 2011. Complementary function and integrated wiring of the evolutionarily distinct Drosophila olfactory subsystems. Journal of Neuroscience, 31(38), pp.13357–13375.

Stone, T., Webb, B., Adden, A., Weddig, N.B., Honkanen, A., Templin, R., Wcislo, W., Scimeca, L., Warrant, E. and Heinze, S., 2017. An anatomically constrained model for path integration in the bee brain. Current Biology, 27(20), pp.3069–3085.

Strauss, R. and Heisenberg, M., 1993. A higher control center of locomotor behavior in the Drosophila brain. Journal of Neuroscience, 13(5), pp.1852–1861.

Sun, X., Yue, S. and Mangan, M., 2020. A decentralised neural model explaining optimal integration of navigational strategies in insects. Elife, 9, p.e54026.

Sun, X., Yue, S. and Mangan, M., 2021. How the insect central complex could coordinate multimodal navigation. bioRxiv.

Suver, M.P., Matheson, A.M., Sarkar, S., Damiata, M., Schoppik, D. and Nagel, K.I., 2019. Encoding of wind direction by central neurons in Drosophila. Neuron, 102(4), pp.828–842.

Talay, M., Richman, E.B., Snell, N.J., Hartmann, G.G., Fisher, J.D., Sorkaç, A., Santoyo, J.F., Chou-Freed, C., Nair, N., Johnson, M. and Szymanski, J.R., 2017. Transsynaptic mapping of second-order taste neurons in flies by trans-Tango. Neuron, 96(4), pp.783–795.

van Breugel, F. and Dickinson, M.H., 2014. Plume-tracking behavior of flying Drosophila emerges from a set of distinct sensory-motor reflexes. Current Biology, 24(3), pp.274–286.

Vergassola, M., Villermaux, E. and Shraiman, B.I., 2007. ‘Infotaxis’ as a strategy for searching without gradients. Nature, 445(7126), pp.406–409.

Webster, D.R. and Weissburg, M.J., 2001. Chemosensory guidance cues in a turbulent chemical odor plume. Limnology and Oceanography, 46(5), pp.1034–1047.

Wolf, H. and Wehner, R., 2000. Pinpointing food sources: olfactory and anemotactic orientation in desert ants, Cataglyphis fortis. Journal of Experimental Biology, 203(5), pp.857–868.

## Methods References

Giovannucci A, Friedrich J, Gunn P, Kalfon J, Brown BL, Koay SA, Taxidis J, Najafi F, Gauthier JL, Zhou P, Khakh BS, Tank DW, Chklovskii DB, Pnevmatikakis EA. 2019. CaImAn an open source tool for scalable calcium imaging data analysis. eLife 8:e38173. DOI: https://doi.org/10.7554/eLife.38173, PMID: 30652683

Muir DR, Kampa BM. 2014. FocusStack and StimServer: a new open source MATLAB toolchain for visual stimulation and analysis of two-photon calcium neuronal imaging data. Frontiers in Neuroinformatics 8:85. DOI: https://doi.org/10.3389/fninf.2014.00085, PMID: 25653614

Nagel KI, Wilson RI. 2016. Mechanisms underlying population response dynamics in inhibitory interneurons of the Drosophila Antennal Lobe. The Journal of Neuroscience 36:4325–4338. DOI: https://doi.org/10.1523/JNEUROSCI.3887-15.2016, PMID: 27076428

Pnevmatikakis EA, Giovannucci A. 2017. NoRMCorre: an online algorithm for piecewise rigid motion correction of calcium imaging data. Journal of Neuroscience Methods 291:83–94. DOI: https://doi.org/10.1016/j.jneumeth.2017.07.031, PMID: 28782629

